# Heterogeneous non-canonical nucleosomes predominate in yeast cells *in situ*

**DOI:** 10.1101/2021.04.04.438362

**Authors:** Zhi Yang Tan, Shujun Cai, Alex J. Noble, Jon K. Chen, Jian Shi, Lu Gan

## Abstract

Nuclear processes depend on the organization of chromatin, whose basic units are cylinder-shaped complexes called nucleosomes. A subset of mammalian nucleosomes *in situ* (inside cells) resembles the canonical structure determined *in vitro* 25 years ago. Nucleosome structure *in situ* is otherwise poorly understood. Using cryo-ET and 3-D classification analysis of budding yeast cells, here we find that canonical nucleosomes account for less than 10% of total nucleosomes expected *in situ*. In a strain in which H2A-GFP is the sole source of histone H2A, class averages that resemble canonical nucleosomes both with and without GFP densities are found *ex vivo* (in nuclear lysates), but not *in situ*. These data suggest that the budding yeast intranuclear environment favors multiple non-canonical nucleosome conformations. Using the structural observations here and the results of previous genomics and biochemical studies, we propose a model in which the average budding yeast nucleosome’s DNA is partially detached *in situ*.

## INTRODUCTION

Eukaryotic chromosomes are polymers of DNA-protein complexes called nucleosomes. An octamer of proteins, consisting of a heterotetramer of histones H3 and H4 and two heterodimers of histones H2A and H2B, resides at the nucleosome’s center (Luger et al., 1997). Canonical nucleosomes resemble 10 nm wide, 6 nm thick cylinders and have 145 – 147 base pairs of DNA bent in 1.65 left-handed superhelical gyres around the histone octamer (Zhou et al., 2019; Zlatanova et al., 2009). In contrast, non-canonical nucleosomes have either partially detached DNA, partially detached histones, fewer than 8 histones, or a combination of these features (Zlatanova et al., 2009). Both X-ray crystallography and single-particle cryo-EM have shown that reconstituted nucleosomes, either alone or within a complex, are largely canonical *in vitro* (Zhou et al., 2019). Nucleosome structures *in situ* inside cells remain mysterious.

In the context of chromatin, nucleosomes are not discrete particles because sequential nucleosomes are connected by short stretches of linker DNA. Variation in linker DNA structure is a source of chromatin conformational heterogeneity (Collepardo-Guevara and Schlick, 2014). Recent cryo-EM studies show that nucleosomes can deviate from the canonical form *in vitro*, primarily in the structure of DNA near the entry/exit site (Bilokapic et al., 2018; Fukushima et al., 2022; Sato et al., 2021; Zhou et al., 2021). In addition to DNA structural variability, nucleosomes *in vitro* have small changes in histone conformations (Bilokapic et al., 2018). Larger-scale variations of DNA and histone structure are not compatible with high-resolution analysis and may have been missed in single-particle cryo-EM studies.

Molecular-resolution (2 – 4 nm) studies of unique objects like cells may be obtained by cryo-electron tomography (cryo-ET), a form of cryo-electron microscopy (cryo-EM) that generates 3-D reconstructions called cryotomograms. These studies reveal life-like snapshots of macromolecular complexes because the samples are prepared and then imaged in an unfixed, unstained, frozen-hydrated state. Most eukaryotic cells are too thick for cryo-ET, so thinner frozen-hydrated samples are made by cutting by cryomicrotomy or thinning by cryo-focused ion beam (cryo-FIB) milling (Ng and Gan, 2020; Strunk et al., 2012). These two approaches respectively produce cryosections and plank-like samples called cryolamellae. Subvolumes called subtomograms contain independent copies of the cells’ macromolecular complexes. These subtomograms can be further studied by averaging, which increases the signal-to-noise ratio, and classification, which facilitates the analysis of heterogeneity. Large macromolecular complexes such as ribosomes and proteasomes have been identified *in situ* by 3-D classification followed by comparison of the class averages to known structures – an approach called purification *in silico* (Beck and Baumeister, 2016). Using the purification *in silico* approach, we previously showed that canonical nucleosomes exist in cryotomograms of yeast cell lysates *ex vivo* and in a HeLa cell cryolamella (Cai et al., 2018a; Cai et al., 2018b; Cai et al., 2018c). Herein, we use the term *ex vivo* to describe nucleosomes from lysates instead of the term *in vitro*, which is more commonly used to describe either reconstituted or purified mononucleosomes. However, our 3-D structural analysis did not generate canonical nucleosome structures from cryosectioned fission yeast cells (Cai et al., 2018b). The discrepancy between nucleosome class averages *ex vivo* and *in situ* could have either technical or biological origins. As a biological explanation for the absence of canonical nucleosome class averages *in situ*, we hypothesized that yeast nucleosomes are either conformationally or constitutionally heterogeneous.

In this work, we test this heterogenous-nucleosome hypothesis by using cryo-ET to image both wild-type cells and strains that have nucleosomes bearing GFP as a density tag. We use the budding yeast *S. cerevisiae*, herein called yeast, because it has only two copies of each histone gene and because it is more amenable to gene editing. Our work compares the chromatin in both lysates and thin cellular cryo-EM samples prepared primarily by cryo-FIB milling. To obtain more information about nucleosomes *in situ*, we create one strain in which H2A-GFP is the sole source of H2A, meaning that the nucleosomes are expected to project one or two extra densities from their surface. Canonical nucleosomes are abundant in nuclear lysates and, in the H2A-GFP-expressing strain’s nuclear lysates, have extra densities consistent with GFP. In contrast, canonical nucleosomes account for less than 10% of the expected number of total nucleosomes in wild-type cell cryolamellae. Furthermore, neither canonical nucleosomes nor nucleosome-like particles with extra protruding densities were detected in the H2A-GFP-expressing strain. These findings suggest that the yeast intracellular environment disfavors the canonical nucleosome conformation.

## RESULTS

### Canonical nucleosomes are abundant in wild-type yeast lysates

The crystal structure of the reconstituted yeast nucleosome (White et al., 2001) shows a canonical structure that is largely indistinguishable from the first published one (Luger et al., 1997). To describe the various views of the nucleosome, herein we use the compact nomenclature introduced by (Zhou et al., 2019): the disc view is along the superhelical axis, and the gyre view is along the pseudo-dyad axis (Figure 1A). The side view, which was not defined by Zhou et al, is orthogonal to both the disc and gyre views.

**Figure 1.**
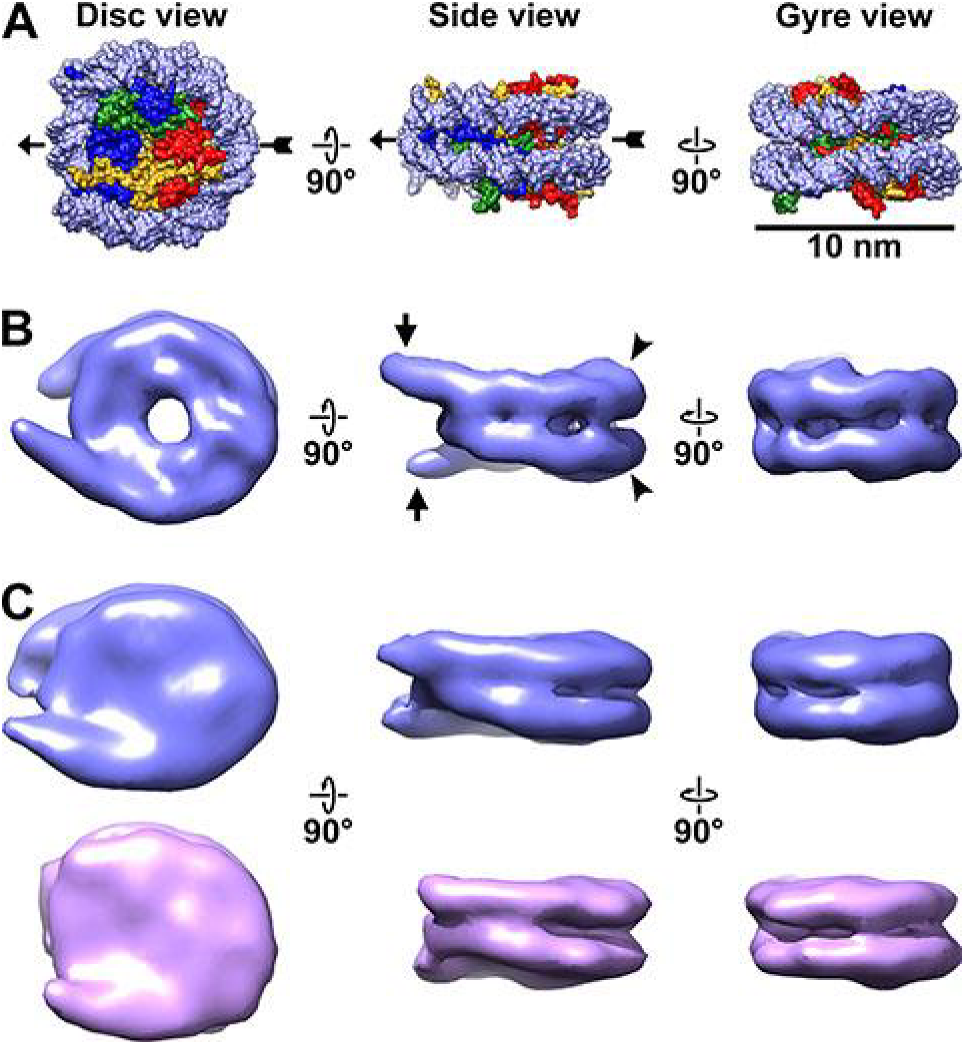
Canonical nucleosomes *in vitro* and *in situ*. (A) Space-filling model of the reconstituted yeast nucleosome crystal structure (PDB 1ID3) (White et al., 2001), showing from left to right, the disc, side, and gyre views. The pseudo-dyad axis is indicated by the arrow. The DNA is rendered as light blue and the histones in the core are shaded blue (H3), green (H4), red (H2B), and yellow (H2A). (B) Subtomogram average of nucleosomes from wild-type (BY4741) yeast nuclear lysates. The linker DNA is indicated by the short arrows and the DNA gyre motifs are indicated by the arrowheads. (C) Subtomogram averages of nucleosomes in wild-type cell cryolamellae, oriented similarly to the nucleosomes in the other two panels. The upper (blue) class has more ordered linker DNA than the lower (magenta) class. Note that the subtomogram average in panel B looks different from those in panel C (and in Figures 3 and S17) because it is at higher resolution (18 Å vs 24 Å). The gap in the disc view of the nuclear-lysate-based average is due to the lower concentration of amino acids there, which is not visible in panel A due to space-filling rendering. This gap’s visibility may also depend on the contrast mechanism because it is not visible in the VPP averages.

The original RELION subtomogram-analysis workflow (Bharat and Scheres, 2016) involves 2-D classification, followed by 3-D classification (Figure S1A), whereas in the alternative approach, 2-D classification is bypassed and subtomograms are subjected directly to 3-D classification (Figure S1B); for brevity, we term this alternative method “direct 3-D classification”. Direct 3-D classification is limited by computer hardware (Kimanius et al., 2016), but can detect more canonical nucleosomes (Cai et al., 2018a); see the Methods for more details. In this study, the original workflow is used on a subset of samples to show example 2-D class averages for comparison with other studies. However, the conclusions in this paper are drawn from direct 3-D classification.

Our previous subtomogram analysis of nuclear lysates (Cai et al., 2018c) revealed that nucleosomes from the wild-type strain YEF473A (Bi and Pringle, 1996) adopt the canonical structure *ex vivo*. We repeated this experiment on the strain BY4741 (Brachmann et al., 1998), which serves as the wild-type and parent strain for the histone-GFP tagging mutants described later. In this experiment, yeast nuclei are isolated, lysed, then deposited on an EM grid. Cryotomograms of BY4741 nuclear lysates reveal the crowded nucleosome-like particles and other cellular debris (Figure S2). Next, we template matched for nucleosome-like particles using a nucleosome-sized featureless cylinder as a reference. Direct 3-D classification (Figure S3, A and B) followed by 3-D refinement produced an 18 Å-resolution subtomogram average of a BY4741 canonical nucleosome class (Figure S3C). The subtomogram average of wild-type yeast nucleosomes in nuclear lysates has longer linker DNA (Figure 1B) than the crystal structure, which was reconstituted using a 146-bp DNA fragment. Because the nucleosome-repeat length of budding yeast chromatin is ∼168 bp (Brogaard et al., 2012), this extra length of DNA may come from an ordered portion of the ∼22 bp linker between adjacent nucleosomes.

### Canonical nucleosomes are rare in wild-type cells *in situ*

Cryo-FIB milling is a compression-free method to thin frozen-hydrated cells (Hayles et al., 2007; Mahamid et al., 2015; Marko et al., 2006; Medeiros et al., 2018; Rigort et al., 2010; Villa et al., 2013). This technique uses a beam of gallium ions to thin a cell under cryogenic conditions, producing a frozen-hydrated plank-like cellular sample called a cryolamella. Canonical nucleosomes are detectable in a HeLa cell cryolamella (Cai et al., 2018a), meaning that cryo-FIB milling does not grossly perturb canonical nucleosomes *in situ*. We prepared cryolamellae of wild-type BY4741 cells and then collected defocus-phase-contrast tilt series. Cryotomograms of yeast cryolamellae showed that nuclei were packed with nucleosome-like particles (Figure S4). Two-dimensional class averages of template-matched nucleosome-like particles reveal densities that have the approximate size and shape of nucleosomes (Figure S5A). We then subjected the particles belonging to the most nucleosome-like 2-D class averages to 3-D classification, following the original RELION classification workflow. However, none of these 3-D class averages resemble canonical nucleosomes (Figure S5B). While many of the nucleosome-like class averages have dimensions similar to the canonical nucleosome, none of them have densities that resemble the distinctive 1.65 left-handed gyres of DNA. To rule out the possibility that canonical nucleosomes were missed during 2-D classification, we performed direct 3-D classification using 100 classes. None of the resultant 100 class averages resemble a canonical nucleosome (Figure S6).

To determine if our template-matching and classification workflow missed the canonical nucleosomes, we re-did both template matching and 3-D classification of BY4741 wild-type yeast cryolamellae with an intentionally biased reference. Instead of a featureless cylinder, we used the yeast nucleosome crystal structure (White et al., 2001) as the reference (Figure S7A). If canonical nucleosomes are abundant in yeast, they should be detected as a class average that resembles a low-resolution nucleosome crystal structure. No canonical nucleosome class averages were seen in this control experiment (Figure S7B and C).

Our previous study of a HeLa cell (Cai et al., 2018a), which detected canonical nucleosomes, used a Volta phase plate (VPP). VPP data has more low-resolution contrast than defocus phase-contrast data. To test if canonical nucleosomes in yeast cryolamellae are detectable in VPP data, we recorded VPP tilt series and reconstructed tomograms of BY4741 cell cryolamellae (Figure S8). Subtomogram analysis of the VPP tomograms by 2-D classification (Figure S9A), followed by 3-D classification did not reveal a canonical nucleosome class average in BY4741 (Figure S9B). When we performed direct 3-D classification using 100 classes, we detected one class average that resembles a canonical nucleosome (Figure S10A, Movie S1). A second round of classification revealed two types of class averages, one of which resembles a slightly elongated nucleosome (Figure S10B, Movie S2). Refinement of these two types of nucleosomes produced 24 Å-resolution averages that differ in the amount of linker DNA visible (Figures 1C and S10C). The class average that has more ordered linker DNA vaguely resembles the chromatosome, a form of the nucleosome that has linker DNA crossing at the entry-exit site and in contact with a linker histone (Bednar et al., 2017; Zhou et al., 2015). However, the DNA does not appear to cross at the DNA entry-exit site, and, at the present resolution, it is not possible to determine if the linker histone is present. Unlike plunge-frozen complexes, which interact with the air-water interface and have biased orientations (Noble et al., 2018), complexes *in situ* are not subject to such biases. Visualization of the angular distribution of the canonical nucleosomes shows that the disc views are undersampled and likely missed by the classification analysis (Figure S10D), which we also observed in our analysis of HeLa nucleosomes *in situ* (Cai et al., 2018a). This missing hemisphere of views results in roughly half of the canonical nucleosomes going undetected. In summary, the use of VPP and relatively thin (≤ 160 nm) cryolamellae made it possible to detect two canonical nucleosome classes that differ in the amount of ordered linker DNA, similar to what we observed in a HeLa cell (Cai et al., 2018a).

Following our previous work (Cai et al., 2018a), we performed a negative control by analyzing a tomogram of a region in the cytoplasm (Figure S11), which does not have any nucleosomes. We performed template matching using the same reference as for our analysis of nuclei and then direct 3-D classification into 100 classes. None of the resultant class averages resemble a canonical nucleosome (Figure S12), confirming that our analysis was not biased by the cylindrical reference.

Our classification detected only 769 canonical nucleosomes. If we account for the undersampling of disc views (Figure S10D), we estimate there are ∼1,500 canonical nucleosomes detected in the 5 tomograms. In comparison, we estimate that in the single HeLa cell cryolamella cryotomogram (Cai et al., 2018a), there were more than 2,000 nucleosomes in a nucleus volume ∼1/6th of the total analyzed here in BY4741. The percentage of HeLa nucleosomes that are non-canonical *in situ* is unknown and will require further study. To visualize the distribution of canonical nucleosomes, we remapped the two class averages back into their positions in the original tomogram that contains the largest number of canonical nucleosomes (Figure 2). The remapped model shows that the canonical nucleosomes are scattered throughout the sampled nuclear volume. There are no large clusters of canonical nucleosomes like what we saw near the nuclear envelope of a HeLa cell. Using experimentally determined values for nucleosome number and chromatin volume (Oberbeckmann et al., 2019; Uchida et al., 2011), the tomograms we analyzed are expected to hold 25,000 nucleosomes. Therefore, the nucleosomes (both canonical and non-canonical) should pack with an order-of-magnitude higher density than visualized in our remapped model (Figure 2B). In summary, our calculations suggest that the vast majority (> 90%) of the nucleosomes in BY4741 yeast are non-canonical.

**Figure 2.**
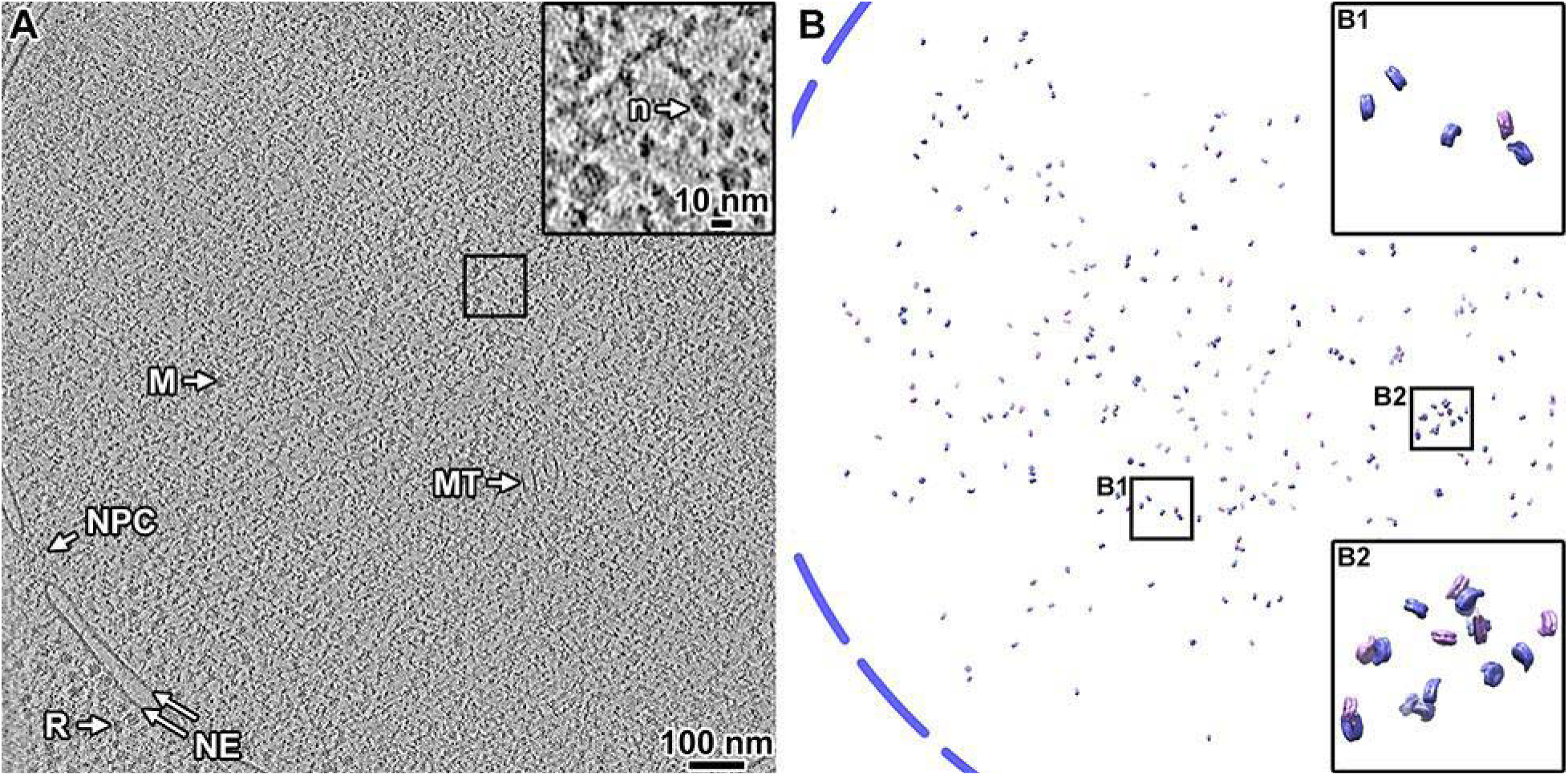
Canonical nucleosomes are a minority of the expected total in wild-type cells. (A) Volta phase plate tomographic slice (12 nm) of a BY4741 cell cryolamella. Large subcellular structures are labeled: nuclear pore complex (NPC), nuclear megacomplex (M), nuclear microtubule (MT), nuclear envelope (NE), and ribosome (R). The inset is a 4-fold enlargement of the boxed area, and a nucleosome-like particle (n) is indicated. (B) Remapped model of the two canonical nucleosome class averages in the tomogram from panel A: the class averages were oriented and positioned in the locations of their contributing subtomograms. The approximate location of the nuclear envelope is indicated by the blue dashed line. The insets B1 and B2 show 4-fold enlargements of the corresponding boxed areas. Note that the remapped model projects the full 150 nm thickness of this cryolamella. In this tomogram, we estimate there are ∼7,600 nucleosomes (see Methods on how the calculation is done), of which 297 are canonical structures. Accounting for the missing disc views, we estimate there are ∼594 canonical nucleosomes in this cryolamella (< 8% the expected number of nucleosomes).

### Histone GFP-tagging and visualization *ex vivo*

Our previous cryo-ET analysis of nucleosomes in a HeLa cell revealed that subtomogram 3-D classification is sensitive to features much smaller than the nucleosomes, as evidenced by the separation of canonical-nucleosome class averages that differ by ∼10 bp of linker DNA near the dyad (Cai et al., 2018a). Furthermore, studies of flagella (Oda and Kikkawa, 2013) and pilus machines (Chang et al., 2016) showed that subtomogram averages of complexes *in situ* can reveal either the presence or absence of protein densities as small as fluorescent proteins. These observations led us to attempt to use a GFP tag to facilitate nucleosome identification *in situ*. Our strategy is to compare subtomogram averages of nucleosome-like particles in strains that express only wild-type histones versus those that express GFP-tagged histones. Note that this tagging strategy did *not* work as intended because we could not detect tagged nucleosomes 3-D classes *in situ*. However, this negative result provided an important clue about the nature of nucleosomes inside yeast cells (see below).

Histones can accept a genetically encoded GFP tag at either the N- or C-terminus. An N-terminal GFP tag is not expected to be visible in subtomogram averages because it would be separated from the histone’s globular domain by the long, flexible N-terminal “tail”. Therefore, we fused GFP to the histone C-terminus, which does not have a flexible tail (Figure S13, A and B) and we further confined the GFP by eliminating the peptide linker that is included in popular GFP-tagging modules (Figure S13C). *S. cerevisiae* has two copies of each histone gene (Figure S13D), which are arranged as gene pairs. The H2A and H2B genes are arranged as gene pairs *HTA1-HTB1* and *HTA2-HTB2* (Hereford et al., 1979). To maximize our chances of detecting nucleosome class averages that have an extra density, we first sought to create strains in which a histone-GFP fusion is the sole source of one class of histones (Figure S13, E and F). We deleted the entire *HTA2-HTB2* gene pair to prevent its amplification as circular DNA molecules by the flanking retrotransposon elements (Libuda and Winston, 2006); the resulting strain is called LGY0012. Next, we inserted the GFP gene at the 3’ end of *HTA1*, without a linker, to generate LGY0016, making H2A-GFP the sole source of H2A. We confirmed LGY0016’s genotype by PCR analysis (Figure S14, A and B), Sanger sequencing, and immunoblots using anti-H2A or anti-GFP antibodies (Figure S14C). Accordingly, the LGY0016 nuclei showed bright fluorescence (Figure S14D). We also constructed the strain LGY0015, which expresses both a H2B-GFP fusion without a linker peptide (Figure S14, E – H) and an untagged copy of H2B. We were unable to create strains that have either H2B-GFP, H3-GFP, or H4-GFP as the sole H2B, H3, and H4 sources, respectively (see H3- and H4-tagging experiments below). Consistent with this low tolerance for a GFP-tagged histone as the sole source of a histone type, the LGY0016 doubling time is ∼50% longer than for wild-type BY4741 (130 minutes versus 85 minutes) in rich media.

We next performed cryo-ET of the nuclear lysates of LGY0016 and LGY0015 cells (Figures S15 and S16). The 2-D class averages of LGY0016 and LGY0015 nuclear lysates resemble those seen in single-particle cryo-EM studies of reconstituted nucleosomes (Chua et al., 2016), though with lower-resolution features (Figure 3A, S17A). Note that 2-D classification uses a circular mask, meaning that it will not bias the shape of the resultant class averages to resemble, for example, the double-lined motifs seen in nucleosome side and gyre views. To increase the number of detected nucleosomes, we used direct 3-D classification into 40 classes. We obtained canonical nucleosome 3-D class averages this way in the lysates of both strains (Figure S18 and S19; Movies S3 and S4). These class averages have the unmistakable structural motifs of canonical nucleosomes, such as a 10-nm diameter, 6-nm thick cylindrical shape, and the left-handed path of the DNA densities. There is more DNA than the crystal structure’s 1.65 gyres because lysate chromatin samples have linker DNA. All these properties are consistent with the subtomogram analysis of nucleosomes from nuclear lysates of wild-type strains BY4741 (Figure 1B) and YEF473A (Cai et al., 2018c).

**Figure 3.**
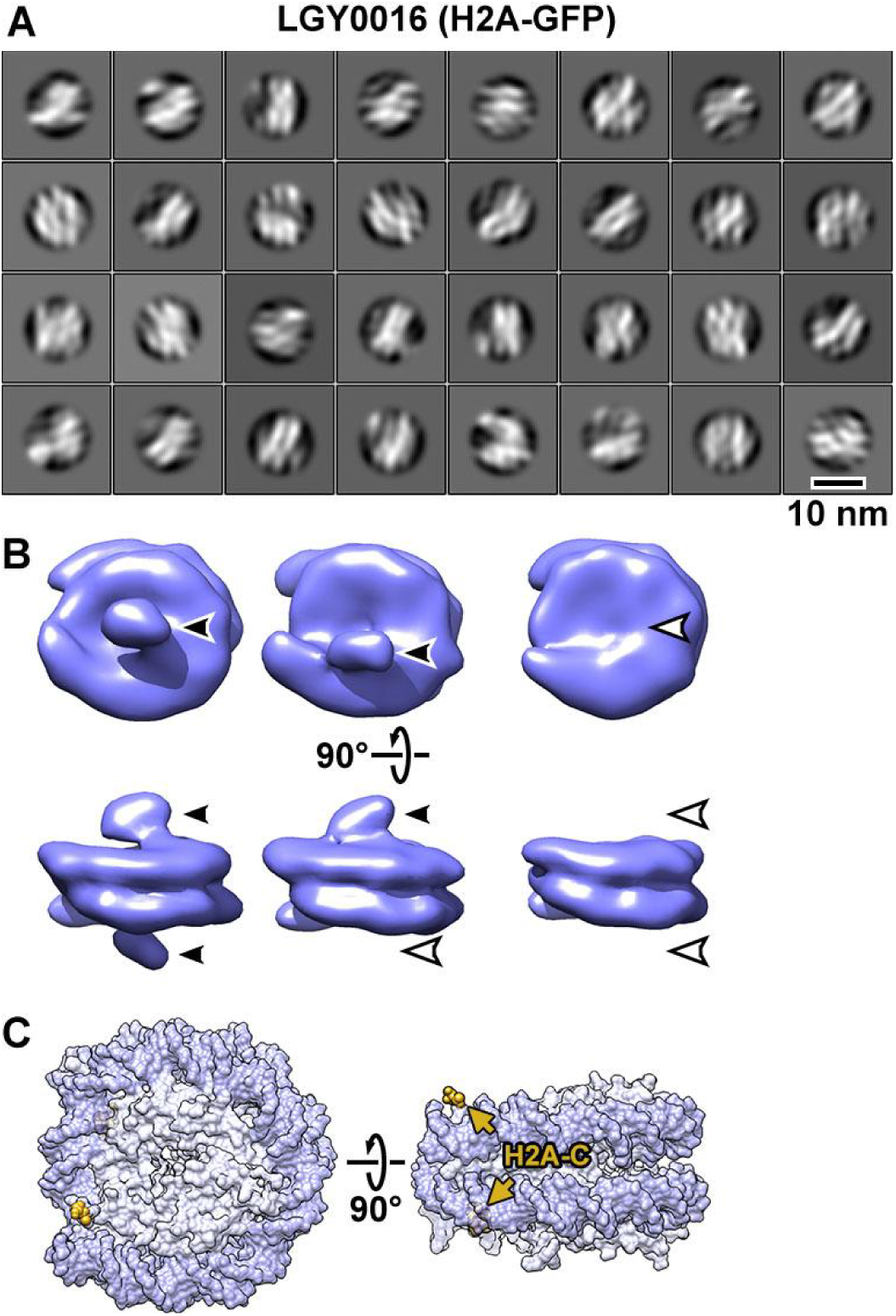
Visualization of GFP-tagged nucleosomes *in vitro*. (A) Example 2-D class averages of nucleosome-like particles that were template-matched with a featureless cylinder reference. Out of 88,896 template-matched particles, 66,328 were retained after 2-D classification. (B) Class averages (3-D) of nucleosomes from nuclear lysates. Solid arrowheads indicate the extra (GFP) densities. Open arrowheads indicate the positions that lack this density. These class averages were obtained after classification directly from subtomogram averaging, without an intervening 2-D classification step. (C) The approximate positions of the H2A C-termini are rendered in yellow and indicated by arrows in the crystal structure of the yeast nucleosome (White et al., 2001). Note that in the crystal structure, fewer of the H2A C-terminal amino acids were modeled than for H2B, meaning that the H2A C-terminus is not perfectly ordered. To facilitate comparison, this structure is oriented like the class averages in panel B. The nucleosome densities in panel B are longer along the pseudo-dyad axis (horizontal) because they have linker DNA, which is absent in the nucleosome crystal structure.

Subsequent classification rounds revealed nucleosome class averages that have an extra density projecting from one or both faces (Figures S18B and S19B). One example of each class (zero, one, or two extra densities) was refined to ∼25 Å resolution (Figure 3B, S17B). The position of the extra density is consistent with the H2A C-terminus being closer to the DNA entry-exit point (Figure 3C) and the H2B C-terminus being far from the DNA entry-exit point (Figure S17C). In principle, LGY0015 cells can assemble nucleosomes that have two copies of H2B-GFP. The absence of LGY0015 nucleosome classes with two densities suggest that they are either unstable or too rare to detect by 3-D classification. We focused our *in situ* cryo-ET analysis on LGY0016 cells because the GFP tags are easier to recognize on nucleosomes from nuclear lysates of this strain.

### Canonical nucleosome classes are not detected in LGY0016 cells *in situ*

In an attempt to detect more yeast nucleosomes *in situ*, we did three types of imaging experiments on LGY0016 cells. We performed cryo-ET of cell cryolamellae with and without the VPP (Figures S20 and S21) and we also imaged cryosections with the VPP (Figure S22). The cryolamellae benefit from the absence of compression artifacts while the cryosections benefit from being thinner on average. We performed template matching and then subjected the hits directly to 3-D classification, which was needed to detect canonical nucleosomes in BY4741 above. The 3-D class averages from all three samples resembled neither canonical nucleosomes nor cylindrical bodies with one or more protruding densities that is expected from the lysate samples (Figures 4, S23, S24, Movie S5). Some class averages have two linear motifs that resemble the double DNA gyres opposite the DNA entry-exit point, but these DNA-like densities do not go 1.65 times around the center of mass as expected of canonical nucleosomes (Luger et al., 1997). Small variations in the classification parameters such as mask size, class number did not reveal any canonical nucleosome classes in LGY0016 cell cryotomograms. We therefore conclude that canonical nucleosomes are rare inside budding yeast nuclei and that non-canonical nucleosomes are the vast majority. At present, we do not know what the non-canonical nucleosome structures are, meaning that we cannot even determine if one non-canonical structure is the majority. Until we know the non-canonical nucleosomes’ structures, we will use the term non-canonical to describe all the nucleosomes that do not have the canonical (crystal) structure.

**Figure 4.**
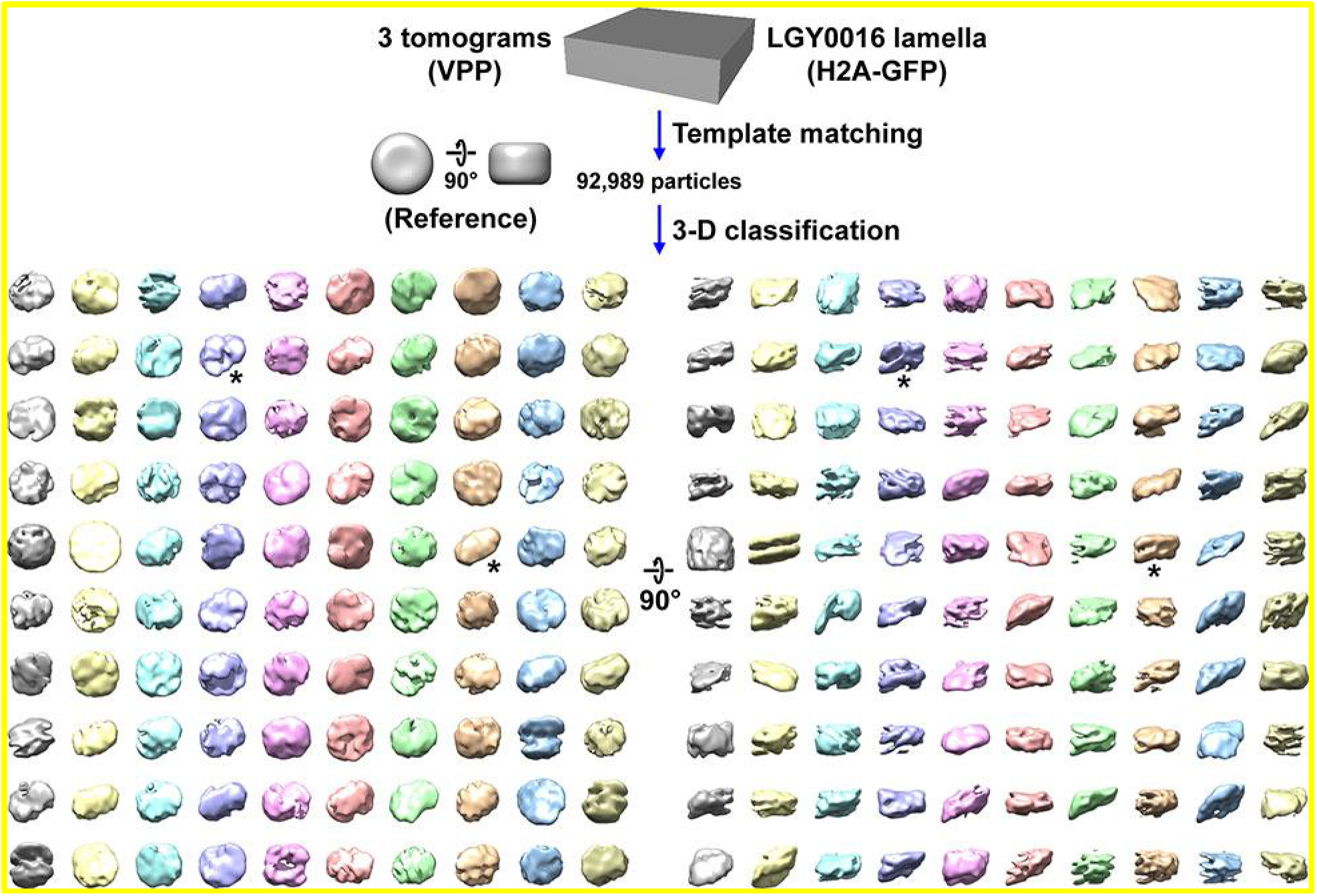
Canonical nucleosome class averages are not detected in LGY0016 (H2A-GFP) cells *in situ*. Class averages (3-D) of nucleosome-like particles in Volta phase plate (VPP) cryotomograms of LGY0016 cryolamellae. The starred classes have two linear motifs. Movie S5 shows the progress of this classification job.

### Ribosome control for sample, data, and analysis pathologies

We may have missed canonical nucleosomes if there were pathologies with either our cryolamellae, data, or workflow, which would result in grossly misclassified and misaligned particles. To test this hypothesis, we performed template matching, 3-D classification and alignment on cytoplasmic ribosomes. Because ribosomes are so large (> 3 megadaltons) and have been studied extensively *in situ*, any of the pathologies listed above would result in either the absence of ribosome class averages or extremely low resolution. Using our control tomogram of the cytoplasm, which is densely packed with ribosomes, we obtained a subtomogram average at between ∼33 Å resolution based on the Fourier shell correlation (FSC) = 0.5 cutoff criterion (or 28 Å using the FSC = 0.143 criterion), from 1,150 particles (**Figure S25**). As a comparison, we simulated densities with the yeast ribosome crystal structure between 15 Å and 30 Å resolution and found that our average has density features consistent with this resolution range. The resolutions of our nucleosome and ribosome averages (∼24 Å and 33 Å) are comparable to recent *in situ* subtomogram averages using similar numbers of particles (∼500 to 1,500) (Laughlin et al., 2022; van den Hoek et al., 2022). Therefore, our cryolamellae, data, and workflow do not show evidence of pathologies.

### GFP tagging of H3 and H4 also does not reveal nucleosomes

In cryolamellae of LGY0016 cells, the absence of nucleosome-like class averages that have an extra density bump suggested that previously unappreciated properties of the H2A-H2B heterodimer make this histone pair a poor candidate for the GFP fusion tag strategy (see Discussion). We therefore revisited GFP tagging of H3 and H4, which are found in all known and speculated forms of nucleosomes and non-canonical nucleosomes (Zlatanova et al., 2009). Because all attempts to make H3- or H4-GFP “sole source” strains failed, we tested strains that had one wild-type copy and one GFP-tagged copy of one of these histones. We tagged members of the *HHF1-HHT1* gene pair, which encode histone H4 (*HHF1*) and H3 (*HHT1*). *HHF1-HHT1* is flanked by the same transposon elements as *HTA2-HTB2*, so gene amplification here would not result in the wild-type copy outnumbering the tagged copy. The genotypes and phenotypes were verified by confirmation PCRs, western blots, Sanger sequencing (not shown), and fluorescence confocal microscopy (Figures S26 and S27). We created H3-GFP strains without a linker (LGY0007) and with linker sequences RIPGLIN (LGY0002) and GGSGGS (LGY0070); this latter linker was introduced previously (Verzijlbergen et al., 2010). For H4, we could only obtain a strain with a H4-GFP-expressing strain by using the flexible GGSGGS linker (LGY0071; Figure S27). Flexible linkers are not expected to facilitate tag-based identification because the tag can occupy a much larger volume and get blurred out as a result but was the only one tolerated in our H4 tagging attempts.

We next prepared nuclear lysates of each of these strains, performed cryo-ET, and subjected them to direct 3-D classification analysis (Figures S28 – S31). Like the other strains, the lysates of each of these four strains had large numbers of canonical nucleosomes (Figures S28A, S29A, S30A, S31A). Of the canonical nucleosomes, we were only able to detect class averages that have the extra density in the LGY0007 nuclear lysates (Figure S28); the nuclear lysates of the other strains do not have nucleosome class averages with a GFP density. Finally, to test if nucleosomes, canonical or non-canonical, with the GFP density bump, could be detected *in situ*, we performed cryo-ET of LGY0007 cell cryolamellae with the VPP (Figure S32). We then performed template matching and subjected the hits to direct 3-D classification analysis. Compared to the BY4741 (wild type) cell cryolamella VPP class averages (Figure S10A), the 3-D class averages in LGY0007 cell cryolamellae resemble neither canonical nucleosomes nor cylindrical bodies with a protruding density (Figure S33). Therefore, H3- and H4-GFP fusions cannot be used to detect non-canonical nucleosomes and much more work needs to be done to identify non-canonical nucleosomes.

## DISCUSSION

Cryo-EM of cells has made the structural analysis of chromatin organization *in situ* feasible. The earliest *in situ* studies were done with projection cryo-EM images and 2-D Fourier analysis of cryosectioned cells, which revealed that long-range order is absent in chromatin *in situ* (Eltsov et al., 2008; McDowall et al., 1986). Cryo-ET studies later showed that short-range order is present in isolated chicken erythrocyte nuclei (Scheffer et al., 2011), but absent in picoplankton and budding yeast (Chen et al., 2016; Gan et al., 2013). These early cryo-ET studies revealed little about nucleosome structure *in situ* because, as Eltsov *et al* observed, the nucleosomes in those samples appeared like smooth ellipsoids (Eltsov et al., 2018). Recent cryo-ET studies using state-of-the-art electron-counting cameras and energy filters have revealed that in cryotomographic slices of cell nuclei, there are densities that resemble nucleosome side and gyre views in which gyre-like features are resolved (Cai et al., 2018a; Cai et al., 2018b; Eltsov et al., 2018). This higher-fidelity data made it possible to use 3-D classification to detect canonical nucleosomes in nuclear envelope-associated chromatin in a HeLa cell (Cai et al., 2018a).

Our use of 3-D classification detected canonical nucleosomes in wild-type yeast nuclear lysates and canonical nucleosomes both with and without GFP tags in nuclear lysates of some strains that bear histone-GFP fusions. We also detected a canonical nucleosome class average in wild-type cell cryolamellae imaged with a VPP. The *in situ* detections suggest that there are only ∼1,500 canonical nucleosomes (taking account of the undersampling of nucleosomes in the disc view) out of the 25,000 nucleosomes expected of the total sampled nuclear volume. Note that the percentage of canonical nucleosomes in lysates cannot be accurately estimated because we cannot determine how many nucleosomes in total are in each field of view. The estimates from the cryolamellae are more reliable because the expected numbers of nucleosomes (canonical or not) can be estimated from genomics, biochemical, and X-ray tomography data (see Methods). When we analyzed LGY0016 cell cryolamellae, in which H2A-GFP is the sole source of H2A, we did not detect canonical nucleosome classes or any cylindrical nucleosome-sized structures that have extra density bumps. Likewise, analysis of cryolamellae of LGY0007 did not reveal either a canonical nucleosome-like class average or any cylindrical nucleosome-sized density with an extra density bump, even though H3-GFP should be incorporated into the H3-H4 tetramer, the central component of the nucleosome.

The absence of either a canonical nucleosome-like class average that has an extra density bump, or a cylinder with an extra density bump in the LGY0016 strain suggests that the H2A-H2B heterodimer is mobile *in situ*. By “mobility”, we are not implying that H2A-H2B is dissociated. We mean that H2A-H2B is attached to the rest of the nucleosome and can have small differences in orientation. The H2A/H2B heterodimers are in contact with the H3/H4 heterotetramer because crosslinking and single-particle fluorescence imaging experiments showed that approximately 80% of H2B – and presumably H2A to which it stably dimerizes – is bound to chromatin *in situ* (Mohan et al., 2018; Ranjan et al., 2020). H2A-H2B mobility was hinted at by the observations from X-ray crystallography that yeast nucleosomes lack the hydrogen bonds that stabilize the two H2A-H2B heterodimers in metazoan nucleosomes (White et al., 2001) and that interfaces between H2A-H2B and H3 are moderately exposed to small-molecule probes (Marr et al., 2021). Furthermore, recent simulations show that the H2A/H2B heterodimer may adopt numerous orientations and small displacements, with either fully or partially unwrapped DNA, while remaining in contact with H3-H4 (Ishida and Kono, 2022). We also cannot rule out the possibility that expression of H2A-GFP makes nucleosomes less ordered *in situ*.

Like LGY0016 (H2A-GFP sole source), none of the LGY0007 (H3 plus H3-GFP) *in situ* nucleosome-like class averages had an extra density consistent with the GFP density tag (only seen in lysates). Furthermore, there were no canonical nucleosome class averages – either with or without the GFP density – seen among the class averages of LGY0007 cell cryolamella nucleosome-like particles. These observations suggest that the expression of H3-GFP makes nucleosomes less ordered *in situ*. Altogether, our experiments show that in budding yeast, canonical nucleosomes are rare *in situ* and that the expression of GFP-tagged histones leads to further perturbations of nucleosome structure *in situ*.

Our data is consistent with a model in which yeast canonical nucleosomes are abundant *ex vivo* (Figure 5A). They adopt multiple non-canonical conformations *in situ* (Figure 5, B and C) and rarely adopt the canonical one (Figure 5D). Note that the blurring in these panels implies the heterogeneity in the positions of the DNA and proteins, not their actual motion. In addition to H2A-H2B and DNA conformational heterogeneity, diverse nucleosome-associated proteins that bind to multiple positions (Figure 5C) would further increase the heterogeneity. A high abundance of non-canonical nucleosomes means that more of the genome would be accessible, consistent with yeast having high levels of transcription. Nucleosome heterogeneity is also consistent with the absence of chromatin long-range order *in situ* because crystalline oligonucleosome arrays can only form if sequential nucleosomes adopt nearly identical conformations (Ekundayo et al., 2017; Robinson et al., 2006; Song et al., 2014). Further investigation is needed to identify the biochemical and biophysical factors responsible for the abundance of non-canonical nucleosomes in yeast and to determine their diverse structures (Zlatanova et al., 2009).

**Figure 5.**
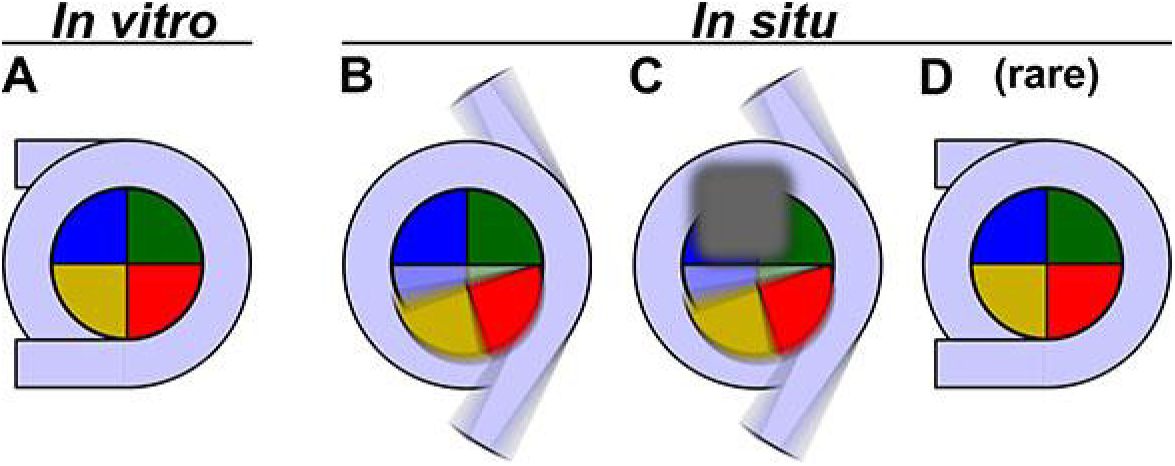
Models of yeast nucleosome heterogeneity. Schematics of DNA (light blue) and histones (shaded pie slices) in the nucleosome disc view. The cartoons only illustrate the 147 bp of “core” DNA. (A) Canonical nucleosome, in which all the histones and 147 bp of DNA are part of an ordered complex. (B) Nucleosome with alternative histone H2A-H2B (yellow, red) conformations and partially dissociated DNA. (C) Nucleosome bound to non-histone proteins (gray). The blurred gray box represents different proteins that can bind, thereby contributing to constitutional heterogeneity. The blurred appearance represents a large range of positions and orientations that protein and DNA components adopt inside cells, which would result in the absence of a class average resembling a canonical nucleosome. (D) Canonical nucleosomes are a minority conformation *in situ*.

In combination with our previous work on a HeLa cell (Cai et al., 2018a), we show that globular complexes as small as canonical nucleosomes (∼200 kilodaltons) can be detected by 3-D classification of subtomograms from cryolamellae that were imaged with a VPP. Given the low throughput of cryo-FIB milling, the low yield of cryolamellae thinner than 160 nm, and the technical difficulties of VPP imaging, it is not yet feasible to systematically determine the conditions needed to detect sub-200 kDa complexes by purification *in silico*. More depositions of eukaryote cryolamellae cryo-ET datasets in the public database EMPIAR (Iudin et al., 2016) may enable a thorough search of data-collection and processing-parameter space.

Our *in situ* cryo-ET-based model adds to the body of work on non-canonical nucleosomes. ChIP-seq analysis has detected sub-nucleosomes *in situ*, though the abundance was unknown (Rhee et al., 2014). MNase-seq has also detected nucleosomes in a partially unwrapped and partially disassembled state *in situ* (Ramachandran et al., 2017). A recent Hi-C variant “Hi-CO” presented evidence that < 147 bp of yeast nucleosomal DNA is protected from MNase attack (Ohno et al., 2019), which is consistent with the partial detachment of core DNA. Molecular dynamics advances in all-atom and coarse-grained simulations are now showing that nucleosomes are far more dynamic than previously appreciated (Armeev et al., 2021; Brandani et al., 2021; Farr et al., 2021; Huertas et al., 2021; Ishida and Kono, 2021). The DNA-unwrapping associated with nucleosome breathing was shown to disfavor ordered helical oligonucleosome structures (Farr et al., 2021). Nucleosome breathing is also evident in high-throughput atomic-force microscopy experiments, which revealed that only ∼30% of nucleosomes are fully wrapped, and that approximately half of the nucleosomes have an opening angle (measured between the entry/exit DNA arms) 60° larger than the fully wrapped one (Konrad et al., 2021). Nucleosomes that have non-canonical nucleosome properties, such as lower stability or exposure of internal surfaces, have been reported in fission yeast (Koyama et al., 2017; Sanulli et al., 2019), which may explain why we did not observe canonical nucleosomes in cryosections in those cells either (Cai et al., 2018b). Some nucleosomes in human and fly cell cryosections appear “gaping”, i.e., with the inter-DNA-gyre distance slightly larger than ∼2.7 nm (Eltsov et al., 2018). Partial DNA detachment has been seen in complexes between nucleosomes and remodelers and transcription factors (Eustermann et al., 2018; Farnung et al., 2017; Liu et al., 2017; Sundaramoorthy et al., 2018; Sundaramoorthy et al., 2017; Willhoft et al., 2018), methyltransferases (Bilokapic and Halic, 2019; Jang et al., 2019), or transcription-related complexes (Dodonova et al., 2020; Kujirai et al., 2018). Larger amounts (up to ∼25 bp) of DNA detachment have been seen very rarely in cryo-EM structures (Bilokapic et al., 2018; Zhou et al., 2021). A recent study of cryosectioned fly embryos has also presented evidence of nucleosome-like structures such as hemisomes and three-gyre structures (Fatmaoui et al., 2022). The prevalence and functional consequences of non-canonical nucleosomes *in situ* remain to be studied in other organisms.

### Alternative hypotheses

We now consider the alternative hypothesis that canonical nucleosomes are the dominant form *in situ* and that we have missed them as a result of our image processing. Many of the 3-D class averages appear to have multiple gyre-like densities, which may arise from nucleosome stacking. However, classes with multiple gyre-like densities are also found in the class averages from the cytoplasm, which does not have nucleosomes (Figure S12). These are “junk” classes that result from the averaging of different particle species into the same class. Importantly, stacked canonical nucleosomes are so conspicuous that they could not have been missed in tomographic slices. For instance, two stacked canonical nucleosomes have twice the mass of a single canonical nucleosome and would have dimensions 12 nm by 10 nm (Figure S34). Multiple stacked canonical nucleosomes would appear like 10-nm-thick filaments.

Another alternative hypothesis is that the lack of disc views, corresponding to nucleosomes whose “face” is parallel to the cryolamella surface, resulted in canonical nucleosomes going undetected. This hypothesis would be supported only if the orientations of canonical nucleosomes were biased. However, biased orientation in cryo-EM experiments is usually a result of interactions with the air-water interface (Noble et al., 2018), which does not exist for nucleosomes inside the nucleus of a cell. Another hypothesis is that the missing views resulted in elongated class averages because of the missing wedge or missing cone in Fourier space. This artifact would only affect the distribution of canonical versus non-canonical nucleosomes if canonical nucleosomes have a preferred orientation, which we have just argued against due to the nature of our cellular samples. The class averages from non-disc-view nucleosomes do not have a missing-wedge/cone artifact because they include nucleosomes whose superhelical axes are in the X-Y plane; when averaged together, the Fourier transforms of these nucleosomes “fill in” Fourier space because they sample numerous rotations about their super-helical axes. Missing-wedge-free reconstructions are exemplified by plunge-frozen actin filaments, which lie parallel to the EM grid. As illustrated in a recent study (Merino et al., 2018), reconstructions of filamentous actin are free of missing-wedge distortions even though the axial views are completely absent.

Some of the particles in the tomographic slices resemble donuts. Because nucleosome disc views also resemble donuts, it is possible that we completely missed these particles in either our template matching or classification analyses. As stated above, even if we missed all the disc views, our conclusions would not change because the orientations of canonical nucleosomes inside of cellular samples are not biased.

Furthermore, the donut-like particles cannot be canonical nucleosomes because they are too wide (> 12 nm). A subset of them were detected by template matching and classification and separated into their own class, which resembles a cylinder with a rounded cap (Movie S5, row 5 column 1).

Another hypothesis for the low numbers of detected canonical nucleosomes is that the nucleoplasm is too crowded, making the image processing infeasible. However, crowding is an unlikely technical limitation because we were able to detect canonical nucleosome class averages in our most-crowded nuclear lysates, which are so crowded that most nucleosomes are butted against others (Figures S15 and S16). Crowding may instead have biological contributions to the different subtomogram-analysis outcomes in cell nuclei and nuclear lysates. For example, the crowding from other nuclear constituents (proteins, RNAs, polysaccharides, etc.) may contribute to *in situ* nucleosome structure, but is lost during nucleus isolation.

### Limitations of the study

Because naked linker DNA cannot yet be seen *in situ* in yeast, the connectivity between the nucleosomes (both canonical and non-canonical) could not be followed. The subtomogram analysis was unable to resolve the structures of the non-canonical nucleosomes. Because of the low resolution, we could not assess the variability in the structure and composition of the canonical nucleosomes. All of these limitations could potentially be addressed by increasing the data quality and the numbers of classes and subtomograms per class, which require advances at all stages of structural cell biology. These advances include improvements in cryo-FIB milling throughput and reproducibility (Buckley et al., 2020; Tacke et al., 2021; Zachs et al., 2020), cryo-EM cameras, laser phase plates (Turnbaugh et al., 2021), template-matching / segmentation software (Bepler et al., 2020; Moebel et al., 2021), and subtilt refinement software such as emClarity (Himes and Zhang, 2018), EMAN2 (Chen et al., 2019), Warp/M (Tegunov et al., 2021) and RELION 4 (Zivanov et al., 2022). Because structural cell biology is still a nascent field, the optimum combination of sample prep, imaging, and data analysis will require thorough exploration of the parameter space at each step. The nucleosome VPP subtomogram averages presented here were limited to ∼24 Å, even though the data was recorded close to focus as suggested by a higher-resolution study of purified ribosomes (Khoshouei et al., 2017). Limiting factors include lower particle numbers, alignment inaccuracy, VPP charging, and the lack of contrast transfer function (CTF) modeling. In single-particle cryo-EM analysis, high-resolution analysis (3 Å or better) requires accurate modeling and then compensation for the CTF. A key first step of CTF modeling is the visualization of Thon rings in Fourier power spectra. We plotted Fourier power spectra in both 2-D with CTFFIND (Rohou and Grigorieff, 2015) and as a 1-D rotational average with IMOD (Mastronarde, 1997), but could not see Thon rings (Figure S35), meaning that CTF compensation is not feasible. A study of VPP single-particle cryo-EM analysis recommends intentionally underfocussing to 500 nm to increase the number of Thon rings, making CTF modeling easier (Danev et al., 2017). However, the samples used in that study were much thinner (plunge-frozen proteasomes) and the dose used per image was much higher (∼40 electrons / Å^2^). Because the subtomogram analysis of smaller complexes inside cryolamellae faces multiple challenges, more investigation is needed to determine the optimum parameters for increased resolution.

Another challenge is the positive identification of the non-canonical nucleosome species. The approaches attempted here (the use of a GFP tag) failed. An alternative approach would be to remap the various averages and show that the linker DNA of sequential non-canonical nucleosomes point to each other, as we have previously done for HeLa chromatin *in situ* (Cai et al., 2018a). Unfortunately, this approach is not suitable for yeast because only the canonical class averages have linker-DNA densities – none of the other class averages have densities that resemble linker DNA. Because non-canonical yeast nucleosomes *in situ* are unknown structures, we would have to identify individual instances using a “GFP of cryo-EM”, in the form of a compact fusion protein-like structure that can be directly visualized in tomographic slices without classification and subtomogram averaging. Several attempts have been made in recent decades (Diestra et al., 2009; Mercogliano and DeRosier, 2007; Nishino et al., 2007; Wang et al., 2011). More work is needed to test the suitability of such tags as “GFPs of cryo-EM” *in situ*.

## ACKNOWLEDGMENTS

We thank Kerry Bloom, Vu Nguyen, and Carl Wu for advice on fluorescent-protein tagging of histones; Bill Rice and Ed Eng for help with cryo-EM data collection; John Heumann for discussions about PEET and for implementing a parallelized duplicate-removal routine; Rado Danev, Kliment Verba, and Shenping Wu for advice on the VPP. The Quadro P6000 used in this work was kindly donated by the NVIDIA Corporation. ZYT, SC, JKC, and LG were supported by a Singapore Ministry of Education Tier 1 grant R-154-000-A49-114 and Tier 2 grant MOE2019-T2-1-140. AJN was supported by a grant from the NIH National Institute of General Medical Sciences (NIGMS) (F32GM128303). Some of this work was performed at the Simons Electron Microscopy Center and National Resource for Automated Molecular Microscopy located at the New York Structural Biology Center, supported by grants from the Simons Foundation (SF349247), NYSTAR, and the NIH National Institute of General Medical Sciences (GM103310) with additional support from NIH S10 RR029300.

## AUTHOR CONTRIBUTIONS

SC, ZYT, AJN, JKC, LG – project design, experiments, writing; JS, LG – training.

## MATERIALS AND METHODS

### Yeast Strain and Growth Conditions

All yeast strains were streaked on yeast extract peptone dextrose (YPD) agar plates (2% w/v peptone, 1% w/v yeast extract, 2% w/v glucose for liquid media; additional 2% w/v agar for agar plates) at 30°C and cultured in YPD in conical flasks shaking at 200 – 250 RPM at 30°C. All yeast strains are of mating type a. Modifications or lack thereof to all histone genes were authenticated by PCR and Sanger sequencing (Bio Basic Asia Pacific Pte Ltd, Singapore) in all yeast strains.

### Bacterial Growth Conditions

All plasmids were provided in DH5-Alpha *Escherichia coli*. The bacteria were streaked on LB agar plates with ampicillin (40 g/L LB Broth with agar (Miller), 100 μg/mL ampicillin) at 37°C and cultured in LB liquid medium with ampicillin (25 g/L LB Broth (Miller), 100 μg/mL ampicillin) in vent cap tubes shaking at 200 – 250 RPM at 37°C.

### Plasmid extraction and linearization

The plasmid pFA6a-GFP(S65T)-His3MX6 (Longtine et al., 1998) was a gift from John Pringle (Addgene 41598; Addgene, Watertown, MA) and pFA6a-link-yoTagRFP-T-Kan (Lee et al., 2013) was a gift from Wendell Lim & Kurt Thorn (Addgene 44906); both were given in the form of bacterial stabs. Note that pFA6a-link-yoTagRFP-T-Kan is the source of the KanR marker used to delete HTB2-HTA2. Five mL of bacteria were cultured overnight, then plasmids were extracted with the QIAprep Spin Miniprep Kit (QIAGEN, Hilden, Germany) following the manufacturer’s instructions.

Extracted plasmids were linearized by double digestion with a reaction containing 1 µg of plasmid DNA, 5 µL of 10× rCutSmart™ Buffer (New England BioLabs, Ipswich, MA), 10 units each of *Sal*I and *EcoR*V restriction enzymes (New England BioLabs) topped up to 50 µL with nuclease-free water. The reaction mixture was heated at 37°C for 15 minutes for the digestion reaction, then 80°C for 20 minutes to inactivate the enzymes.

### Strain construction

The strain details are shown in Table S1. Primers were from IDT (Integrated DNA Technologies, Inc., Singapore) and listed in Table S2. Q5 PCR Master Mix (New England BioLabs) was used for PCR reactions. BY4741 (Brachmann et al., 1998) served as the wild-type strain. Tagging and deletion cassettes were amplified from linearized pFA6a plasmids (Lee et al., 2013; Longtine et al., 1998) with a PCR reaction containing 1 ng of template DNA and 0.5 μM of each primer, using a PCR program of 98°C for 30 seconds, 30 cycles of 98°C for 5 seconds, 60°C for 10 seconds and 72°C for 1.5 minutes, then 72°C for 5 minutes.

Cells were transformed using the lithium acetate / PEG4000 method reported in (Nishimura and Kanemaki, 2014). Overnight cell culture in YPD was diluted to 0.1 OD_600_ in 25 mL of YPD and grown to 0.3 OD_600_. Ten mL of cells were collected, centrifuged at 1,600 *× g* at 25°C for 3 minutes, and the supernatant was removed. The cells were then washed twice with 10 mL sterile water with centrifugation at 1,600 *× g* at 25°C for 3 minutes. The cells were resuspended in 1 mL sterile water, transferred to a new 1.5 mL collection tube, centrifuged at 17,900 × *g* at 25°C for 1 minute, washed in 1 mL of TE/LiAc (10 mM tris, 1 mM ethylenediaminetetraacetic acid (EDTA), 100 mM lithium acetate), with centrifugation at 17,900 × *g* for 1 minute, and resuspended in 50 μL of the same buffer. Fifty μL of cell suspension was transferred to a new 1.5 mL collection tube containing 5 μL of 10 mg/mL salmon sperm DNA (Sigma-Aldrich, Burlington, MA) plus 5 μL of PCR-amplified cassette DNA. 360 µL of TE/LiAc/PEG (10 mM tris, 1 mM EDTA, 100 mM lithium acetate, 40% w/v Polyethylene glycol 4000) was added and incubated with shaking at 25°C for 30 minutes. Forty µL of dimethyl sulfoxide was added, the suspension was incubated at 42°C for 15 minutes in a water bath, then cooled on ice for 2 minutes. The suspension was centrifuged at 13,000 x *g* for 1 minute, the liquid was removed, and the cells were resuspended in 300 μL of 1× TE buffer, pH 8.0 (10 mM tris, 1 mM EDTA). Two hundred μL of this cell suspension was plated on a selection plate and incubated for several days at 30°C. Histidine selection plates were created with 6.7 g/L yeast nitrogen base without amino acids (Sigma-Aldrich), 1.92 g/L yeast synthetic drop-out medium supplements without histidine (Sigma-Aldrich), 2% w/v glucose and 2% w/v agar. G418 selection plates were created by adding G418 to the molten agar (YPD for G418 single selection, histidine auxotrophy medium for double selection) to a concentration of 200 mg/L before it was poured.

Transformants were verified by PCR and Sanger sequencing. Genomic DNA was extracted with the DNeasy Blood & Tissue Kit (QIAGEN) following the manufacturer’s instructions. Confirmation PCR was performed with 1 µg of template genomic DNA and 0.5 μM of each primer, using a PCR program of 94°C for 2 minutes, 30 cycles of 94°C for 1 minute, 62°C for 1 minute and 72°C for 3.5 minutes, then 72°C for 5 minutes. PCR products were purified with the QIAquick PCR Purification Kit (QIAGEN) following the manufacturer’s instructions, with water used for the final elution before they were sent for Sanger sequencing.

### DNA gels and immunoblots

PCR products were electrophoresed in 2% agarose in Tris-acetate-EDTA and visualized with FloroSafe DNA Stain (Axil Scientific Pte Ltd, Singapore). The electrophoresis was performed at 100 Volts for 60 – 80 minutes before visualization with a G:Box (Syngene).

To generate protein samples for immunoblot analysis, ∼20 OD_600_ units of yeast cells were pelleted and stored at −80°C for at least 1 hour. Cells were then resuspended in 200 µL of ice-cold 20% trichloroacetic acid (TCA) and vortexed with glass beads at 4°C for 1 minute 4 times. For each sample, 500 µL of ice-cold 5% TCA was added, mixed with the pellet, then transferred to a new 1.5 mL collection tube. Another 500 µL of ice-cold 5% TCA was mixed with each pellet and transferred to the same 1.5 mL collection tube as before, so that each collection tube had 1 mL total volume. The tubes were then left on ice for 10 minutes and centrifuged at 15,000 *× g* at 4°C for 20 minutes. The TCA was aspirated from the tube, then the pellet was resuspended in 212 µL of Laemmli sample buffer with 2-mercaptoethanol added (Bio-Rad, Hercules, CA). To neutralize the residual TCA, 26 µL of 1 M Tris, pH 8 was added. This mixture was heated at 95°C for 5 minutes, centrifuged at 25°C at 15,000 *× g* for 10 minutes, then 5 µL of the supernatant was subjected to SDS-PAGE.

The primary and secondary antibodies used for immunoblots are shown in Table S3. Proteins were electrophoresed in 10% Mini-PROTEAN® TGX™ Precast Protein Gels (Bio-Rad). Either Precision Plus Protein™ WesternC™ Standards (Bio-Rad) or Invitrogen™ MagicMark™ XP Western Protein Standard (Thermo Fisher Scientific, TFS, Waltham, MA) served as the ladder. The proteins were electrophoresed at 100 Volts for 1 hour at 25°C. The gels were transferred onto Immun-Blot® PVDF Membranes (Bio-Rad) at 4°C in transfer buffer (3.02 g/L tris, 14.4 g/L glycine, 20% methanol). The transfer was performed at 100 Volts for 30 minutes. The membranes were then blocked with 2% BSA in 1× Tris-Buffered Saline, 0.1% Tween® 20 Detergent (TBST) for 1 hour. This was followed by 50 µg/mL avidin in TBST for 30 minutes, then a wash with TBST for 20 minutes if Precision Plus Protein™ WesternC™ Standards ladder was used. All antibody dilution factors are reported in Table S3. The membranes were probed with primary antibodies at the stated dilutions in 2% BSA in TBST for 1 hour at 25°C. Membranes were washed with TBST for 20 minutes three times at 25°C. The membranes were then probed with secondary antibodies in 2% BSA in TBST for 1 hour, with additional Precision Protein™ StrepTactin-HRP Conjugate (Bio-Rad) at 1:10,000 dilution if Precision Plus Protein™ WesternC™ Standards ladder was used. Finally, the membranes were washed with TBST for 10 minutes 3 times, then treated with a 50:50 mixture of Clarity Western Peroxide Reagent and Clarity Western Luminol/Enhancer Reagent (Bio-Rad) for 5 minutes before visualization by chemiluminescence on an ImageQuant LAS 4000 (Cytiva, Marlborough, MA).

### Fluorescence microscopy

Cells were grown to log phase (OD_600_ = 0.1 – 1.0), of which 2 OD_600_ units of cells were collected, pelleted at 5,000 *× g* for 1 minute, then resuspended in 1 mL of YPD. Four µL of cell culture was then applied to a glass slide and pressed against a coverslip. The cells were imaged live at 23°C with an Olympus FV3000 Confocal Laser Scanning Microscope (Olympus, Tokyo, Japan) equipped with a 1.35 NA 60× oil-immersion objective lens. GFP fluorescence was acquired using the 488 nm laser-line, with a DIC image recorded in parallel with the fluorescence image. Images were captured as Z-stacks thick enough to sample the GFP signals through all the nuclei in each stage position. Additional details regarding data collection are shown in Table S4.

### Preparation of nuclear lysates

Yeast nuclei were prepared with reagents from the Yeast nuclei isolation kit (Abcam 206997, Cambridge, United Kingdom), unless noted otherwise. Yeast cells (30 mL, OD_600_ ∼1) were pelleted at 3,000 *× g* for 5 minutes at 25°C. The pellet was washed twice with 1 mL water (3,000 *× g*, 1 minute). The pellet was then resuspended in 1 mL Buffer A (pre-warmed to 30°C) containing 10 mM dithiothreitol. The suspension was incubated in a 30°C water bath for 10 minutes. Cells were then pelleted at 1,500 *× g* for 5 minutes and then resuspended in 1 mL Buffer B (prewarmed to 30°C) containing lysis enzyme cocktail (1:100 dilution). The suspension was incubated in a 30°C shaker for 15 minutes for cell-wall digestion and then pelleted at 1,500 *× g* for 5 minutes at 4°C. The pellet was resuspended in 1 mL pre-chilled Buffer C with protease inhibitor cocktail (1:1,000 dilution). The cells were lysed by 15 up-and-down strokes with a pre-chilled glass Dounce homogenizer on ice. The lysate was incubated with shaking for 30 minutes at 25°C. The cell debris was pelleted at 500 *× g* for 5 minutes at 4°C. The supernatant was then transferred to a new tube. The nuclei were pelleted at 20,000 *× g* for 10 minutes at 4°C. The nuclear pellet was resuspended in 10 – 20 µL pre-chilled lysis buffer (50 mM EDTA and 1:1,000 protease inhibitor cocktail dilution) and incubated for 15 minutes on ice.

Nuclei lysates (3 µL) were added to a glow-discharged CF-4/2-2C-T grid (Protochips, Morrisville, NC). The grid was plunge-frozen using a Vitrobot Mark IV (blot time: 1 second, blot force: 1, humidity: 100%, temperature: 4°C).

### Preparation of cryosections

Self-pressurized freezing was done based on a modified version of a published protocol (Yakovlev and Downing, 2011). Yeast cells (30 mL, OD_600_ = 0.2 – 0.6) were pelleted and resuspended in a dextran stock (40 kilodalton, 60% w/v, in YPD) to a final concentration of 30%. Cells were then loaded into a copper tube (0.45 / 0.3 mm outer / inner diameters). Both ends of the tube were sealed with flat-jaw pliers. The tube was held horizontally and dropped into the liquid-ethane cryogen. The tube’s ends were removed under liquid nitrogen with a tube-cut tool (Engineering Office M. Wohlwend, Sennwald, Switzerland).

Gold colloid solution (10-nm diameter, 5 µL at 5.7 ×10^12^ particles/mL in 0.1 mg/mL BSA) was applied to a continuous-carbon grid (10-nm thick carbon) and then air-dried overnight. Cryosections were controlled by a custom joystick-based micromanipulator (MN-151S, Narishige Co., Ltd., Tokyo, Japan) (Ladinsky et al., 2006; Ng et al., 2020). Seventy nm-thick frozen-hydrated sections were cut at −150°C in a Leica UC7/FC7 cryo-ultramicrotome (Leica Microsystems, Vienna, Austria). The EM grid was positioned underneath the ribbon using a Leica micromanipulator (Studer et al., 2014). The cryosection ribbon (∼3 mm long) was then attached to the grid by operating the Crion (Leica Microsystems) in “charge” mode for ∼30 seconds (Pierson et al., 2010).

### Preparation of cryolamellae

Cells were plunge-frozen and then cryo-FIB milled using the method of Medeiros and Bock (Medeiros et al., 2018) as follows. Immediately before plunge-freezing, mid-log phase (OD_600_ ∼0.6) yeast cells were pelleted at 4,000 *× g* for 5 minutes. They were then resuspended in YPD media containing 3% (v/v) dimethyl sulfoxide as cryo-protectant to a final OD_600_ of approximately 2.5. Four µL of the cells were subsequently deposited onto Quantifoil R2/4 200 mesh copper grids (Quantifoil Micro Tools GmbH, Jena, Germany), which were then manually blotted from the back with Whatman® Grade 1 filter paper for approximately 3 – 5 seconds. The grids were then plunged into a 63/37 propane/ethane mixture (Tivol et al., 2008) using a Vitrobot Mark IV (humidity: 100%, temperature: 4°C). Cryo-FIB milling was performed on a Helios NanoLab 600 DualBeam (Thermo Fisher Scientific, TFS, Waltham, MA) equipped with a Quorum PolarPrep 2000 transfer system (Quorum Technologies, Laughton, United Kingdom). Plunge-frozen yeast samples were coated with a layer of organometallic platinum using the in-chamber gas injection system and the cold-deposition method (Hayles et al., 2007). Cryolamellae were then generated as follows: bulk material was first removed using the FIB at 30 kV 2.8 nA, followed by successive thinning of the cryolamellae at lower currents of 0.28 nA and 48 pA.

### Cryo-ET data collection and reconstruction

All cryo-ET data were collected on Titan Krioses (TFS). Tilt series were collected with either TFS Tomo4, Leginon (Suloway et al., 2009), SerialEM (Mastronarde, 2003), or PACE-tomo (Eisenstein et al., 2022). Images were recorded either on a Falcon II (TFS) in integration mode or as movie frames on a K2 or K3 summit camera (Gatan, Pleasanton, CA) in super-resolution mode. The pixel sizes for the K2 and K3 data were chosen so that when binned to the same level, they closely match the ∼7 Å pixel size in our previous *in situ* study of HeLa chromatin (Cai et al., 2018a). Smaller pixel sizes were chosen for Falcon II data because this camera has lower detective quantum efficiency than the K-series cameras (Ruskin et al., 2013). Movies were aligned with either MotionCor2 (Zheng et al., 2017) or IMOD alignframes (Mastronarde, 1997). Prior to starting data collection of cryolamellae, the stage was pre-tilted to either −10° or −15° to account for the milling angle. Additional data-collection details are shown in Table S5.

Volta Phase Plate (VPP) imaging was done using the protocol of Fukuda and colleagues (Fukuda et al., 2015). The VPP position and condenser stigmator were adjusted such that the Ronchigram (the projected pattern visible in the camera) was free of elongated features, indicating no astigmatism, and is flat rather than grainy, indicating that the VPP is on-plane. Notably, the VPP needs to be heated, or else the phase shift will rapidly exceed π/2 radians (Danev et al., 2014). Two different VPP assemblies were used, which we will refer to as the NYSBC-2019 and NUS-2020 ones. These VPPs required different heater power settings to maintain a stable phase shift. The NYSBC-2019 and NUS-2020 VPP heaters were operated at ∼100 mW and ∼370 mW, respectively.

Cryotomograms were reconstructed using IMOD’s *eTomo* workflow (Mastronarde, 1997). Tilt series of lysates and cryosections were aligned using the gold beads as fiducials while those of cryolamellae were aligned with patches as fiducials. Only the tilt series that exhibited the minimal amount of drift and sample warping were analyzed. To further improve the tilt series alignment around the chromatin, fiducials (beads or patches) were chosen on the chromatin regions. For cryolamella patch tracking, tilt series were binned to a pixel size of 6.8 Å, the “Break contours into pieces w/ overlap” was set to 10, and the low-frequency rolloff sigma and cutoff radius were set to 0.03 and 0.1 pixel^−1^, respectively. A boundary model was created so that only the chromatin was enclosed. Fiducial patches were manually deleted if they overlapped with debris like ice crystals or if they mistracked. We did not detect a correlation between the alignment residual (Table S6) and our ability to detect canonical nucleosomes.

CTF estimation and phase flipping were done on the defocus phase contrast datasets using IMOD’s *ctfplotter* and *ctfphaseflip* programs. Prior to reconstruction, the cryosection and cryolamellae tilt series were binned to a final pixel size of 6.8 Å using the *eTomo* antialiasing option. Two tomogram versions were reconstructed for each tilt series. For visualization purposes, the tilt series were low-pass filtered to attenuate spatial frequencies beyond 25 Å to 30 Å resolution, prior to tomogram reconstruction. For classification analysis, the tilt series were low-pass filtered with a Gaussian rolloff starting at 15 Å resolution for lysates and 20 Å for cryosections and cryolamellae. In the classification jobs, the resolution of the data was further limited to 25 Å or 20 Å (see next section for details). More details of the datasets analyzed in this paper are shown in Table S6.

### Nucleosome template matching, classification, and subtomogram averaging

A featureless round-edged 10 nm-diameter, 6 nm-thick cylindrical template was created using the Bsoft program *beditimg* (Heymann and Belnap, 2007). A cubic search grid with a 12-nm spacing was created with the PEET program *gridInit* (Heumann, 2016). Regions that had high-contrast artifacts from surface contaminants and ice crystals were excluded. Template matching was done using PEET (Heumann, 2016; Heumann et al., 2011; Nicastro et al., 2006), with a duplicate-removal cutoff distance of 6 nm. To accelerate the runs, no orientation search was done around the cylindrical axis and the resolution was attenuated starting at 70 Å on account of the smooth appearance of the template. Candidate hit lists of different cross-correlation cutoffs were generated using the PEET program *createAlignedModel*, then visualized together with the tomograms in *3dmod*. The cross-correlation cutoff that eliminated spurious densities (primarily empty nucleoplasm) was chosen. The final numbers of subtomograms analyzed for each sample are in Table S7 and in the figures and figure legends.

Classification and subtomogram analysis were done with RELION (Kimanius et al., 2016; Scheres, 2012), following the workflows in Figure S1. In the published workflow (Figure S1A) (Bharat and Scheres, 2016), each subtomogram is first averaged by projecting the entire volume (16 nm along the Z axis), which introduces contributions from densities above and below the candidate nucleosome. To minimize the influence of other densities, pseudo-projections were created by averaging ∼12 nm along the Z axis, using the *ot_relion_project.py* script. 2-D classification using mask diameters ranging from 120 Å to 140 Å produced clear nucleosome-like classes, though the smaller masks included fewer adjacent densities. Note that in RELION, only circular masks are available for 2-D classification, meaning that it is not possible for the pseudo-projected densities to appear cylinder-like due to truncation by the mask. Densities that belonged to the most nucleosome-like classes were exported for 3-D classification, split into 30 classes. The resolution cutoffs were 25 Å for 2-D and 20 Å for 3-D classification. To eliminate the influence of adjacent densities during 3-D classification, a smooth cylindrical mask with a cosine-shaped edge was applied. The mask was created using *beditimg* and *relion_mask_create*. Because the GFP densities protrude from the nucleosome surface, we used a 9 nm-tall cylindrical mask for the analysis of nucleosomes with GFP tags, such as LGY0016 nucleosomes. The percentage of subtomograms belonging to each class was extracted with the script *count_particles.awk* from Guillaume Gaullier (Gaullier, 2021).

As observed in our previous study (Cai et al., 2018a), some canonical nucleosomes were lost in the 2-D classification process. We therefore used the alternative workflow in which the template matching hits were directly classified in 3-D (Figure S1B). To accommodate the increased diversity of complexes (some of which would have been removed had 2-D classification been done), we used 40 classes for BY4741, LGY0015, and LGY0016 nuclear lysates and 100 classes for cell cryolamellae and nuclear lysates of strains in which H3 or H4 were GFP tagged, without prior 2-D classification. These jobs crashed frequently because the RELION memory usage scales up with the number of classes (Kimanius et al., 2016). We were able to eliminate the crash problem by using higher-memory GPUs and by decreasing the number of translational search steps from 5 to 3 or 4. The canonical nucleosome classes were subjected to “Gold-standard” 3-D refinement (Henderson et al., 2012). No map sharpening was applied. Subtomogram class-average volumes were visualized with UCSF Chimera (Pettersen et al., 2004).

### Biased-reference classification control

A 14 Å-resolution density map was simulated from the yeast nucleosome crystal structure PDB 1ID3 (White et al., 2001) using the Bsoft program *bgex*. To account for the artifacts associated with defocus phase contrast and CTF correction, a 6 µm underfocus was applied and then “corrected” for using the Bsoft program *bctf*. The map was also subjected to the 20 Å-resolution low-pass filter that was used on the tilt series using the IMOD program *mtffilter*. Template matching was done using this reference, including data to higher resolution (28 Å) than that for the less-biased search above and including a search around all Euler angles. The hits were subjected to 2-D classification to remove obvious non-nucleosomal densities. Next, the hits were 3-D classified using the simulated map as an initial alignment reference. To maximize the model bias, the template was only low-pass filtered to 20 Å-resolution instead of the recommended 60 Å (Bharat and Scheres, 2016).

### Estimation of nucleosomes sampled per cell cryolamella

First, the average concentration of nucleosomes in the chromatin was estimated. The absolute number of nucleosomes per cell determined from genomics is 60,000 (Oberbeckmann et al., 2019). Soft X-ray tomography measurements revealed that the average G1 nucleus volume is 2 µm^3^, of which 20% is nucleolus (Uchida et al., 2011). Accordingly, chromatin (the nuclear volume not taken by the nucleolus), which contains the vast majority of nucleosomes, occupies ∼1.6 µm^3^. These two experimental values give an average nucleosome concentration of 37,500 per µm^3^. Next, the nuclear volume sampled by subtomogram analysis was determined by first drawing one closed contour around the chromatin using 3dmod; this closed contour encloses only the portion of the tomogram that was analyzed by template matching. The volume of this closed contour, which is one-voxel thick, was extracted using the command: imodinfo -F model.mod This command outputs the quantity “Cylinder volume”, in cubic pixels (voxels). The total tomographic volume sampled (Table S8) was obtained by multiplying Cylinder volume, the voxel volume (0.31 nm^3^ for 0.68 nm pixel size), and the number of tomographic slices that contain chromatin. The BY4741 VPP tomograms summed to 0.67 µm^3^, which yields ∼25,000 nucleosomes. For the HeLa cell in (Cai et al., 2018a) (EMPIAR-10179), the nucleus volume analyzed was 0.12 µm^3^.

### Simulations of nucleosome tomographic slices

Atomic models of the nucleosome (PDB 1KX5) (Davey et al., 2002) were manually positioned in UCSF Chimera and edited to remove the N-terminal tails. A 3-D density map was calculated with the Bsoft program *bgex*. There is no software that simulates Volta contrast, so we approximated the Volta-induced phase shift by setting the amplitude contrast to 100% and the defocus to zero in the Bsoft program *bctf*, followed by “correction”, also done with *bctf*. A tilt series was calculated from the simulated map using the IMOD program *xyzproj*. The tilt series was then aligned by cross-correlation and then backprojected using IMOD’s *eTomo* workflow. Parameters such as pixel size, tilt range, and tilt angle were kept as close to the experimental ones as possible. Tomographic slices were made in IMOD slicer at the same thickness as for the real cryotomograms.

### Ribosome subtomogram analysis control

Ribosomes were analyzed using the same software packages as for nucleosomes, as detailed in the workflow in Figure S25. Candidate ribosomes were template matched in a single VPP tomogram (from Figure S11), using a 25-nm-diameter sphere as a reference. The candidate ribosomes were filtered by cross correlation so that obvious false positives (vacuum) were excluded, leaving 3,816 hits. These particles were subjected to direct 3-D classification with k = 10, resulting in six non-empty classes. Particles belonging to the two ribosome classes (1,150 total) were pooled and “gold-standard” refined, yielding a density map at 28 Å resolution (FSC = 0.143) / 33 Å resolution (FSC = 0.5). Density maps were simulated at 15 Å, 20 Å, and 30 Å resolution using the Bsoft program *bgex* and the yeast ribosome crystal structure (Ben-Shem et al., 2011). To approximate the use of the VPP, the amplitude contrast was set to 90% and the defocus to −1 µm.

### Materials, data, and code availability

All *S. cerevisiae* strains generated in this study are available upon request. A subtomogram average of a BY4741 canonical nucleosome *ex vivo*, a double-GFP tagged LGY0016 nucleosome *ex vivo*, and the two BY4741 canonical nucleosome classes *in situ* have been deposited at EMDB as entry EMD-31086. All raw cryo-ET data, reconstructed tomograms and BY4741 cryolamellae VPP *in situ* class averages have been deposited in EMPIAR under entry EMPIAR-10678. All auxiliary scripts have been deposited at GitHub (https://github.com/anaphaze/ot-tools) and are publicly available as of the date of publication. Any additional information required to reanalyze the data reported in this paper is available upon request.

---

**Figure S1.**
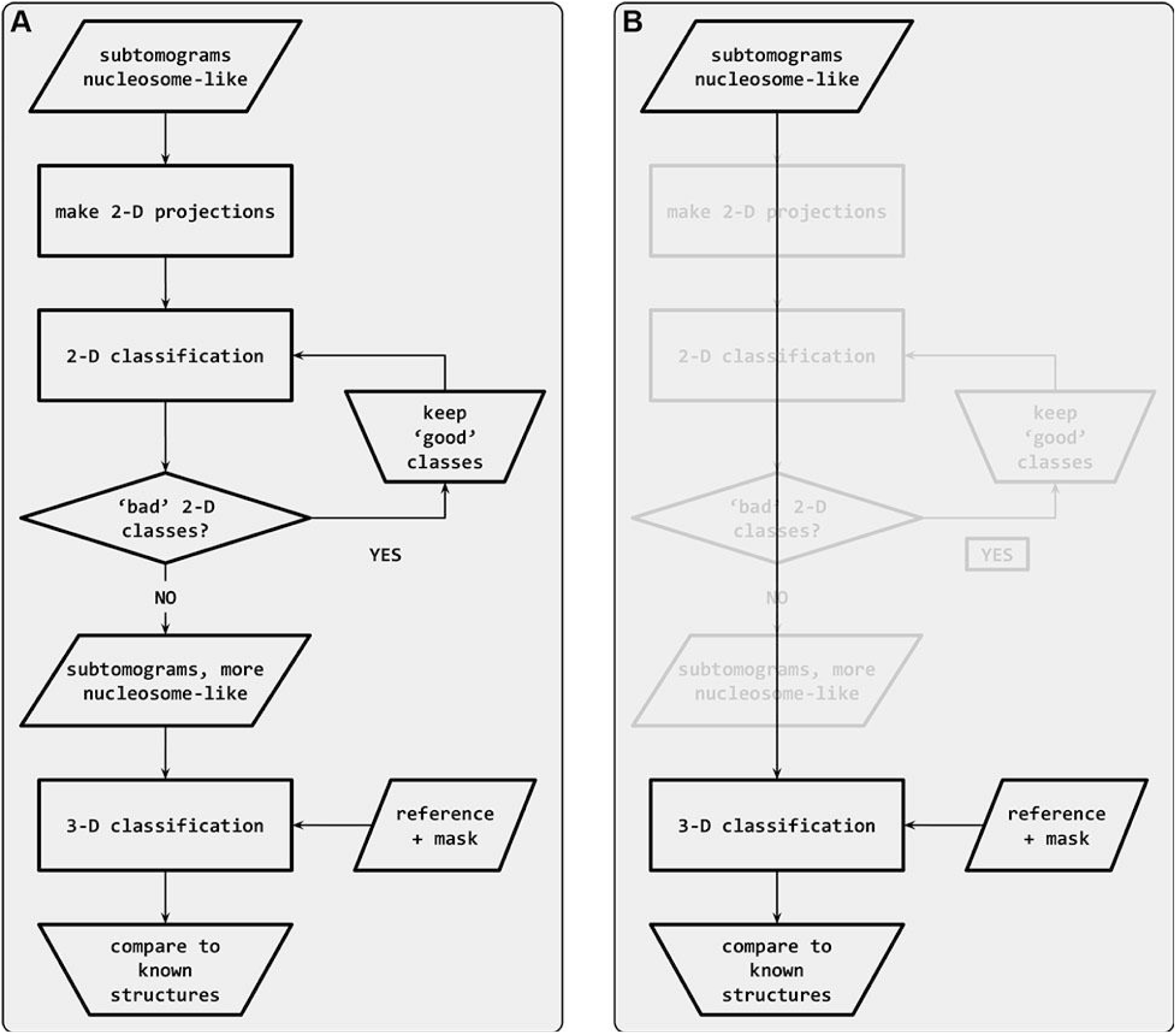
Subtomogram classification workflow. Classification starts with subtomograms that contain nucleosome-like particles, template matched in PEET (Cai et al., 2018). (A) In the workflow recommended by Bharat (Bharat et al., 2015), the subtomograms are first subjected to sequential rounds of 2-D classification to remove particles that belong to ‘bad’ classes. Once the bad classes are removed, the remaining set is subjected to 3-D classification. (B) In the alternative “direct 3-D” classification workflow, the subtomograms are subjected directly to 3-D classification. The 2-D classification steps (greyed out) are bypassed.

**Figure S2.**
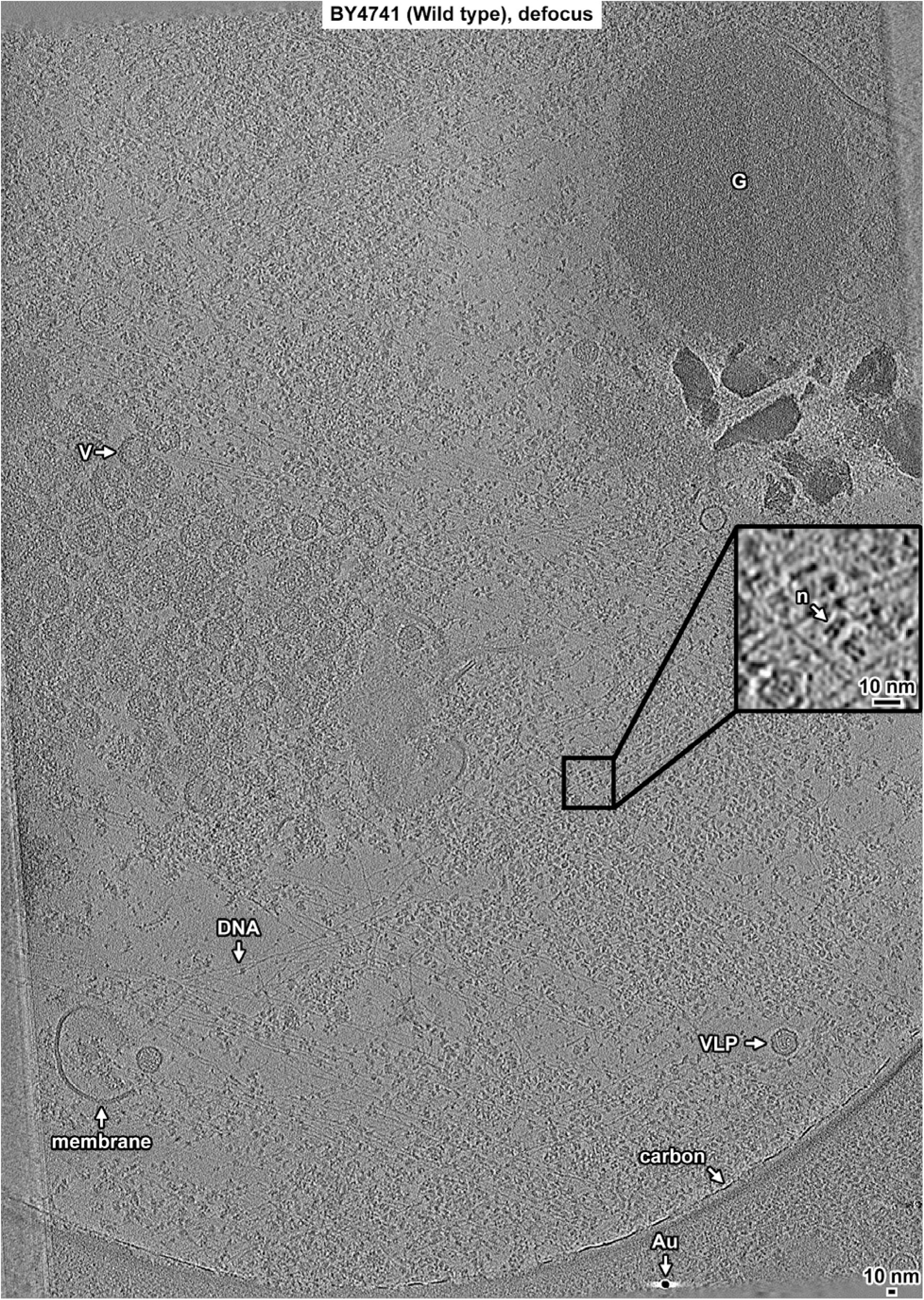
Overview of BY4741 (Wild type) nuclear lysate, defocus data. Tomographic slice (12 nm) of BY4741 nuclear lysates imaged with defocus phase contrast (defocus). Some non-chromatin features are indicated: granule (G), coated vesicle (V), naked DNA (DNA), membrane fragments (membrane), carbon support film (carbon), virus-like particle (VLP), and gold fiducial (Au). The abundant granular densities in this field of view are nucleosome-like particles. One example is indicated in the 4-fold enlargement in the inset (n). The linear features at the lower left and upper right are back-projection artifacts from the image borders. These regions are excluded from analysis.

**Figure S3.**
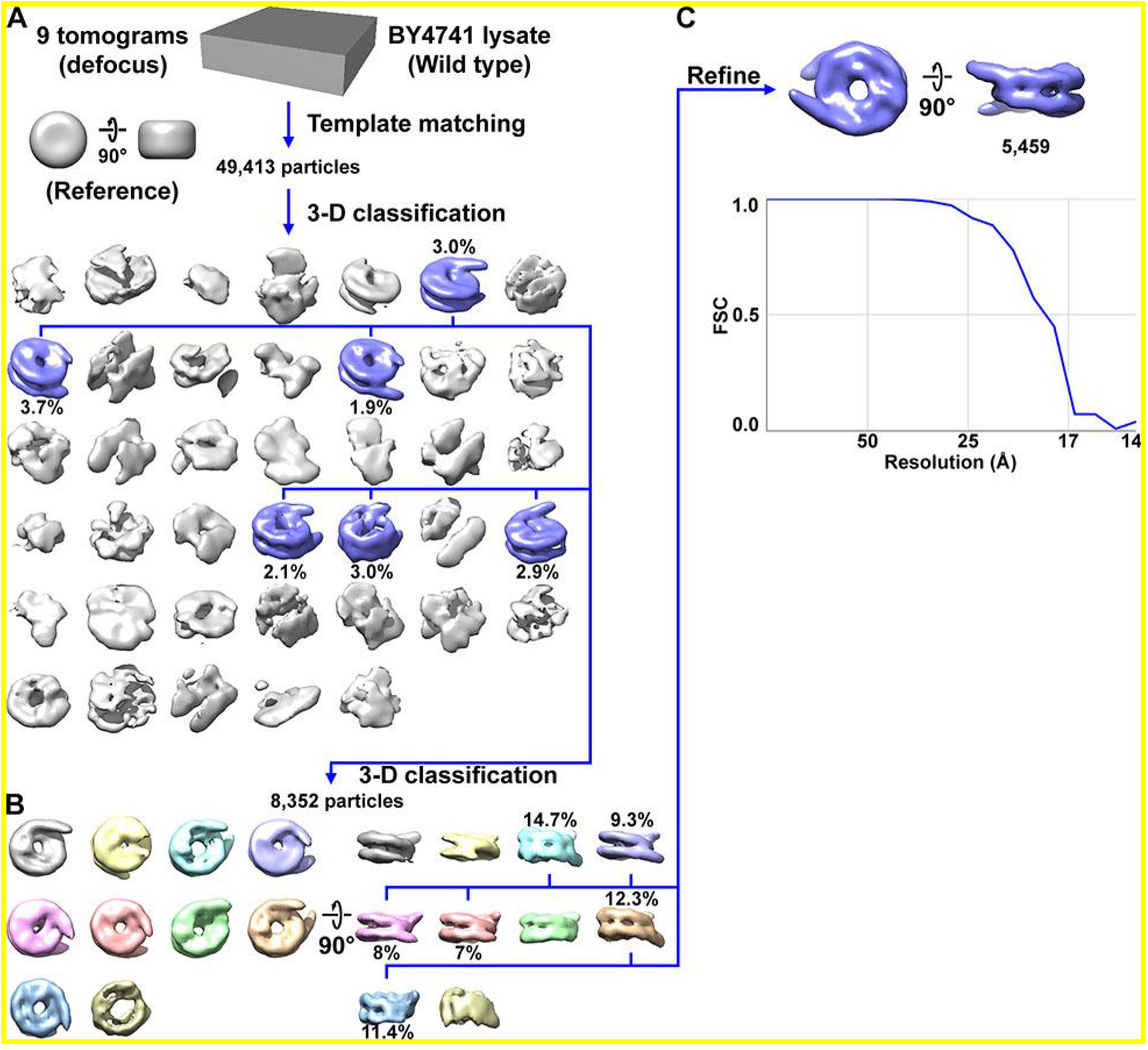
Direct 3-D classification of BY4741 (Wild type) nuclear lysates. (A) Class averages (3-D) of BY4741 candidate nucleosome template-matching hits. The canonical nucleosome class averages are shaded blue and the percent of particles belonging to the canonical classes are indicated under the density map. (B) Second round of classification, using the classes indicated in panel A. (C) The resolution of the refined canonical nucleosome is ∼18 Å by the FSC = 0.5 criterion. The refined density map is reproduced from Figure 1B.

**Figure S4.**
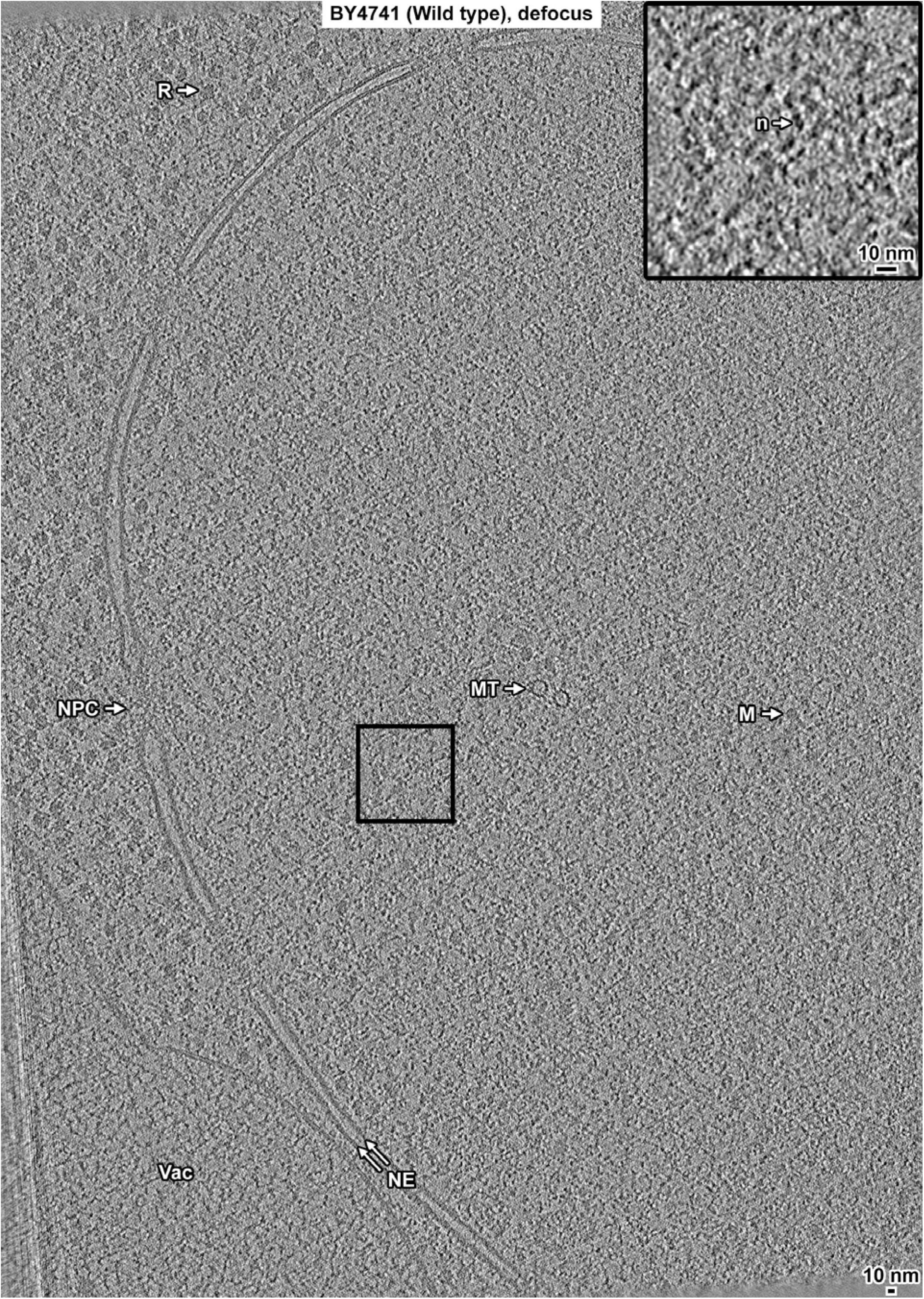
Overview of a BY4741 (Wild type) cell cryolamella, defocus data. Tomographic slice (12 nm) of a BY4741 cell cryolamella imaged with defocus phase contrast. Some non-chromatin features are indicated: nuclear envelope (NE), nuclear microtubule (MT), nucleosome-like particle (n), megacomplex (M), ribosome (R), and a putative vacuole (Vac). The high-contrast linear features at the lower left are back-projection artifacts from image borders.

**Figure S5.**
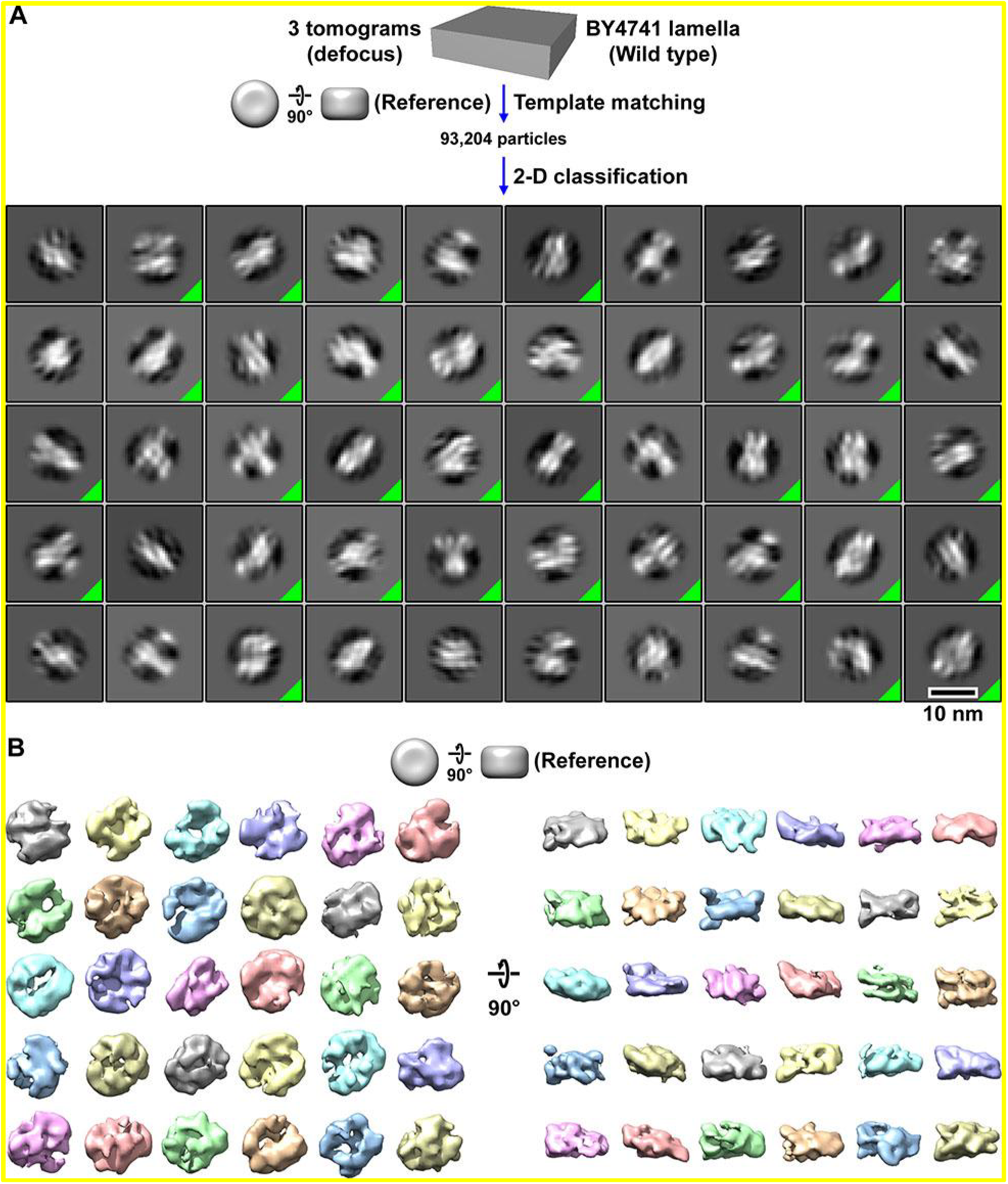
Classification of BY4741 (Wild type) nucleosome-like particles *in situ*. (A) Class averages (2-D) of template-matched nucleosome-like particles in BY4741 (wild-type) cell cryolamellae imaged by defocus phase contrast (defocus). The class averages whose member particles were selected for additional rounds of classification are indicated by a green triangle in the box’s lower right corner. The starting dataset consisted of 93,204 template-matched candidate BY4741 nucleosomes from 3 tomograms, from which 16,608 particles were selected by 2-D classification. (B) Class averages (3-D) of nucleosome-like particles in BY4741 cells. A featureless cylinder was used as the reference for template matching and 3-D classification. This workflow uses the same template-matched candidate nucleosomes as in Figure S6; see below.

**Figure S6.**
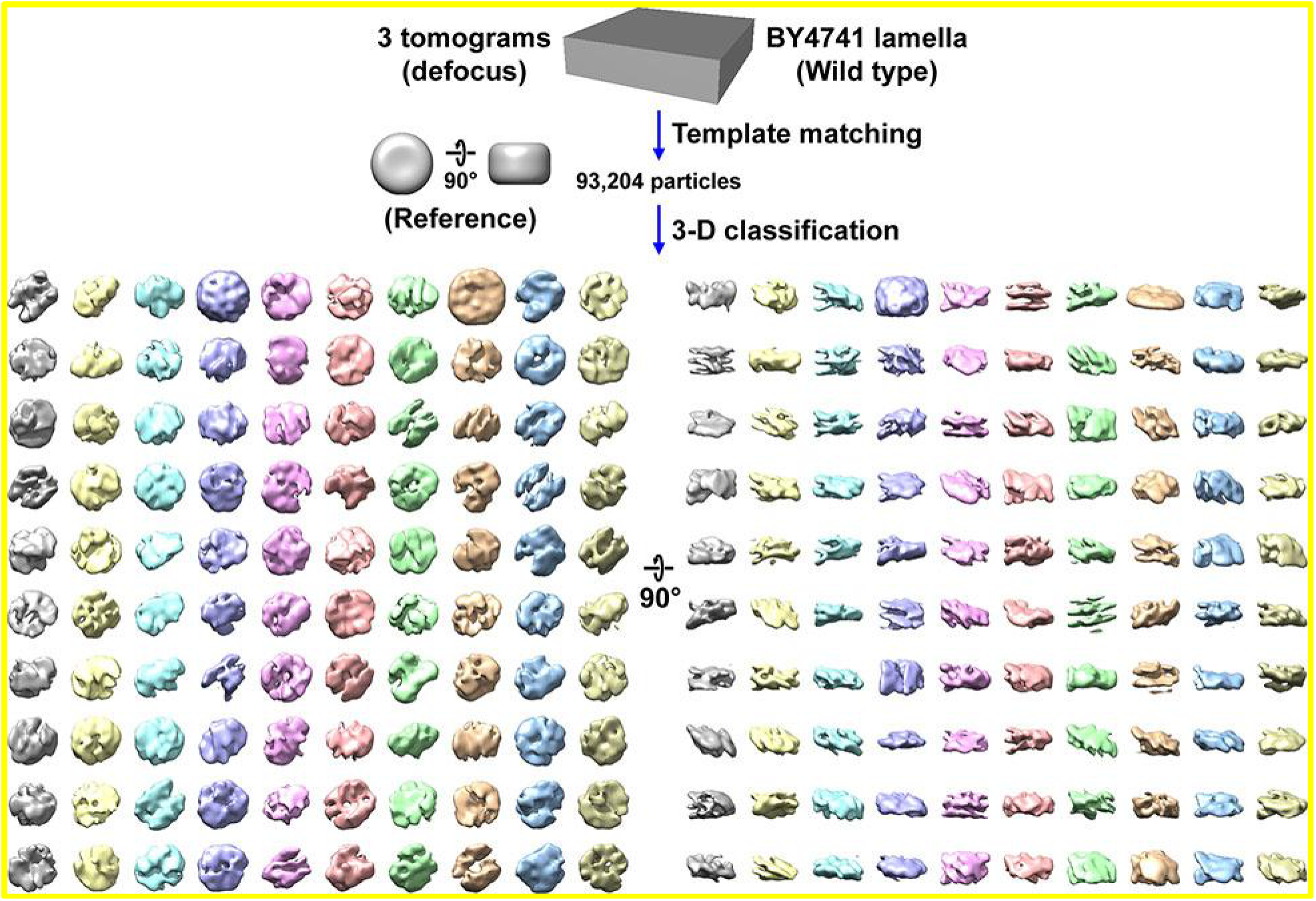
Direct 3-D classification of BY4741 (Wild type) cell cryolamellae densities. Class averages (3-D) of nucleosome-like particles in cryotomograms of BY4741 cell cryolamellae imaged by defocus phase contrast (defocus). In this experiment, 2-D classification was bypassed. This workflow uses the same template-matched candidate nucleosomes as in Figure S5.

**Figure S7.**
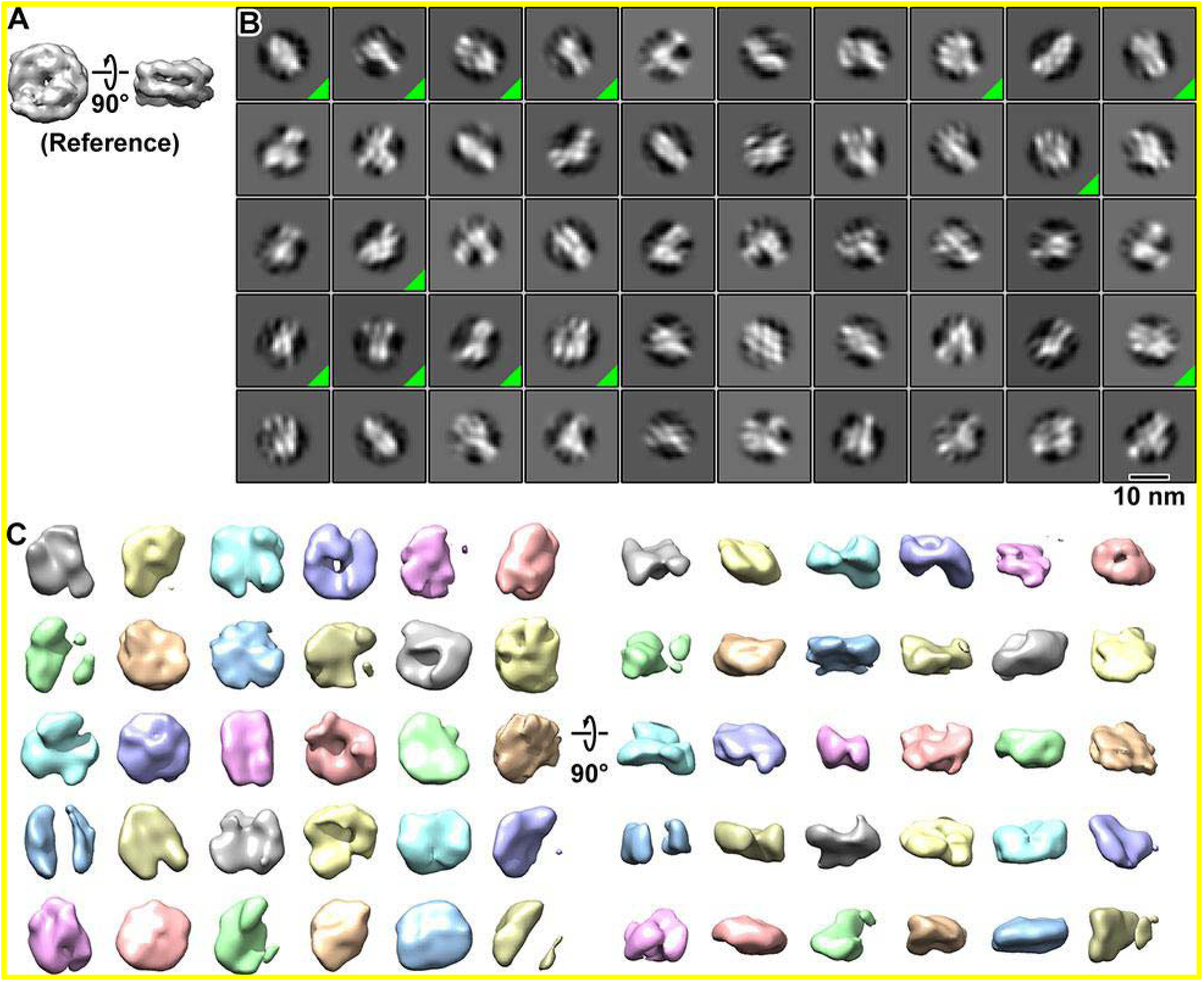
Classification using the nucleosome crystal structure reference. (A) A simulated EM map from a crystal structure of the nucleosome was used as the template-matching and 3-D classification reference. (B) Class averages (2-D) of BY4741 (wild type) nucleosome-like particles that were template matched using the nucleosome crystal structure. Classes that were pooled for 3-D classification are indicated by a green triangle in the lower right corner of its box. Out of 72,190 template-matched particles, 6,439 were selected by 2-D classification. (C) Class averages (3-D) of the most nucleosome-like particles selected by 2-D classification. The classification was performed using the same dataset as for Figures S5 and S6.

**Figure S8.**
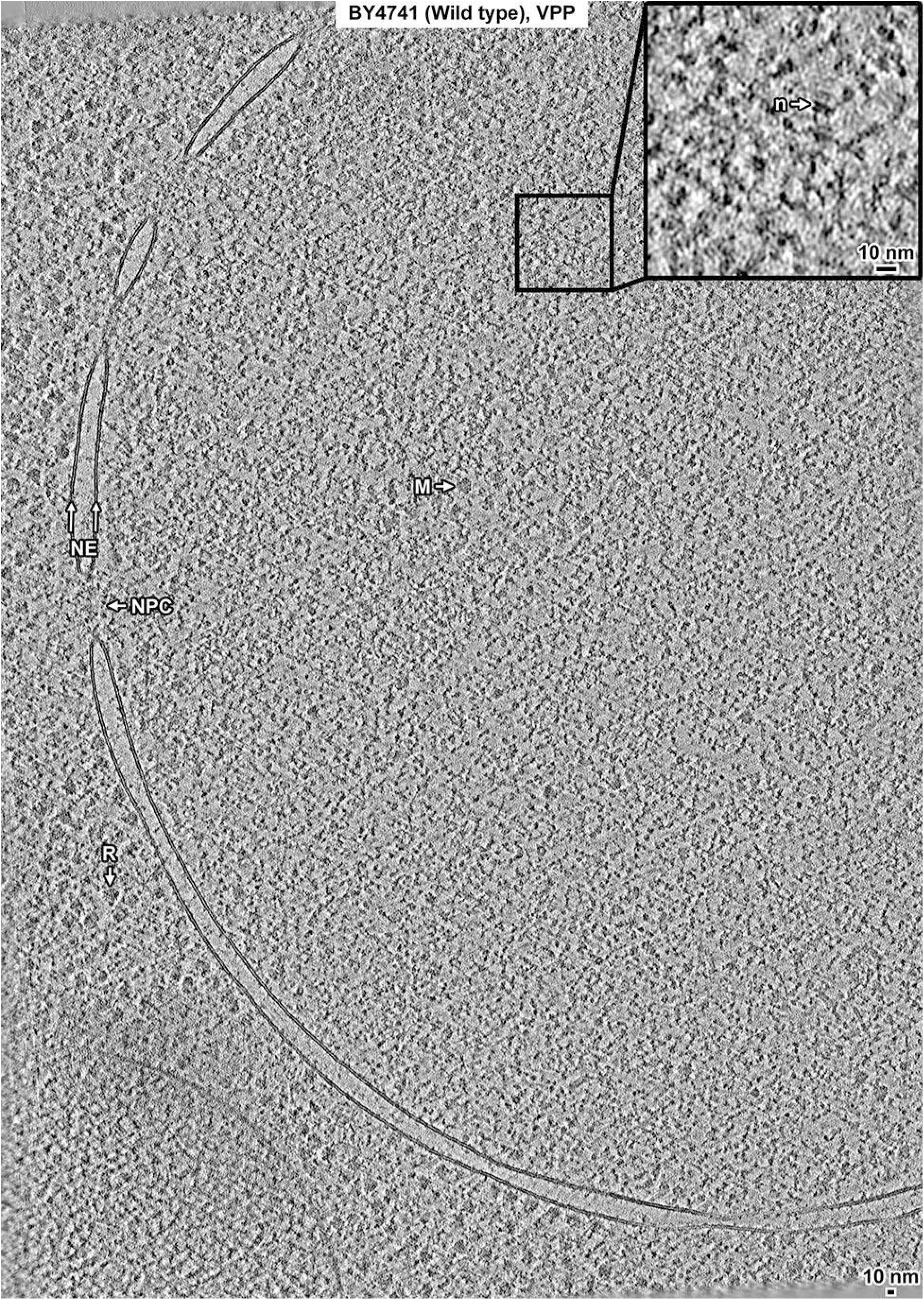
Overview of a BY4741 (Wild type) cell cryolamella, VPP data. Tomographic slice (12 nm) of a BY4741 cell cryolamella imaged with a Volta phase plate (VPP). The nuclear envelope (NE), nuclear pore complex (NPC), ribosome (R), and megacomplex (M) are indicated. The inset shows a threefold enlargement of the boxed area. A nucleosome-like complex (n) is indicated.

**Figure S9.**
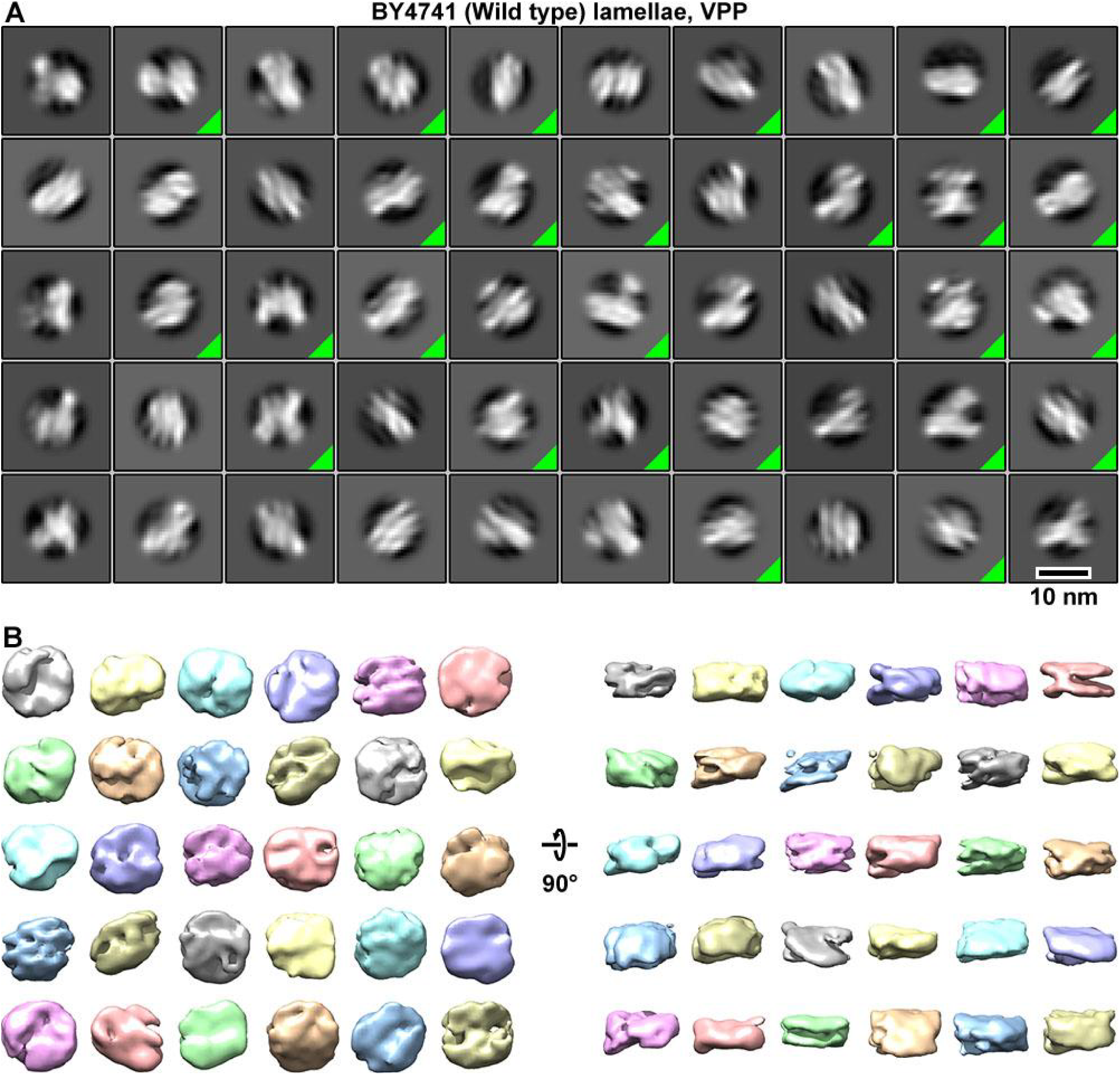
Classification of nucleosome-like particles *in situ* in BY4741 (Wild type) cell VPP data. (A) Class averages (2-D) of template-matched nucleosome-like particles in Volta phase plate (VPP) cryotomograms of BY4741 (wild type) cell cryolamellae. The class averages whose member particles were selected for additional rounds of classification are indicated by a green triangle in the box’s lower right corner. The starting dataset consisted of 129,473 template-matched candidate BY4741 nucleosomes from 3 tomograms. 12,551 particles were selected by 2-D classification. (B) Class averages (3-D) of nucleosome-like particles in BY4741 cells imaged with a VPP.

**Figure S10.**
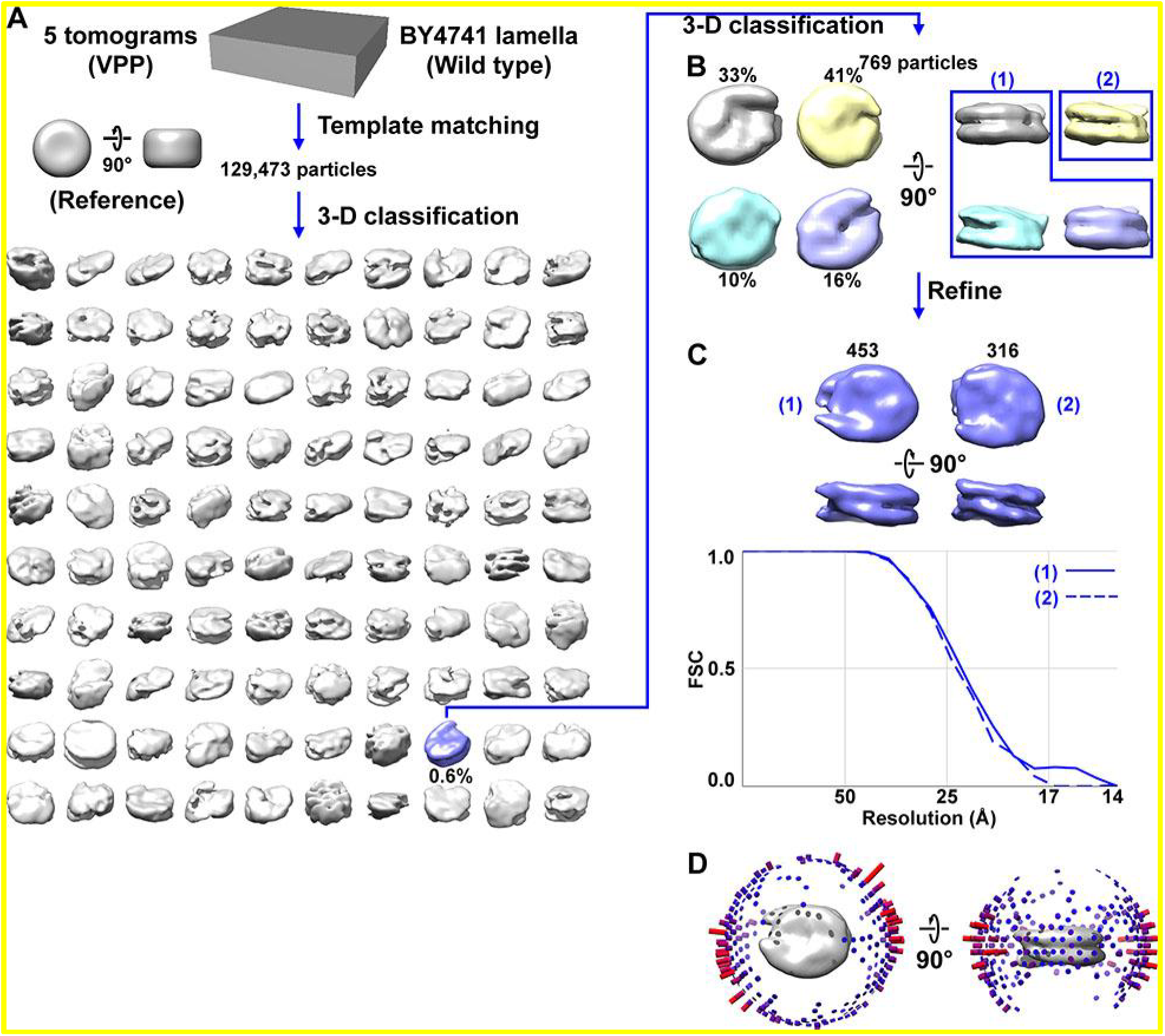
Direct 3-D classification of BY4741 (Wild type) nuclei in VPP tomograms of cryolamellae. (A) Class averages (3-D) of BY4741 nucleosomes from Volta phase plate (VPP) cryotomograms of cell cryolamellae. The canonical nucleosome class average is shaded blue and the percent of particles belonging is indicated under the density map. Note that this percentage is relative to the 129,473 template-matched nucleosome-like particles, not the ∼25,000 expected number of nucleosomes in the 5 tomograms. (B) Second round of classification from the canonical nucleosome class identified in panel A. (C) The resolution of both class averages is ∼24 Å by the FSC = 0.5 criterion. The refined density maps are reproduced from Figure 1C and the number of particles per class is labeled in black. (D) Angular-distribution plot of the canonical nucleosome class from panel A. Movies S1 and S2 show the progress of this classification jobs in panels A and B, respectively.

**Figure S11.**
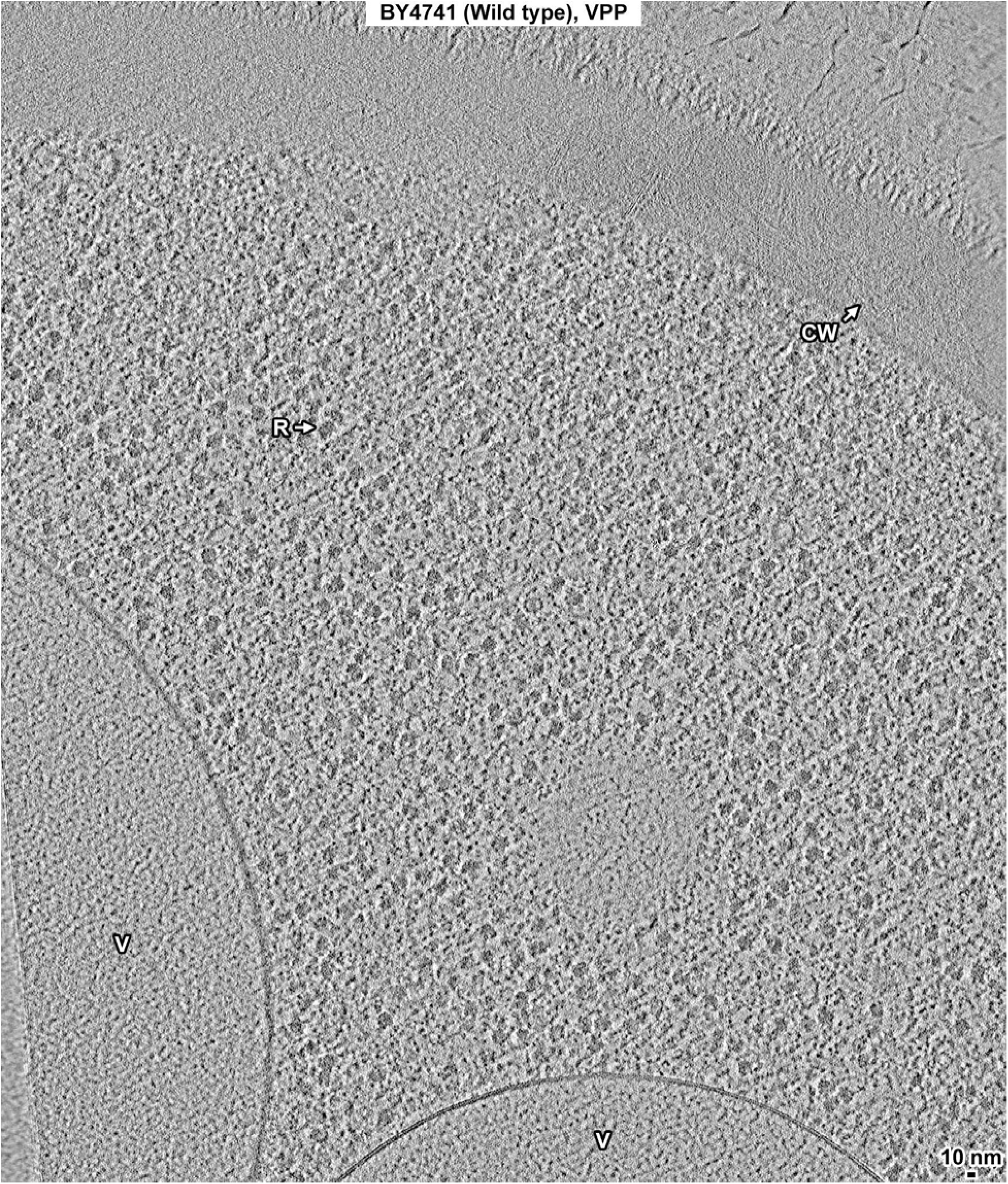
Overview of a BY4741 (Wild type) cell cryolamella imaged in the cytoplasm, VPP data. Tomographic slice (12 nm) of a BY4741 cell cryolamella imaged in the cytoplasm with a Volta phase plate (VPP). The vacuoles (V), a ribosome (R), and cell wall (CW) are indicated.

**Figure S12.**
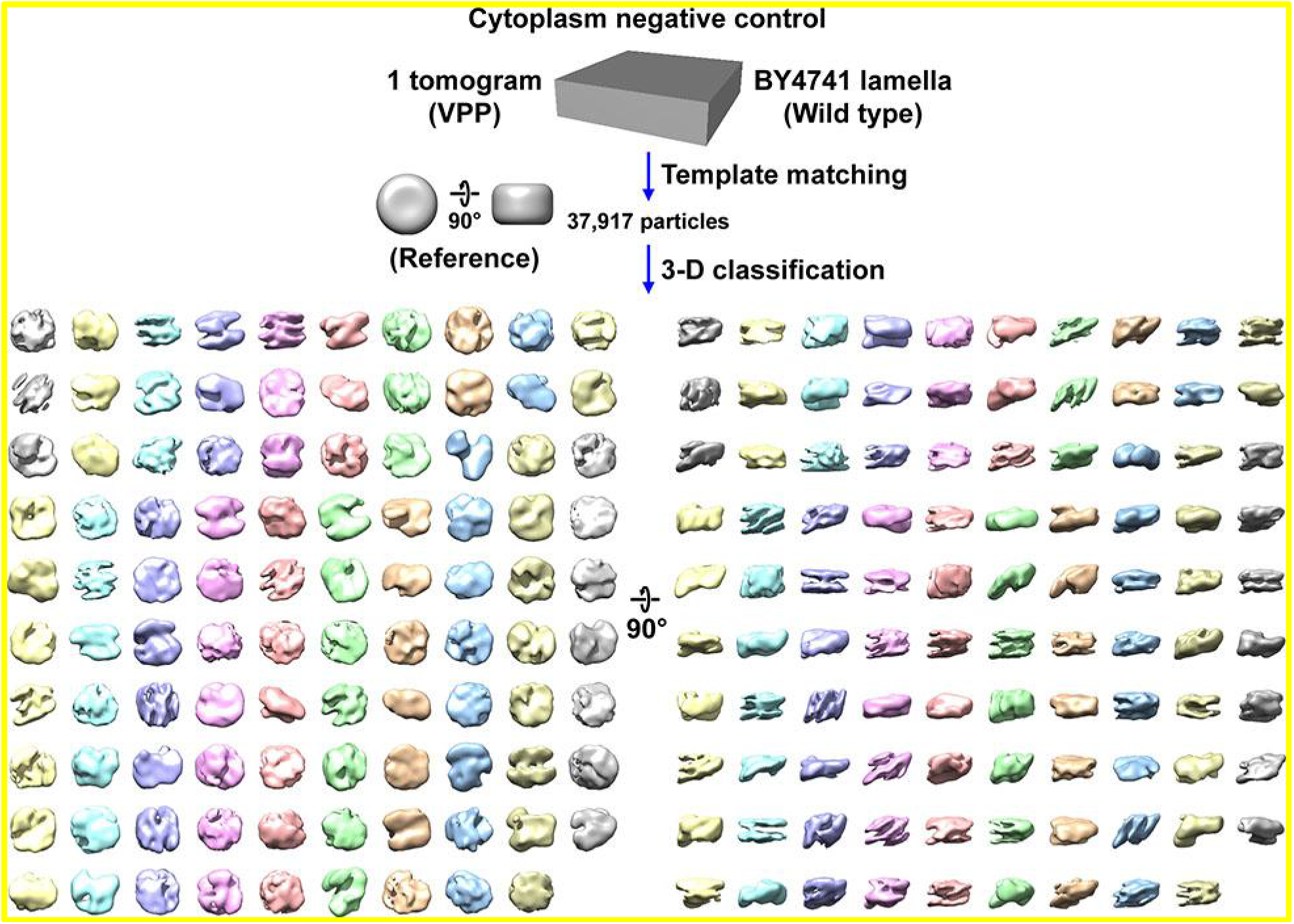
Classification of BY4741 (Wild type) cell cryolamellae VPP densities from the cytoplasm. Class averages (3-D) of “nucleosome-like” particles that were template matched from the cytoplasm in a BY4741 cell cryolamella imaged with a Volta phase plate (VPP). Both the template-matching and 3-D classification parameters were identical to the ones used to analyze the nucleus. 2-D classification was bypassed. One class has no contributing particles and was removed.

**Figure S13.**
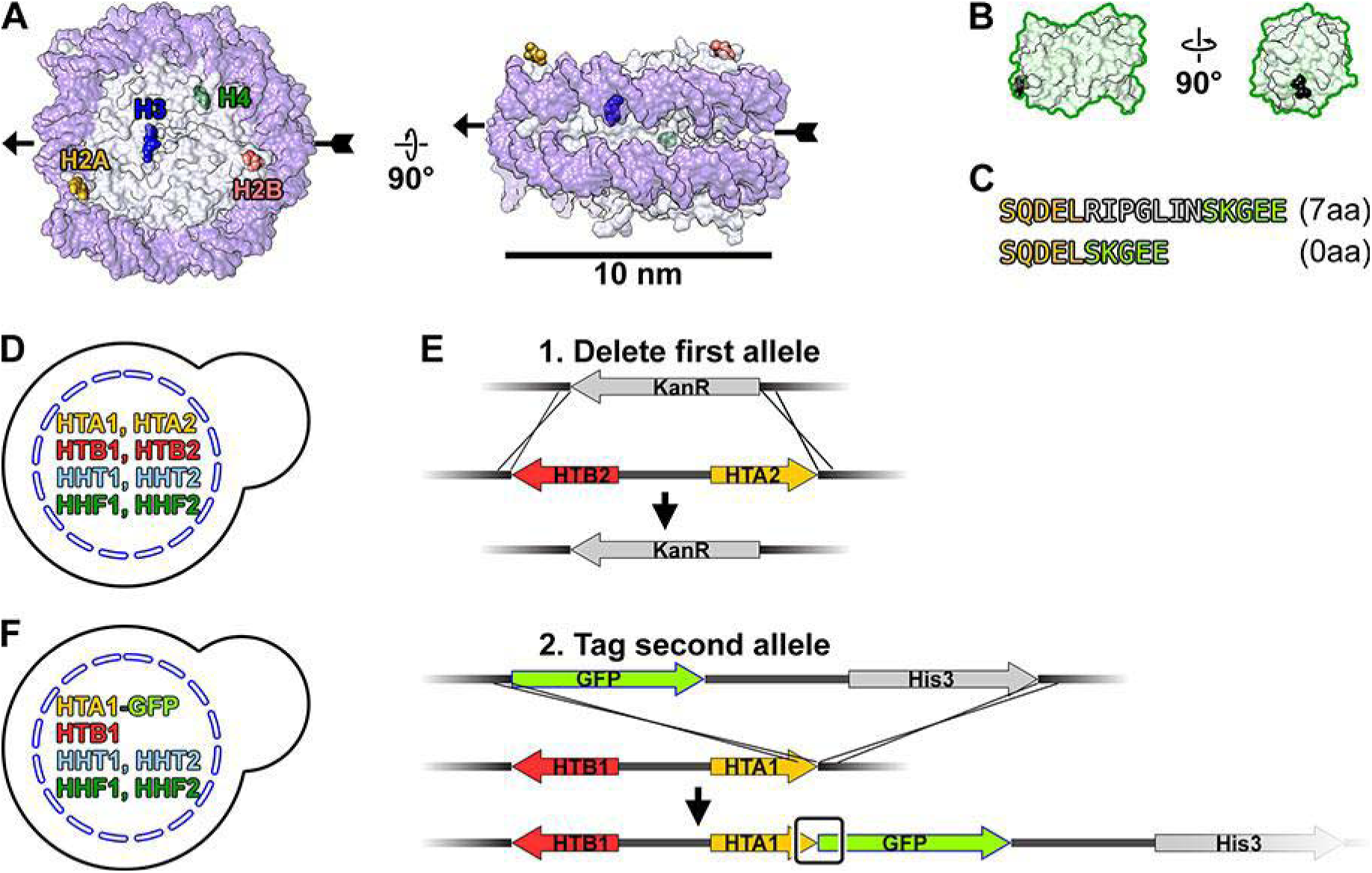
Strategy to tag H2A with GFP. (A) Crystal structure of the yeast nucleosome, PDB 1ID3 (White et al., 2001), in the disc view (left) and side view (right). DNA, dark purple; histone octamer, light gray. One set of histone C-termini are labeled and rendered as space-filling models. Two key points of reference, pseudo-dyad and opposite pseudo-dyad, are indicated by the arrowhead and arrow tail, respectively. The other set of C-termini is related by a 180° rotation around the pseudo-dyad axis. (B) Two views of the GFP crystal structure, PDB 1GFL (Yang et al., 1996), rendered at the same scale as panel A. The N-terminus is rendered as a black space-filling model. (C) Details of the sequence at the H2A-GFP fusion, with (7aa) and without (0aa) the 7 amino-acid linker. The H2A C-terminal sequence is shaded yellow, GFP N-terminal sequence is shaded green, and the flexible linker is shaded gray. LGY0016 has a 0aa linker. (D) Histone genotype of the parent strain BY4741. (E) Homologous recombination strategy to C-terminally tag the sole copy of the H2A gene with GFP. Upper – replacement of the HTA2-HTB2 locus with KanR. Lower – Tagging of HTA1 with GFP. The junction between the HTA1 (H2A) and GFP is boxed. (F) Histone genotype of LGY0016.

**Figure S14.**
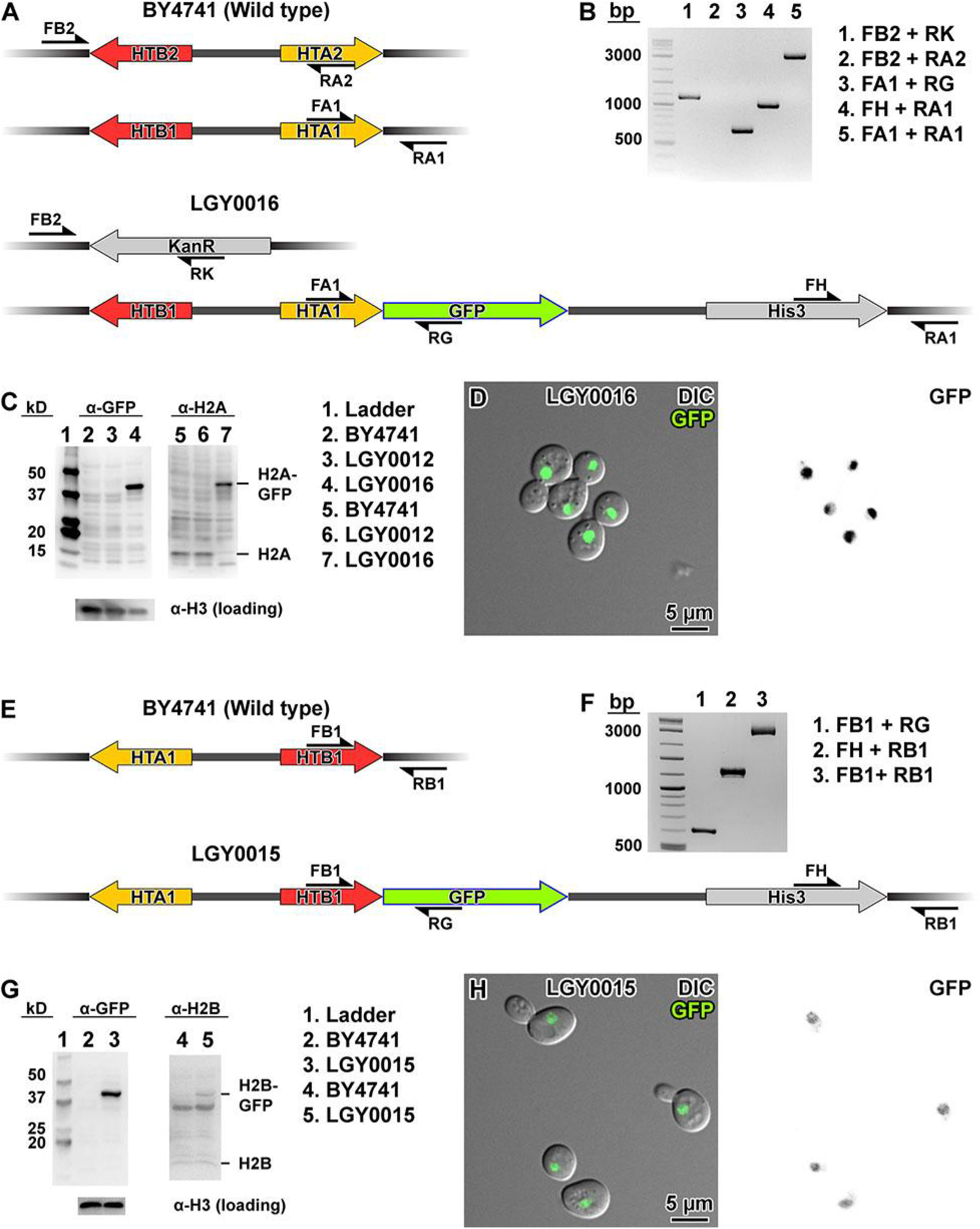
Experimental verification of H2A-GFP and H2B-GFP tagging. (A) Maps of the two H2A-H2B loci in the parent (Wild type) strain BY4741 and the H2A-GFP strain LGY0016. Primers used for PCR verification are indicated with the half arrow symbols. (B) Agarose gel of PCR amplicons expected (or not) from LGY0016 genomic DNA, in which the HTB2-HTA2 locus is deleted and the HTA1 locus is tagged with GFP. (C) Immunoblot analysis of strains LGY0016 (expresses H2A-GFP only) and LGY0012 (expresses H2A only, control). The α-GFP antibody correctly detected the large H2A-GFP fusion protein in LGY0016 (lane 4) but not in LGY0012 (lane 3, negative control). The α-H2A antibody detected H2A-GFP in LGY0016 and H2A in LGY0012. (D) DIC and GFP fluorescence confocal microscopy for LGY0016 cells. In the left panel, the GFP signals are overlaid in green. In the right panel, GFP signals are rendered with inverted contrast. (E) Map of the HTA1-HTB1 locus before (upper) and after (lower) the GFP insertion, with validation primers indicated. The color scheme is the same as in panel A. Note that LGY0015 still has the HTA2-HTB2 gene pair and therefore expresses untagged H2B. (F) Validation PCR for LG0015. (G) Validation immunoblots for strains BY4741 (wild type) and LGY0015 (expresses H2B and H2B-GFP). The α-GFP detected a ∼40 kDa protein (H2B-GFP) in LGY0015 but not BY4741. The α-H2B antibody detects the 40 kDa (H2B-GFP) in LGY0015 but not BY4741. Note that the α-H2B antibody has poorer specificity and does not generate a strong signal for the untagged H2B. (H) In the left panel, the GFP signals are overlaid in green. In the right panel, GFP signals are rendered with inverted contrast.

**Figure S15.**
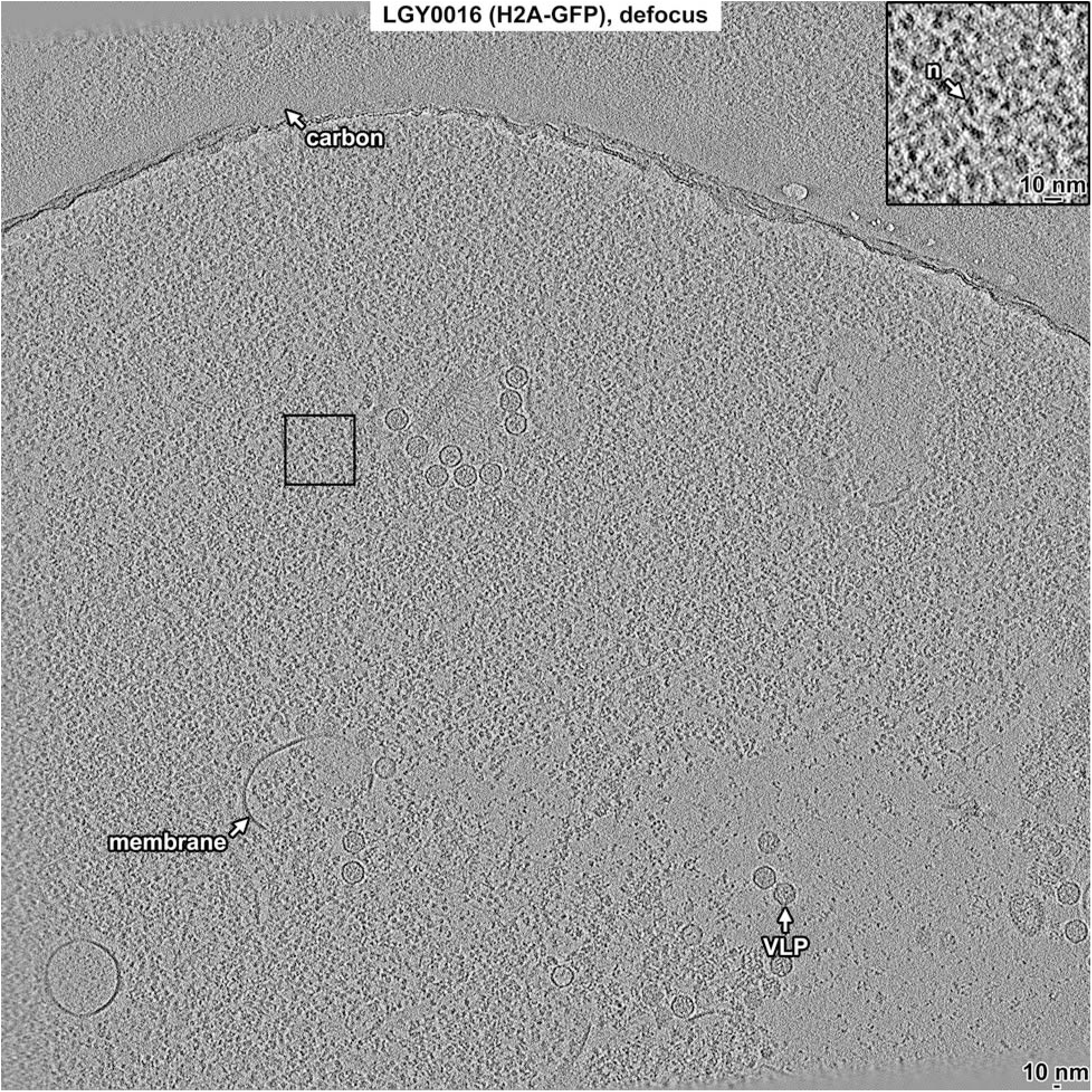
Overview of LGY0016 (H2A-GFP) nuclear lysate, defocus data. Tomographic slice (12 nm) of LGY0016 nuclear lysates imaged with defocus phase contrast (defocus). Some non-chromatin features are indicated: carbon support film (carbon), membrane fragments (membrane), gold fiducial (Au), and virus-like particle (VLP). The abundant granular densities in this field of view are nucleosomes. One small subarea (boxed) is enlarged 3-fold in the inset. A nucleosome-like particle (n) is indicated.

**Figure S16.**
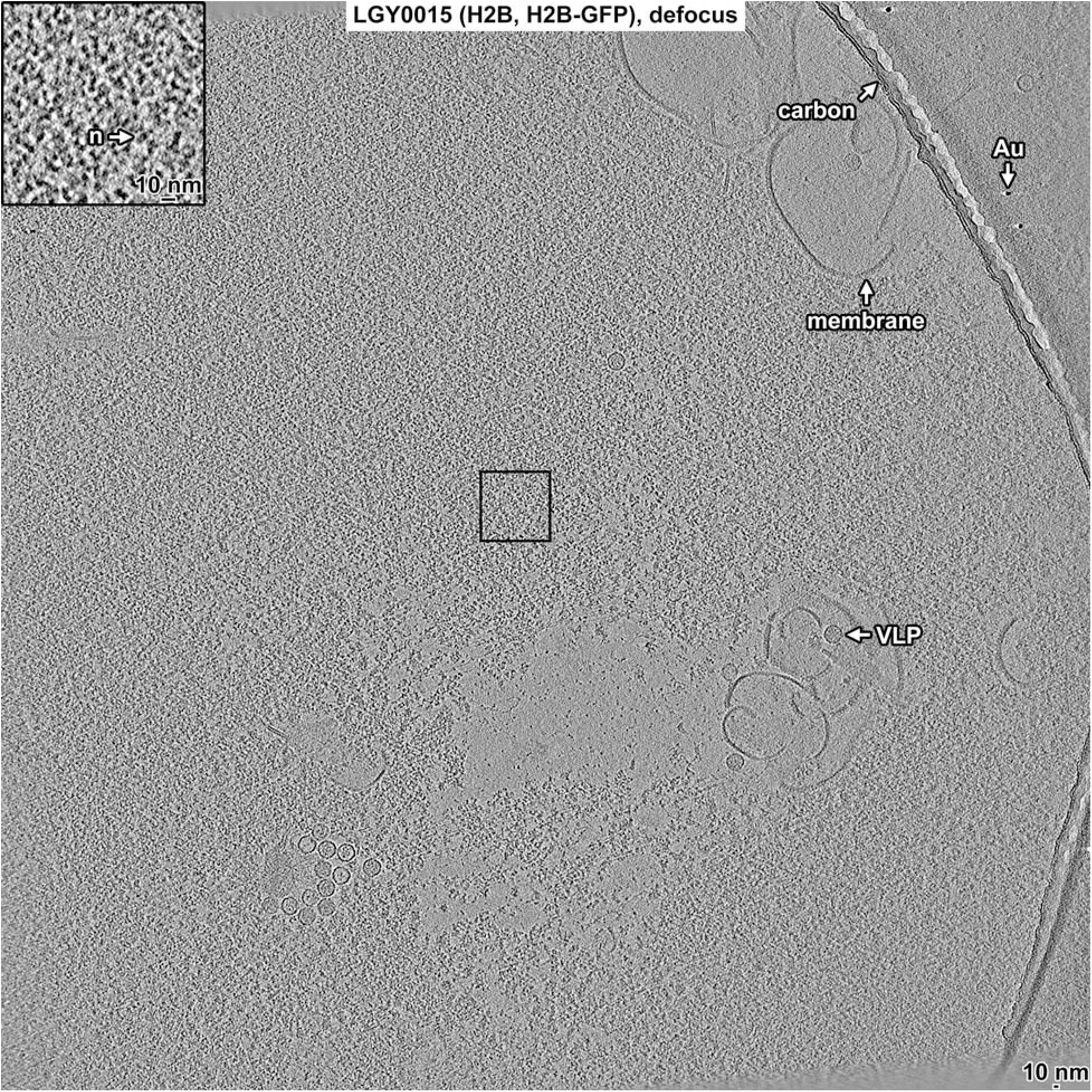
Overview of LGY0015 (H2B, H2B-GFP) nuclear lysate, defocus data. Tomographic slice (12 nm) of LGY0015 nuclear lysates imaged with defocus phase contrast (defocus). Some non-chromatin features are indicated: carbon support film (carbon), membrane fragments (membrane), gold fiducial (Au), and virus-like particle (VLP). The abundant granular densities in this field of view are nucleosomes. One small subarea (boxed) is enlarged 3-fold in the inset. A nucleosome-like particle (n) is indicated.

**Figure S17.**
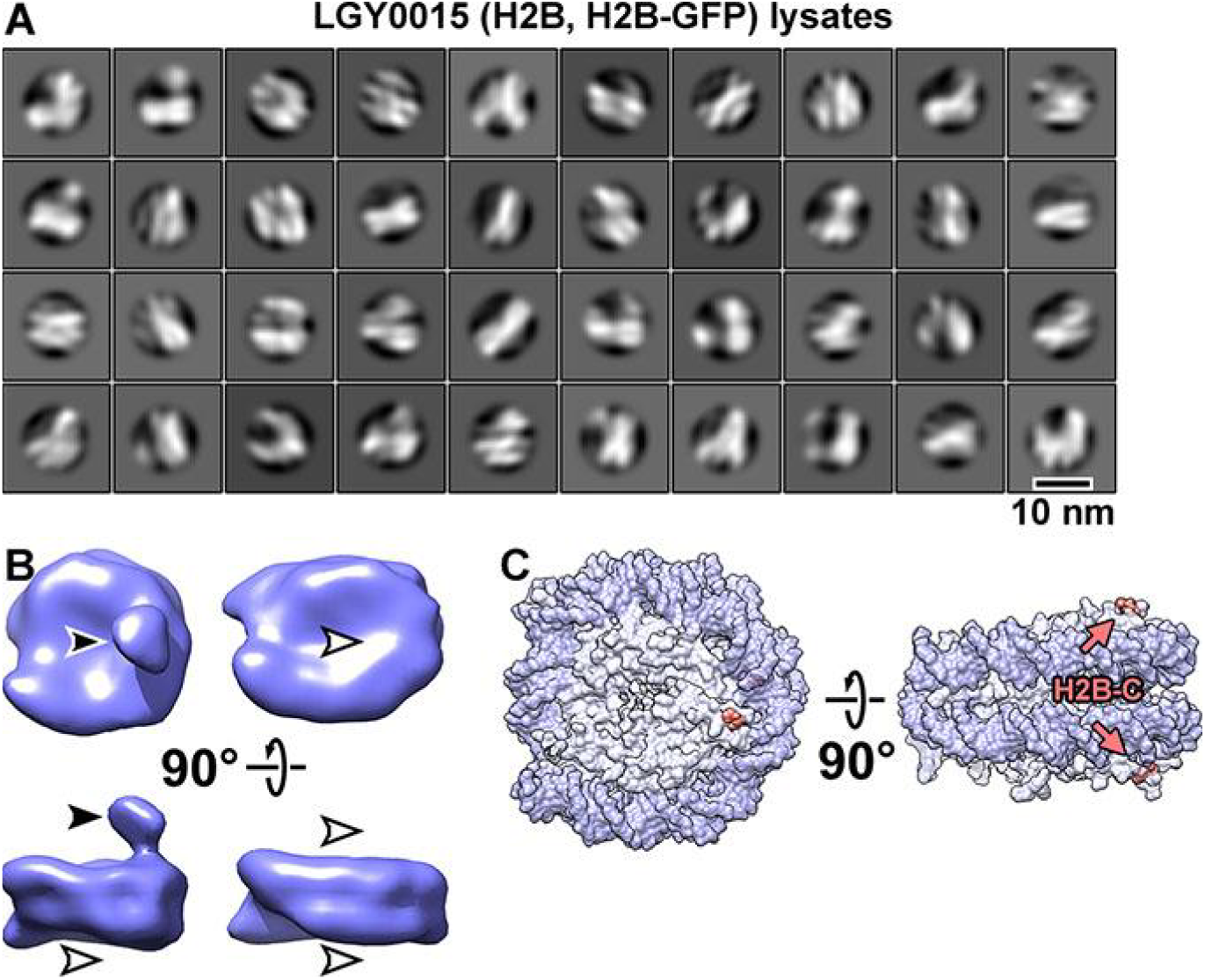
Classification of LGY0015 (H2B, H2B-GFP) nuclear lysates. (A) Class averages (2-D) of nucleosome-like particles. Out of 96,979 template-matched particles from 6 tomograms, 56,872 particles were selected by 2-D classification. (B) Class averages (3-D) of two types of LGY0015 nucleosomes. The solid arrowhead indicates the position of the GFP tag whereas the open arrowheads indicate the positions that lack this density. These are the same class averages shown in the overall workflow in Figure S19. (C) Two views of the nucleosome crystal structure, indicating the location of the H2B C-terminus (salmon).

**Figure S18.**
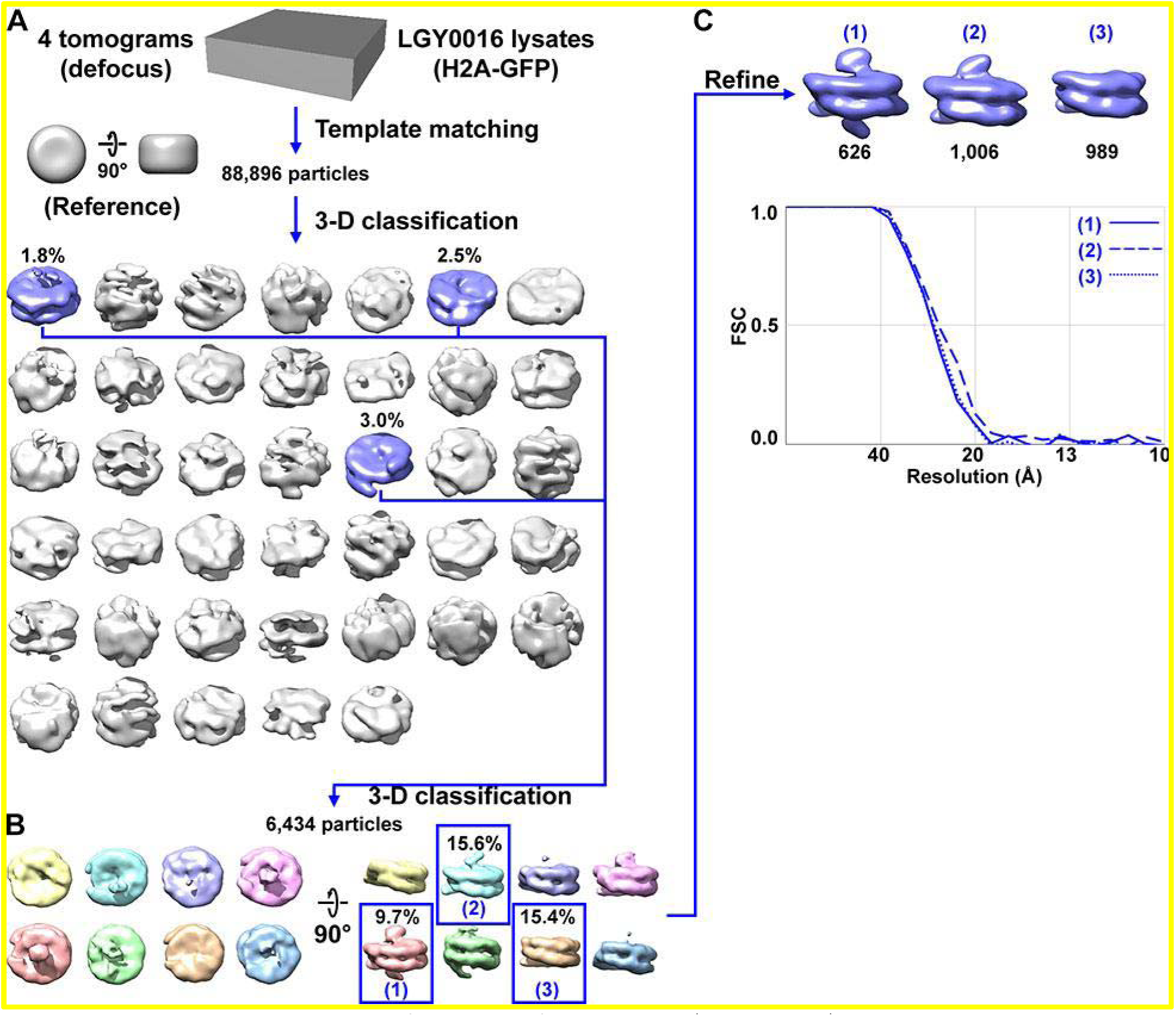
Direct 3-D classification of LGY0016 (H2A-GFP) nuclear lysates. (A) Class averages (3-D) of nucleosome-like particles from nuclear lysates of LGY0016 cells. The canonical nucleosome-like class averages are shaded blue while the non-canonical nucleosome averages are shaded gray. Movie S3 shows the convergence of the classification and shows more views of these class averages. (B) The second round of 3-D classification, using the canonical nucleosomes from panel A. See Movie S4 for more details of this classification job. (C) Refined densities of three types of LGY0016 nucleosomes isolated from nuclear lysates; reproduced from Figure 3B. The class numbers correspond to those in panel B (blue text) while the number of particles per class are labeled in black. The Fourier shell correlation (FSC) plot of the three refined class averages is labeled with the same numbering scheme. The resolution is ∼26 Å by the FSC = 0.5 criterion. Some of the class averages are “missing” one or both expected GFP densities. The possible explanations include mobility of a subpopulation of GFPs or H2A-GFPs, incorrectly folded GFPs, or substitution of H2A for the variant histone H2A.Z.

**Figure S19.**
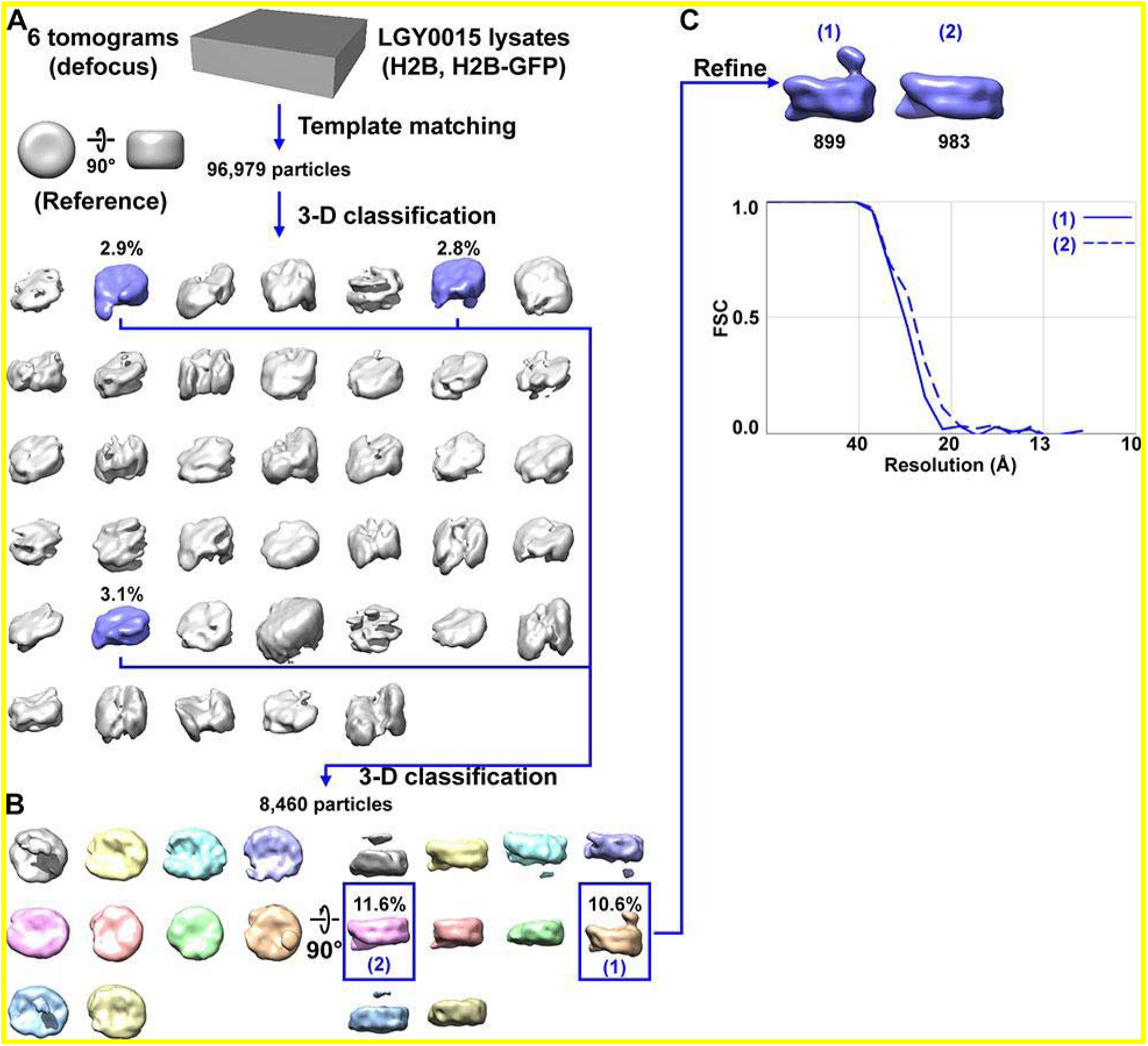
Direct 3-D classification of LGY0015 (H2B, H2B-GFP) nuclear lysates. (A) Class averages of nucleosome-like particles from nuclear lysates of LGY0015 cells. The canonical nucleosome class averages are shaded blue while the non-canonical nucleosome averages are shaded gray. (B) The second round of 3-D classification, using the canonical nucleosomes from panel A. Note that some class averages, such as the gray one, have a density that is not connected to the nucleosome. This density is from a nearby particle that protruded into the mask. (C) The two types of LGY0015 nucleosomes are reproduced from Figure S17B. The numbers of particles per class are indicated below the density maps in black. The FSC plot of the LGY0015 nuclear lysates nucleosome class averages in panel D uses the same numbering scheme. The resolution is ∼26 Å by the FSC = 0.5 criterion.

**Figure S20.**
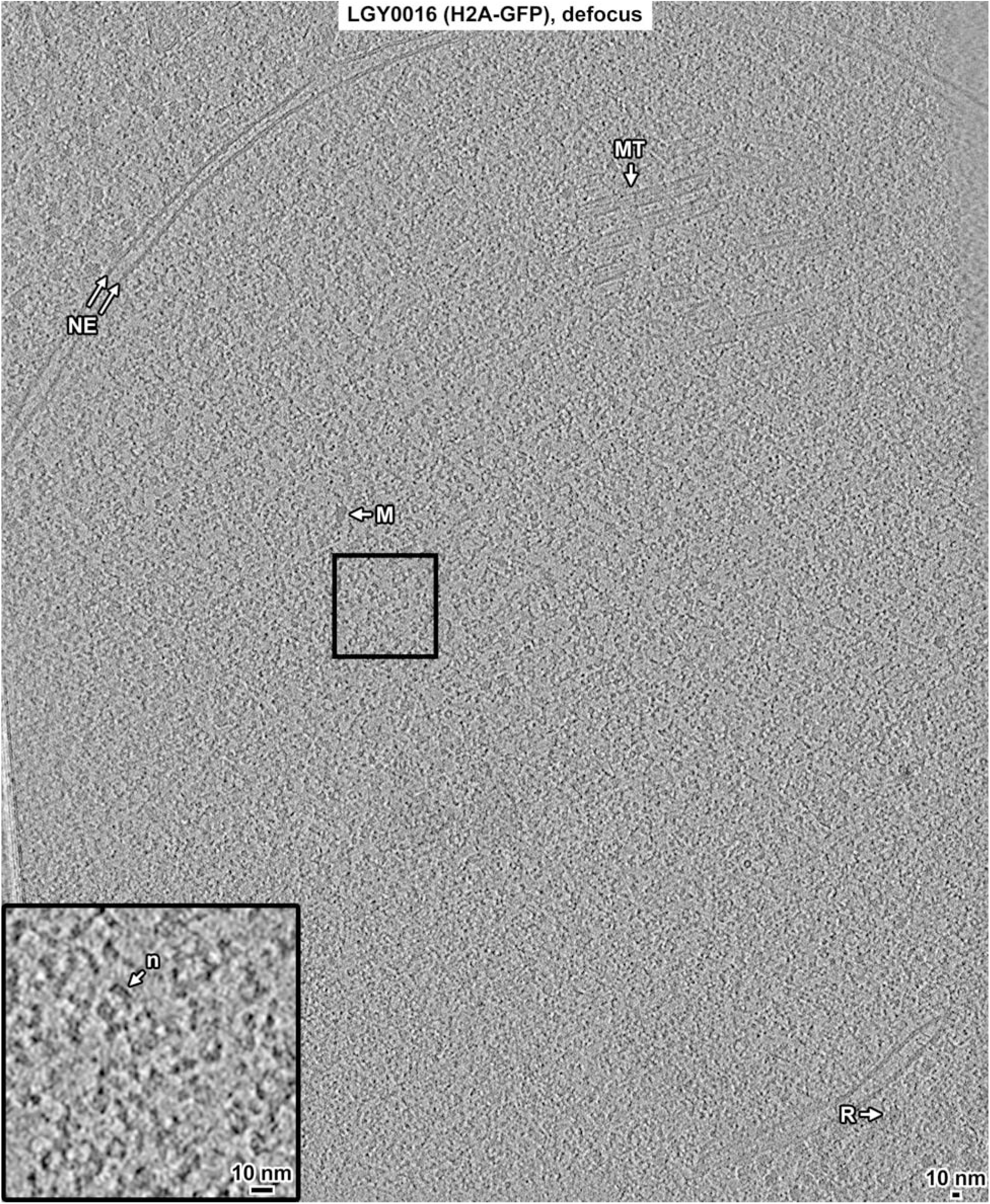
Overview of a LGY0016 (H2A-GFP) cell cryolamella, defocus data. Tomographic slice (12 nm) of a LGY0016 cryolamella, imaged with defocus phase contrast. Some non-chromatin features are indicated: nuclear envelope (NE), nuclear microtubule (MT), megacomplex (M), and ribosome (R). The inset is a threefold enlargement of the boxed area. A nucleosome-like particle (n) is indicated.

**Figure S21.**
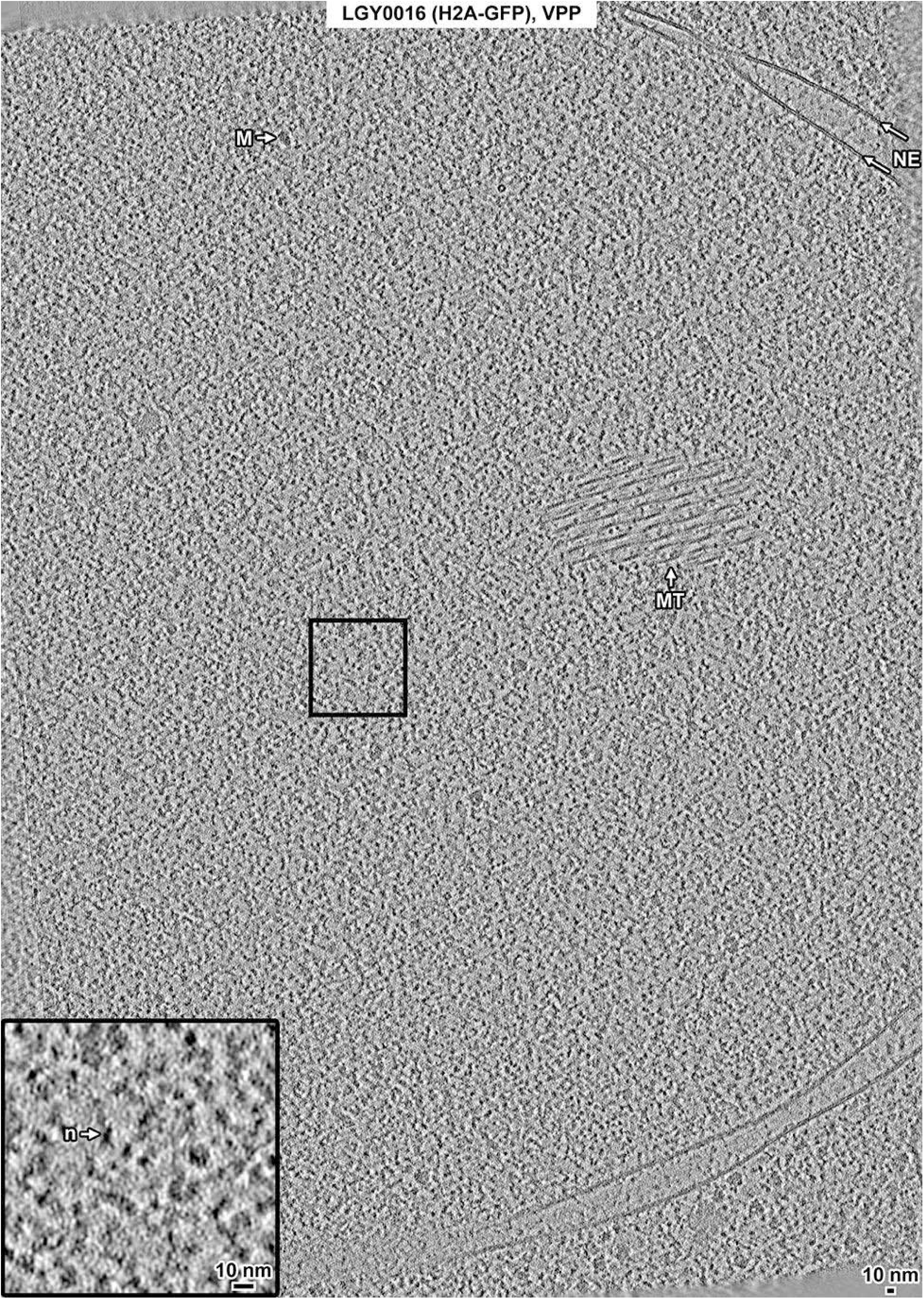
Overview of a LGY0016 (H2A-GFP) cell cryolamella, VPP data. Tomographic slice (12 nm) of a LGY0016 cryolamella imaged with a Volta phase plate (VPP). The nuclear envelope (NE), nuclear microtubule (MT), and a megacomplex (M) are indicated. The inset is a threefold enlargement of the boxed area. A nucleosome-like particle (n) is indicated.

**Figure S22.**
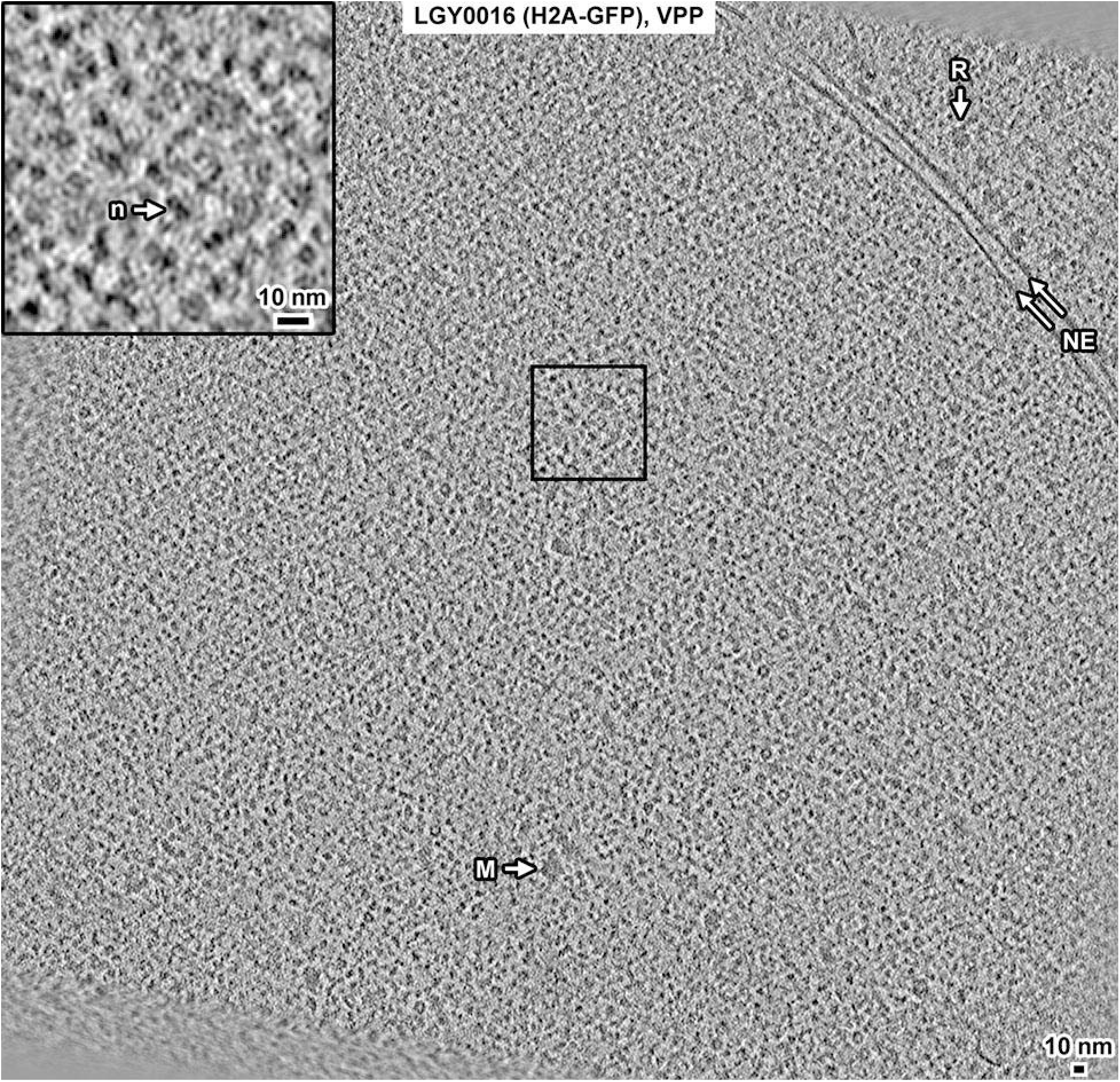
Overview of a LGY0016 (H2A-GFP) cell cryosection, VPP data. Tomographic slice (12 nm) of a LGY0016 cryosection imaged with a Volta phase plate (VPP). Some non-chromatin features are indicated: nuclear envelope (NE), megacomplex (M), and ribosome (R). The inset is a threefold enlargement of the boxed area. A nucleosome-like particle (n) is indicated. The smeared features at the lower left are back-projection artifacts from image borders.

**Figure S23.**
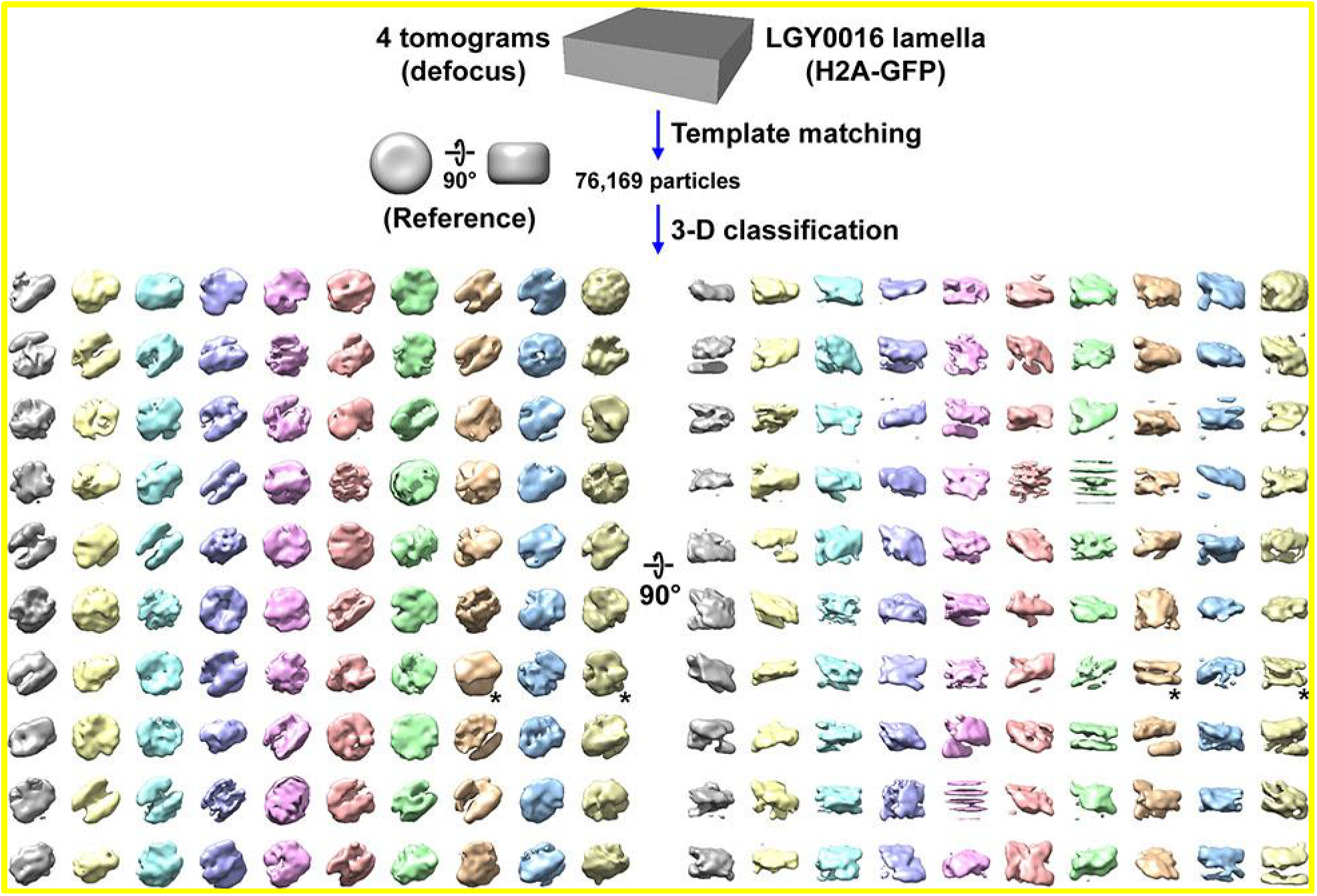
Direct 3-D classification of LGY0016 (H2A-GFP) cell cryolamellae densities. Class averages (3-D) of nucleosome-like particles in cryotomograms of LGY0016 cell cryolamellae imaged with defocus phase contrast (defocus). The starred classes have two linear motifs.

**Figure S24.**
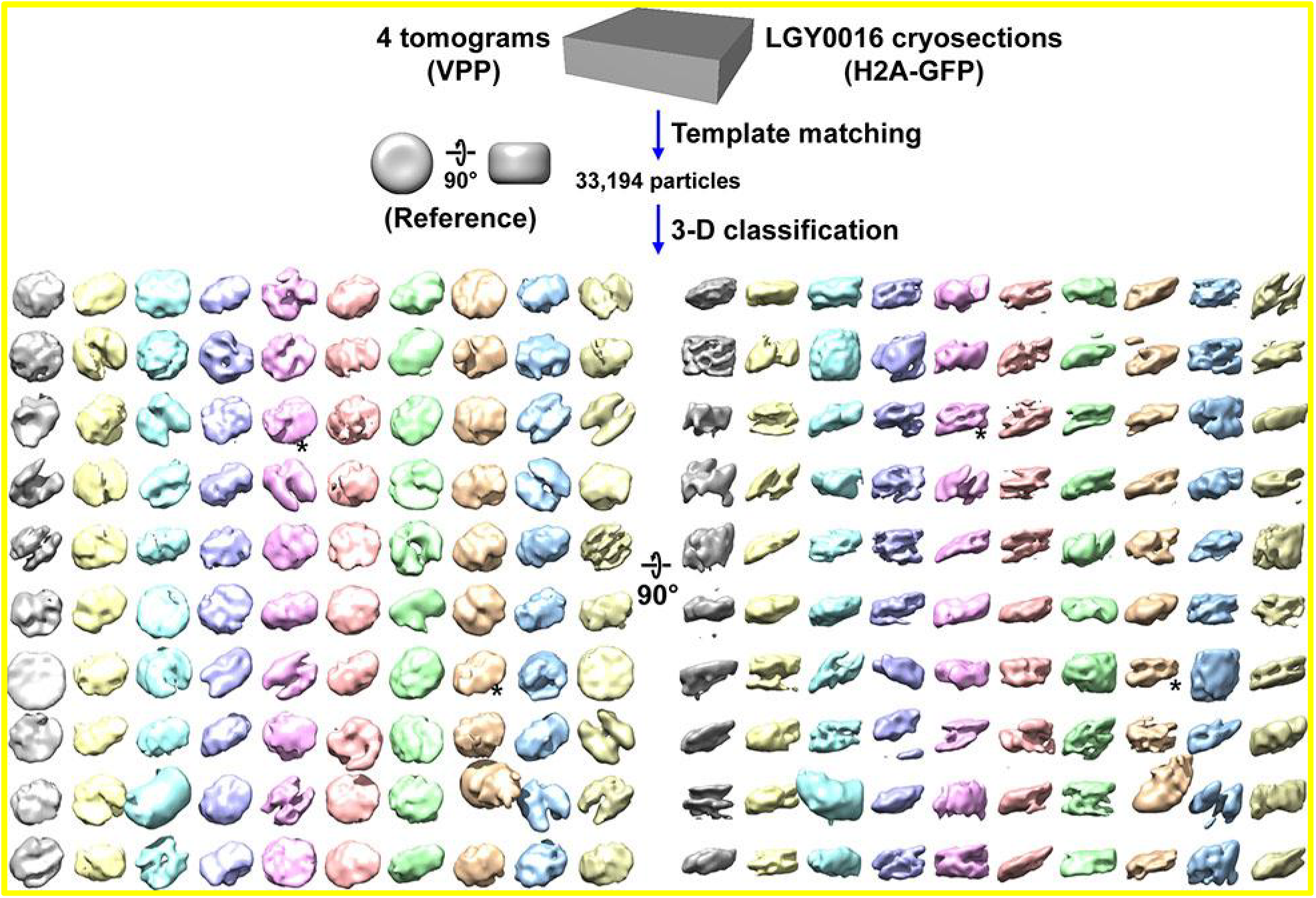
Direct 3-D classification of LGY0016 (H2A-GFP) cell cryosection densities in VPP data. Class averages (3-D) of nucleosome-like particles in VPP cryotomograms of LGY0016 cell cryosections imaged with a Volta phase plate (VPP). The starred classes have two linear motifs.

**Figure S25.**
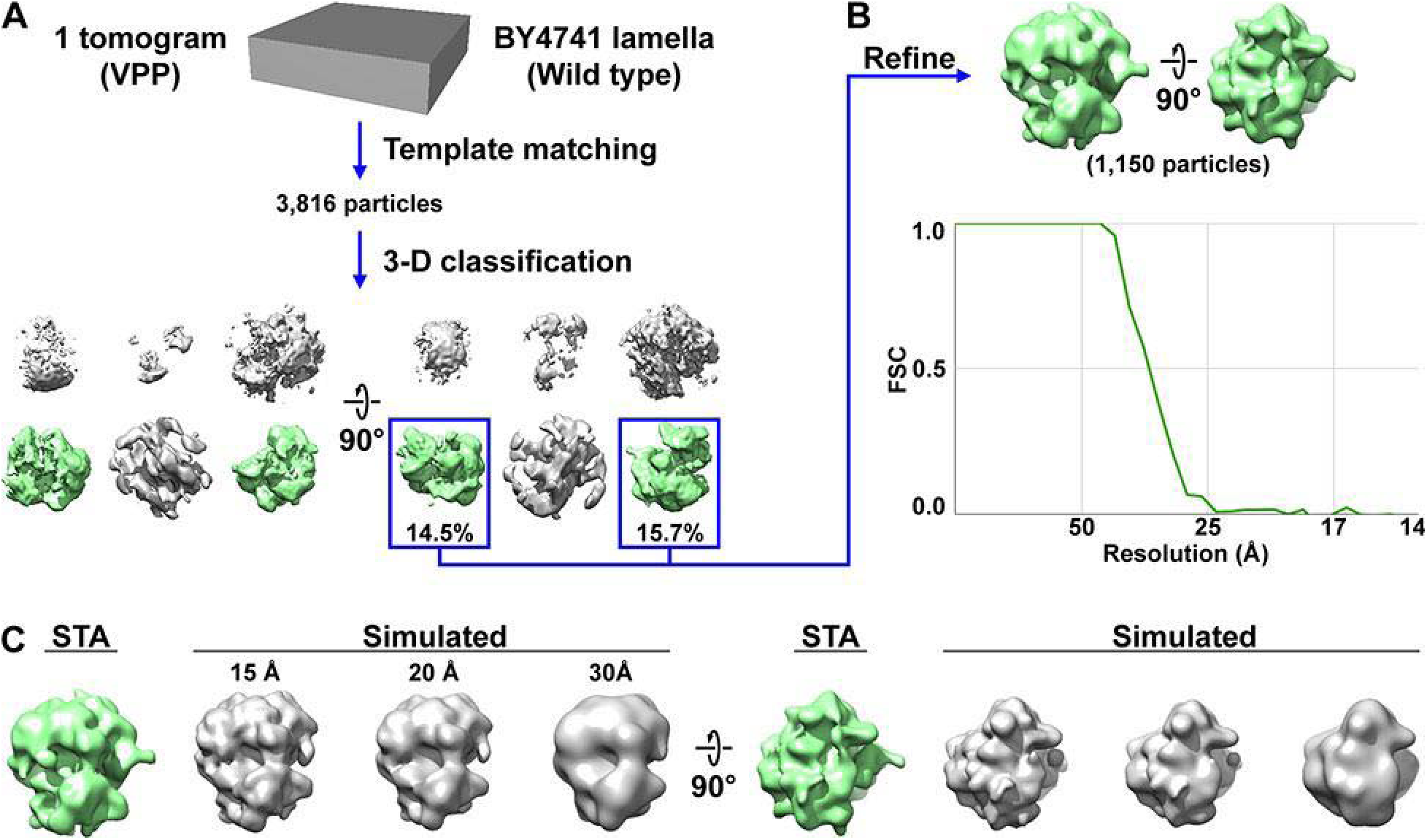
Control VPP subtomogram analysis of ribosomes *in situ*. (A) Candidate ribosomes were template matched from a single Volta phase plate (VPP) tomogram of the cytoplasm from Figure S11, using a 25-nm-diameter sphere as a reference. Direct 3-D classification into ten classes led to six non-empty classes, of which two were of ribosomes. Note that template matching is insufficient to uniquely identify a macromolecular complex, particularly when the reference is a featureless body like a sphere or cylinder. A simple reference was used to reduce the model bias. Template matching done with featureless references generates substantial numbers of false positives, which are confidently removed by classification. Complexes are only considered as identified after the 3-D classification step produces recognizable class averages. Therefore, the term “candidate ribosomes” is used to describe the 3,816 template matching hits. (B) The 1,150 ribosome subtomograms were pooled and 3-D refined, producing an average at 28 Å resolution by the “Gold-standard” FSC (0.143 cutoff) criterion and 33 Å resolution with an FSC = 0.5 cutoff. (C) Comparison of the *in situ* ribosome subtomogram average (STA) with density maps simulated from the yeast ribosome crystal structure (Ben-Shem et al., 2011) at three different resolutions.

**Figure S26.**
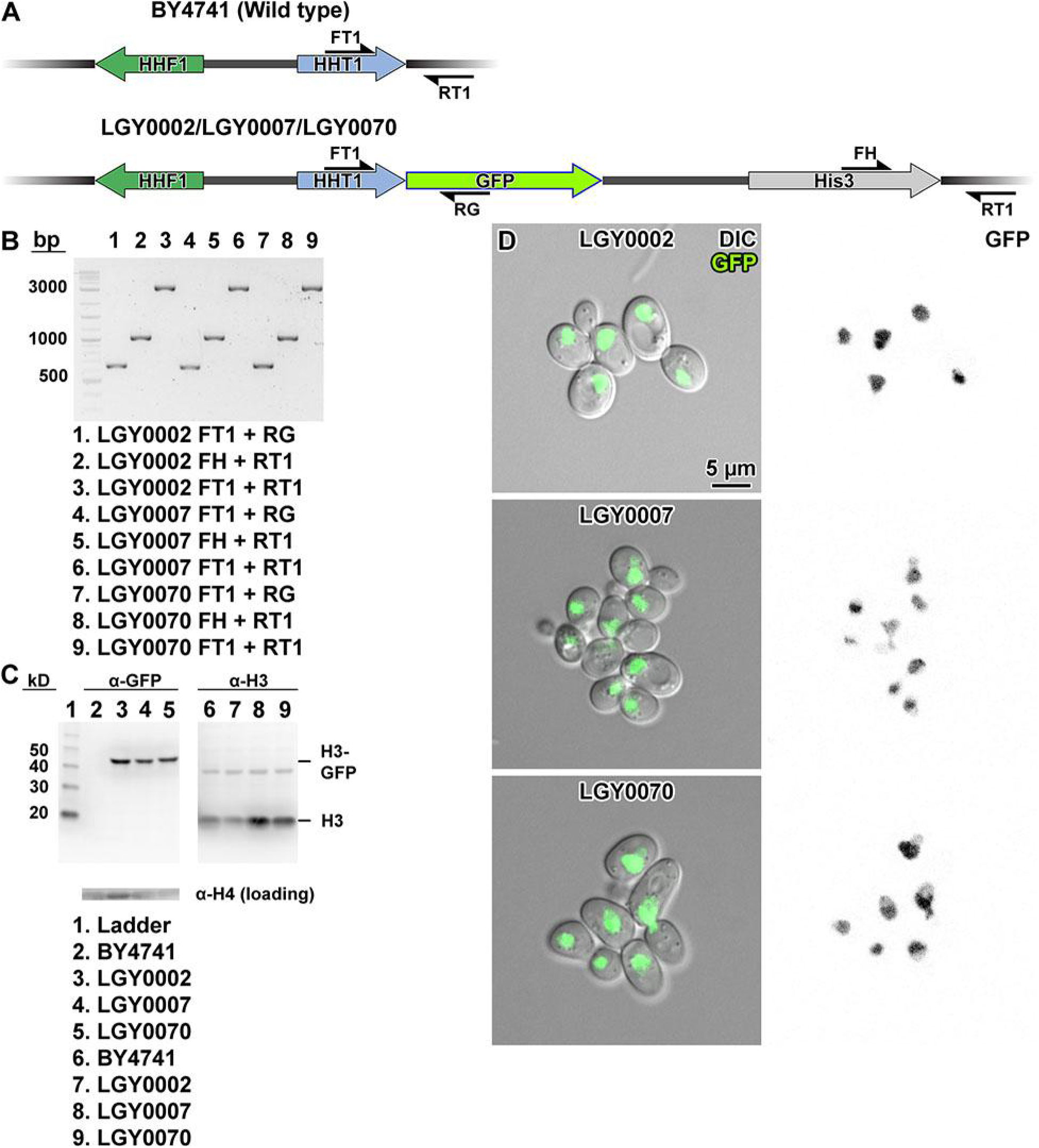
Verification of (H3, H3-GFP) strains. (A) Map of the HHT1-HHF1 locus in the parent (Wild type) strain BY4741 and the H3-GFP strains LGY0002 (H3, H3-RIPGLIN-GFP), LGY0007 (H3, H3-0aa-GFP) and LGY0070 (H3, H3-GGSGGS-GFP). Primers used for PCR verification are indicated with the half arrow symbols. (B) Agarose gel of PCR amplicons expected from LGY0002, LGY0007 and LGY0070 genomic DNA, in which the HHT1 locus is tagged with GFP. The only difference between the strains is the linker separating HHT1 and the GFP, which is too small to observe a difference in PCR amplicon sizes. (C) Immunoblot analysis of strains LGY0002, LGY0007 and LGY0070. The α-GFP antibody correctly detected the large H3-GFP fusion protein in LGY0002 (lane 3), LGY0007 (lane 4) and LGY0070 (lane 5), but not in BY4741 (lane 2, negative control). The α-H3 antibody detected H3 in all lanes, but failed to detect H3-GFP in LGY0002, LGY0007 and LGY0070. (D) DIC and GFP fluorescence confocal microscopy for LGY0002, LGY0007 and LGY0070 cells. In the left panel, the GFP signals are overlaid in green. In the right panel, GFP signals are rendered with inverted contrast.

**Figure S27.**
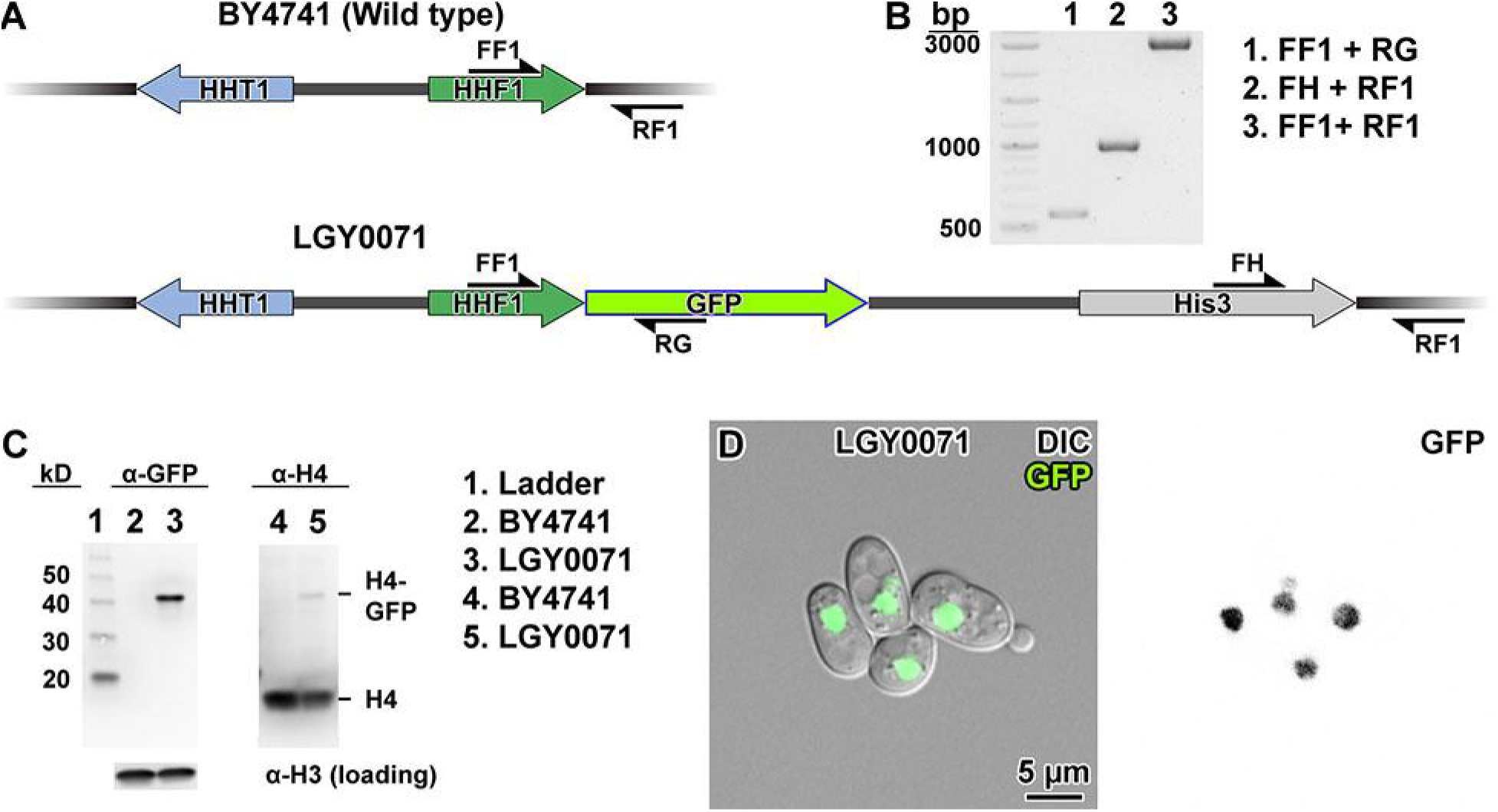
Verification of (H4, H4-GFP) strain. (A) Map of the HHT1-HHF1 locus in the parent (Wild type) strain BY4741 and the H4-GFP strain LGY0071. Primers used for PCR verification are indicated with the half arrow symbols. (B) Agarose gel of PCR amplicons expected from LGY0071 genomic DNA, in which the HHF1 locus is tagged with GFP. (C) Immunoblot analysis of strain LGY0071. The α-GFP antibody correctly detected the large H3-GFP fusion protein in LGY0071 (lane 3), but not in BY4741 (lane 2, negative control). The α-H4 antibody detected H4-GFP in LGY0071 and H4 in both lanes. (D) DIC and GFP fluorescence confocal microscopy for LGY0002, LGY0007 and LGY0070 cells. In the left panel, the GFP signals are overlaid in green. In the right panel, GFP signals are rendered with inverted contrast.

**Figure S28.**
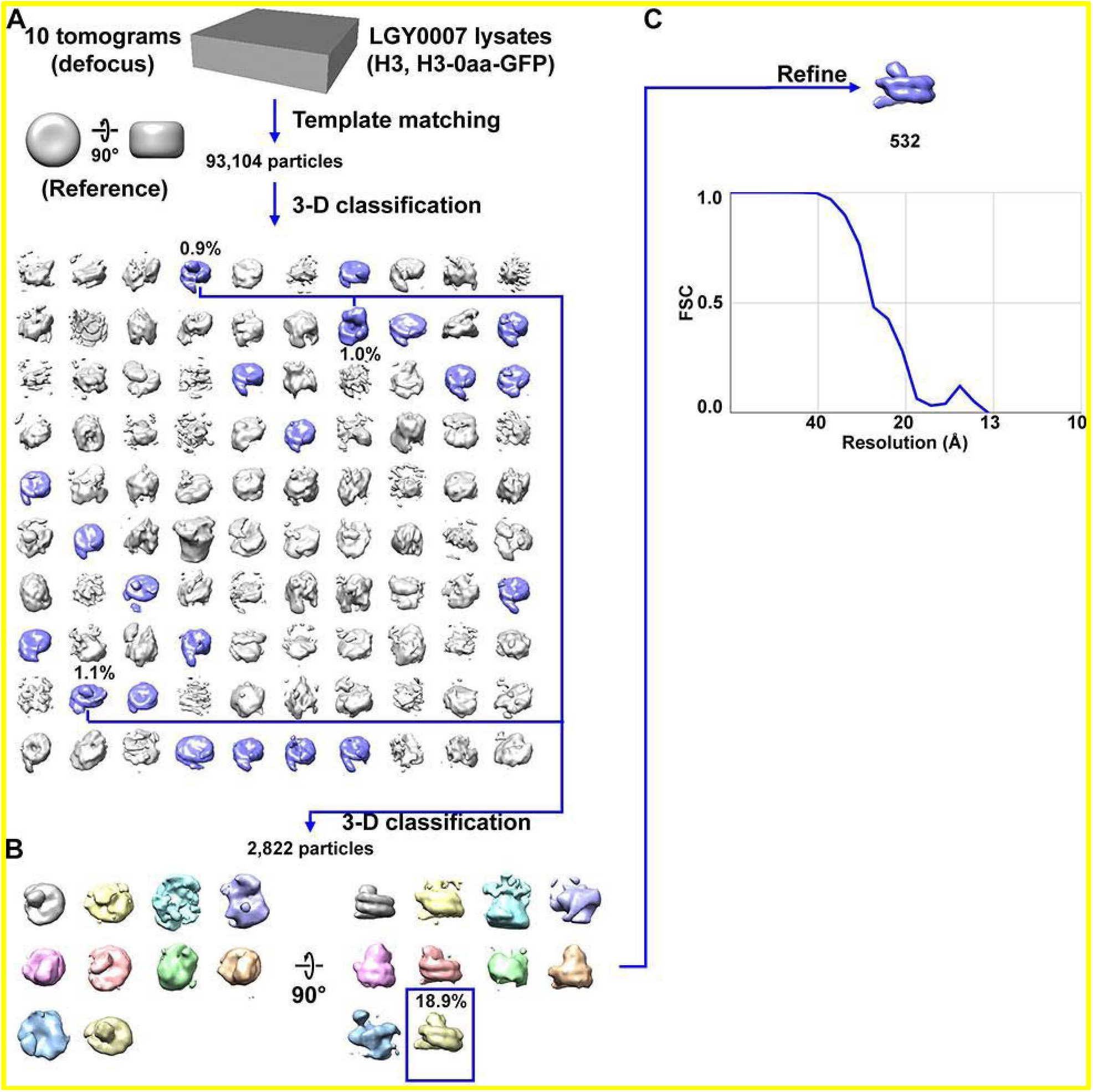
Direct 3-D classification of LGY0007 (H3, H3-0aa-GFP) nuclear lysates. (A) Class averages (3-D) of nucleosome-like particles from nuclear lysates of LGY0007 cells, which express a wild-type copy of H3 and an H3-GFP that has no linker sequence. The canonical nucleosome-like class averages with an additional GFP density are shaded blue while the non-canonical nucleosome averages and canonical nucleosome-like class averages without any additional density are shaded gray. (B) The second round of 3-D classification, using the canonical nucleosomes with an additional GFP density from panel A. The remove duplicates function of RELION was used to remove particles (defined as particles within a radius of 180 Å from another particle) from the indicated class average for the 3-D refinement. Twenty-seven duplicate particles were removed before refining. (C) Refined density of LGY0007 nucleosomes isolated from nuclear lysates. The number of particles in the chosen class is labeled in black. The resolution is ∼23 Å by the FSC = 0.5 criterion.

**Figure S29.**
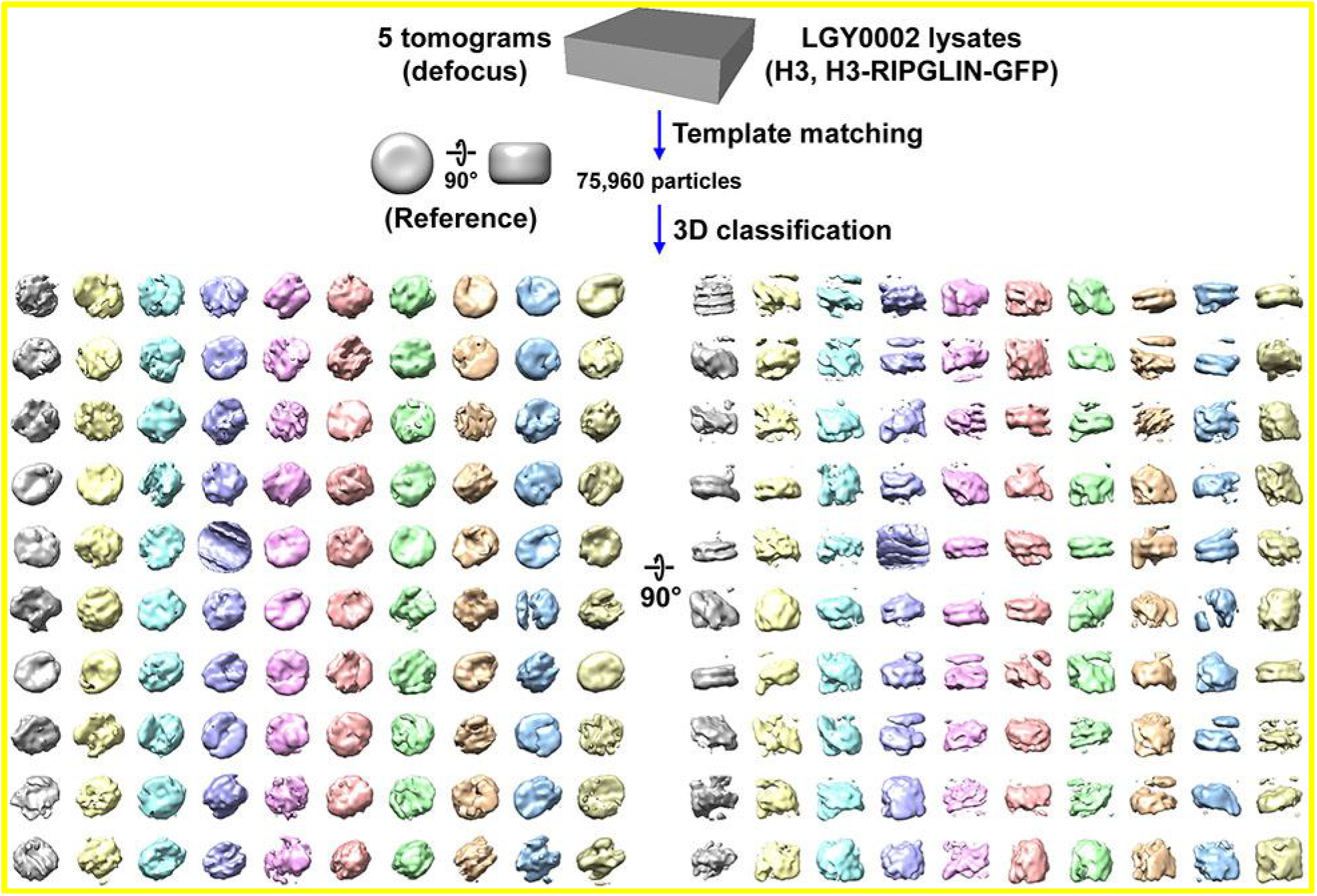
Direct 3-D classification of LGY0002 (H3, H3-RIPGLIN-GFP) nuclear lysates. Class averages (3-D) of nucleosome-like particles from nuclear lysates of LGY0002 cells, which express a wild-type copy of H3 and an H3-GFP that has a RIPGLIN linker sequence. In this experiment, 2-D classification was bypassed.

**Figure S30.**
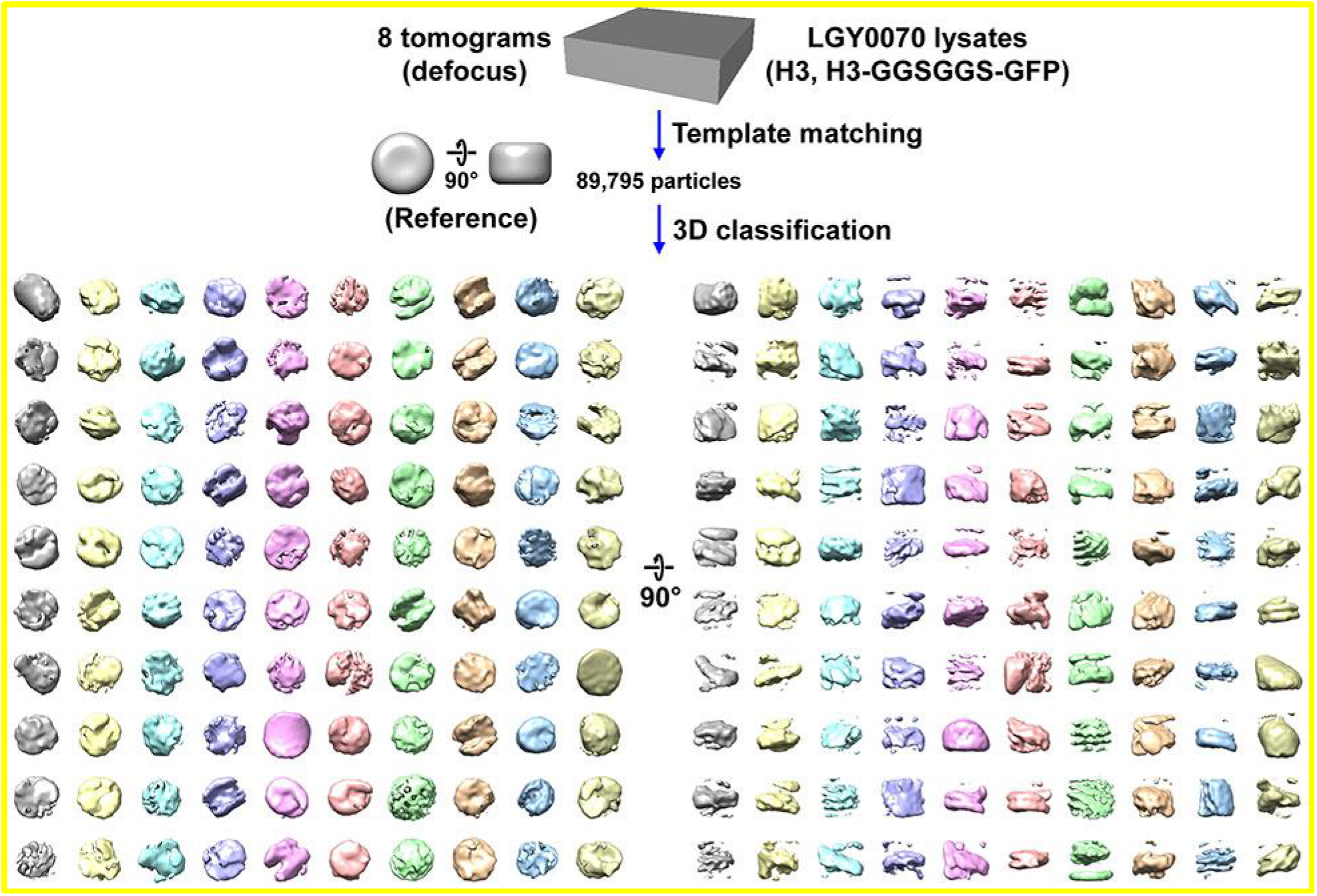
Direct 3-D classification of LGY0070 (H3, H3-GGSGGS-GFP) nuclear lysates. Class averages (3-D) of nucleosome-like particles from nuclear lysates of LGY0070 cells, which express a wild-type copy of H3 and an H3-GFP that has a GGSGGS linker sequence. In this experiment, 2-D classification was bypassed.

**Figure S31.**
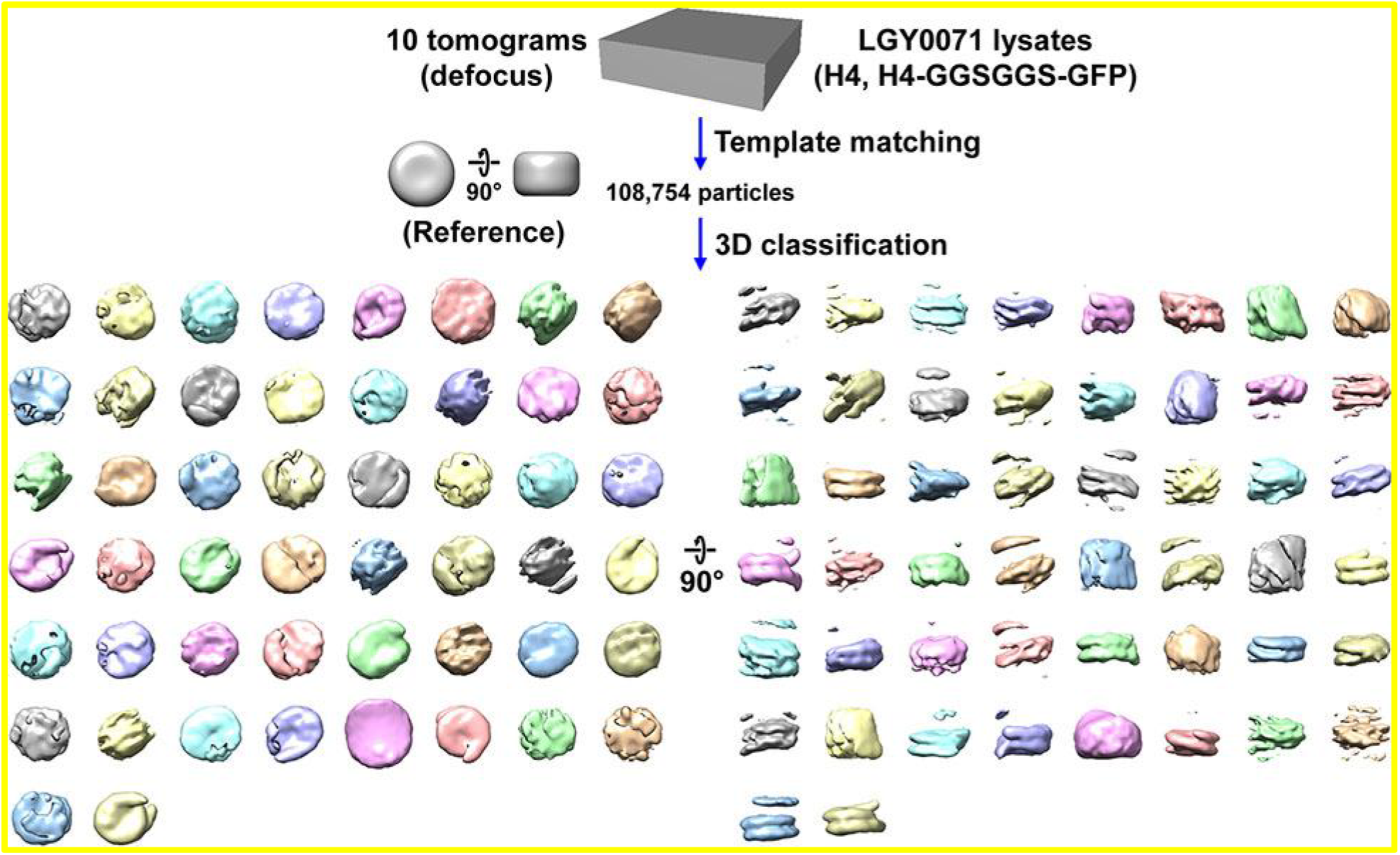
Direct 3-D classification of LGY0071 (H4, H4-GGSGGS-GFP) nuclear lysates. Class averages (3-D) of nucleosome-like particles from nuclear lysates of LGY0071 cells, which express a wild-type copy of H4 and an H4-GFP that has a GGSGGS linker sequence. In this experiment, 2-D classification was bypassed.

**Figure S32.**
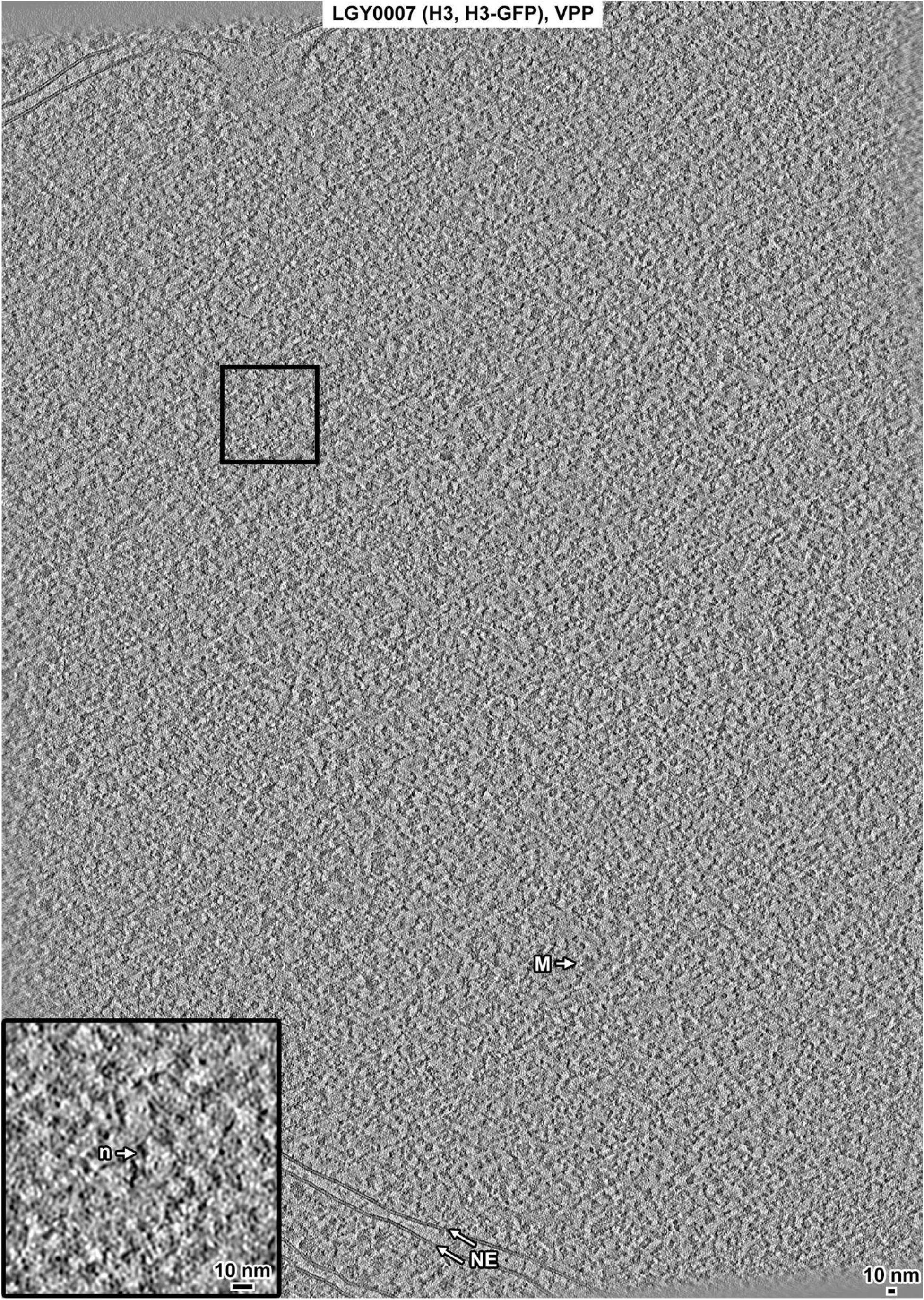
Overview of a LGY0007 (H3, H3-GFP) cell cryolamella, VPP data. Tomographic slice (12 nm) of a LGY0007 cell cryolamella imaged with a Volta phase plate (VPP). Some non-chromatin features are indicated: nuclear envelope (NE), megacomplex (M), and ribosome (R). The inset is a threefold enlargement of the boxed area. A nucleosome-like particle (n) is indicated.

**Figure S33.**
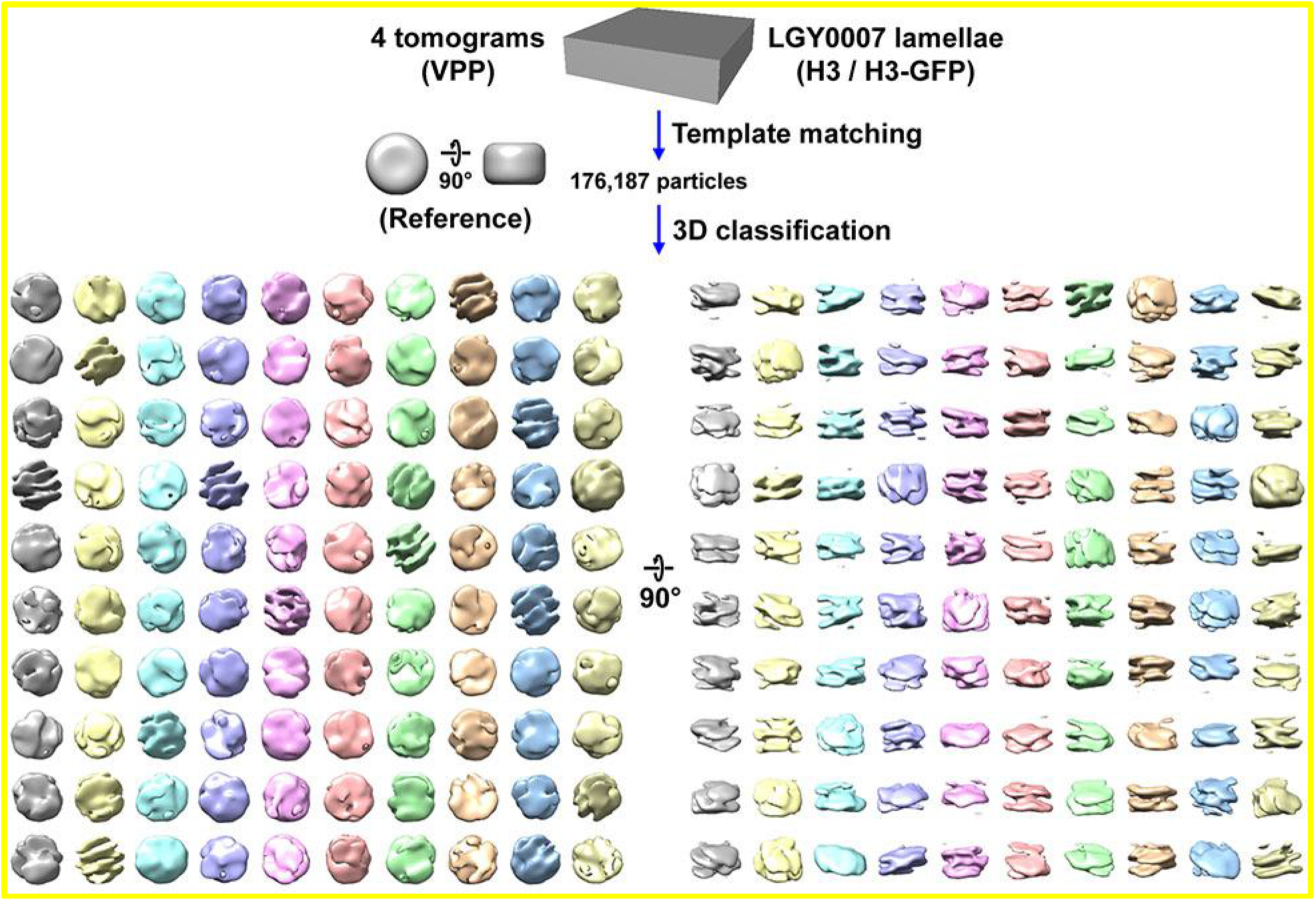
Direct 3-D classification of LGY0007 (H3, H3-GFP) cryolamellae densities. Class averages (3-D) of nucleosome-like particles in cryotomograms of LGY0007 cell cryolamellae. The cryolamellae were imaged with a Volta phase plate (VPP). In this experiment, 2-D classification was bypassed.

**Figure S34.**
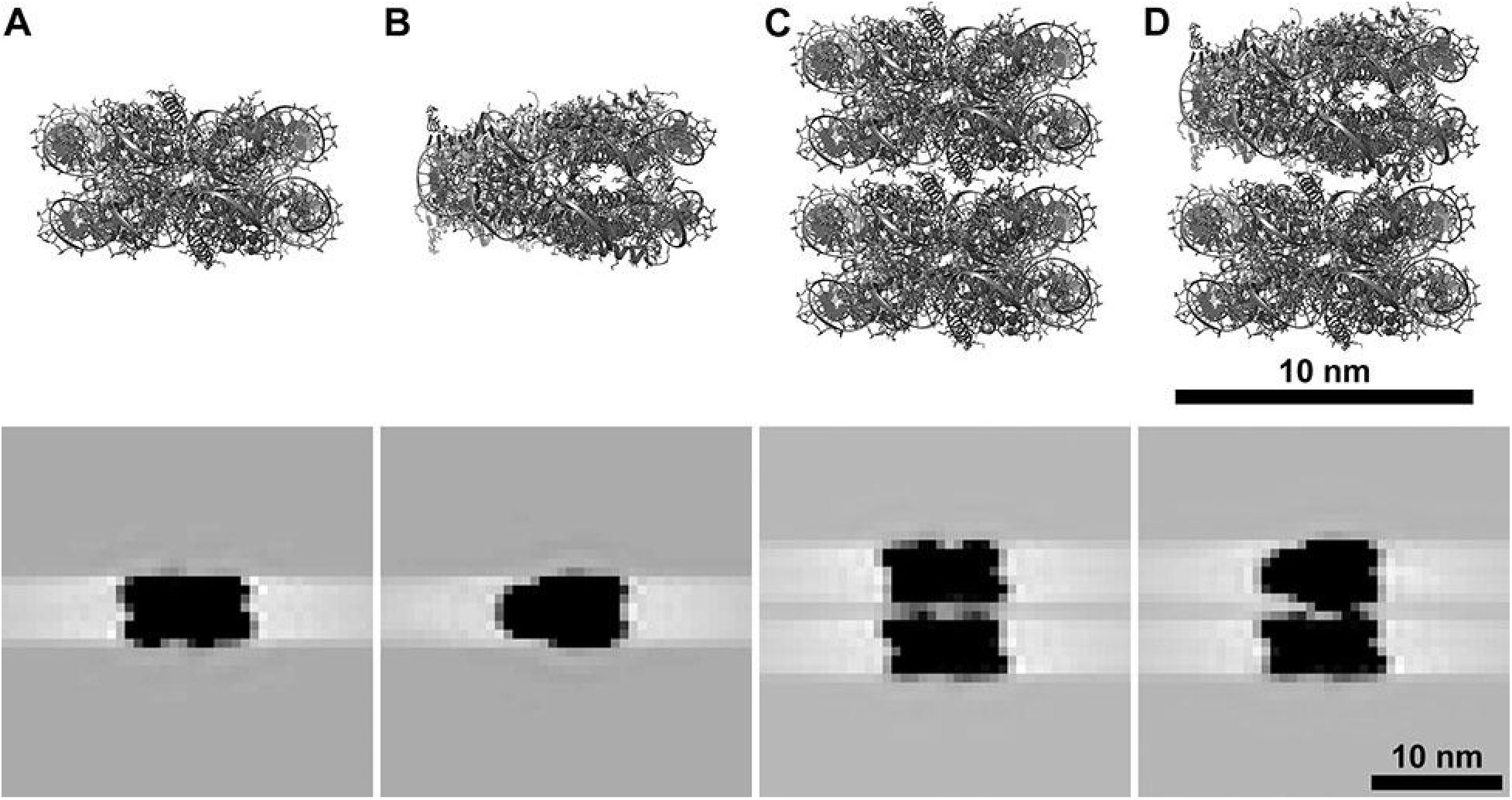
Simulated tomographic slices of single and stacked nucleosomes. Atomic models (upper, from PDB 1KX5) were used to simulate tomographic slices (12 nm, lower). (A and B) Mononucleosomes in the gyre and side views, respectively. (C and D) Two examples of stacked nucleosomes. In panel C, both nucleosomes have the same orientation. In panel D, the upper nucleosome is rotated 90° along the Y axis relative to the lower nucleosome. The stacked nucleosomes were separated by 55 Å center-to-center along the Y axis to emulate a worst-case (for image processing) scenario. Note that these simulated tomographic slices are not intended to accurately model the image-formation process. They serve to show how different side/gyre views of stacked nucleosomes appear.

**Figure S35.**
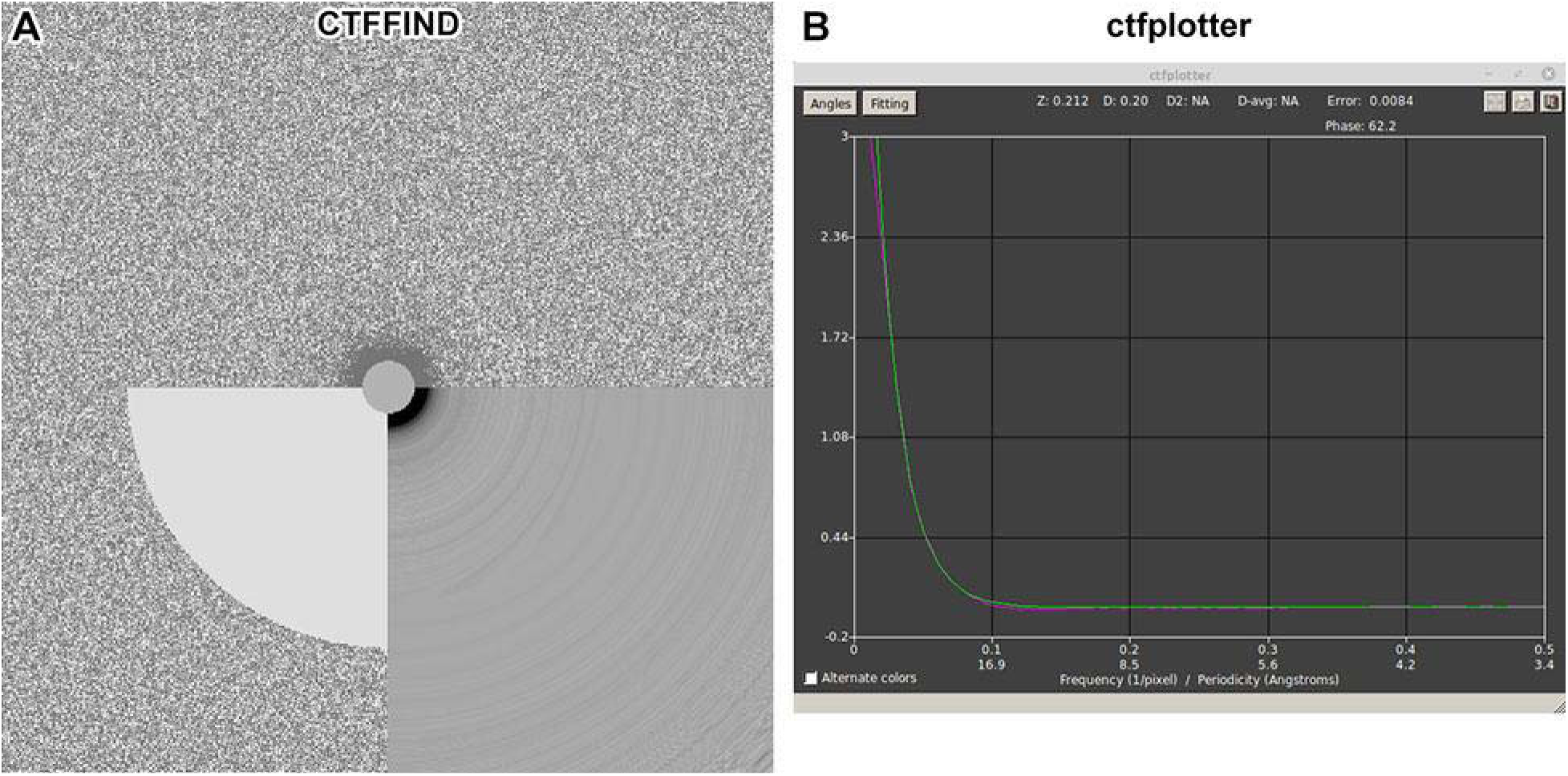
Fourier power spectra analysis of tilt series images. An image corresponding to a ∼12° pre-tilt (to make the cryolamella perpendicular to the electron-optical axis to maximize the signal-to-noise ratio) was extracted from tilt series 20211117_004 and then Fourier transformed using (A) CTFFIND and (B) IMOD ctfplotter. Thon rings could not be detected in the CTFFIND 2-D diagnostic image or the IMOD 1-D radial average of the Fourier power spectrum.

**Movie S1. Direct 3-D classification of BY4741 (Wild type) nucleosome-like particles in VPP tomograms of cell cryolamellae, round 1**

The progress of 30 rounds of 3-D classification is shown. There are 100 classes, initialized with a smooth nucleosome-sized cylindrical reference. The final iteration (30) is also shown in Figure. S10, with only the most nucleosome-like classes shaded.

**Movie S2. Direct 3-D classification of BY4741 (Wild type) nucleosome-like particles in VPP tomograms of cell cryolamellae, round 2**

The most nucleosome-like classes from round 1 were selected and subjected to a second round of classification, with 4 classes, again initialized with a smooth nucleosome-sized cylindrical reference.

**Movie S3. Direct 3-D classification of LGY0016 (H2A-GFP) lysate nucleosomes, round 1**

The progress of 30 rounds of 3-D classification is shown. There are 40 classes, initialized with a smooth nucleosome-sized cylindrical reference. The final iteration (30) is also shown in Figure. S18, with only the most nucleosome-like classes shaded.

**Movie S4. Direct 3-D classification of LGY0016 (H2A-GFP) lysate nucleosomes, round 2**

The most nucleosome-like classes from round 1 were selected and subjected to a second round of classification, with 10 classes, again initialized with a smooth nucleosome-sized cylindrical reference. Two “junk” classes were removed.

**Movie S5. Direct 3-D classification of LGY0016 (H2A-GFP) nucleosome-like particles in VPP tomograms of cell cryolamellae, round 1**

3-D classification analysis of LGY0016 (H2A-GFP) nucleosome-like particles from cryolamellae imaged with a VPP. The initial reference is a featureless cylinder. There are 100 classes.

**Table S1.**
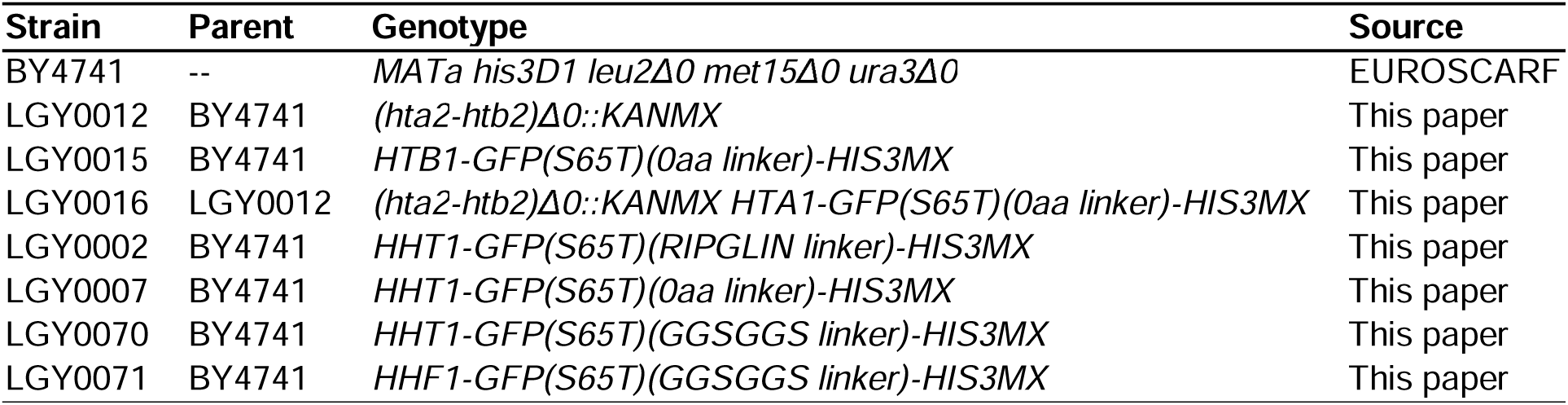
Genotypes of strains used in this paperxs.

**Table S2.**
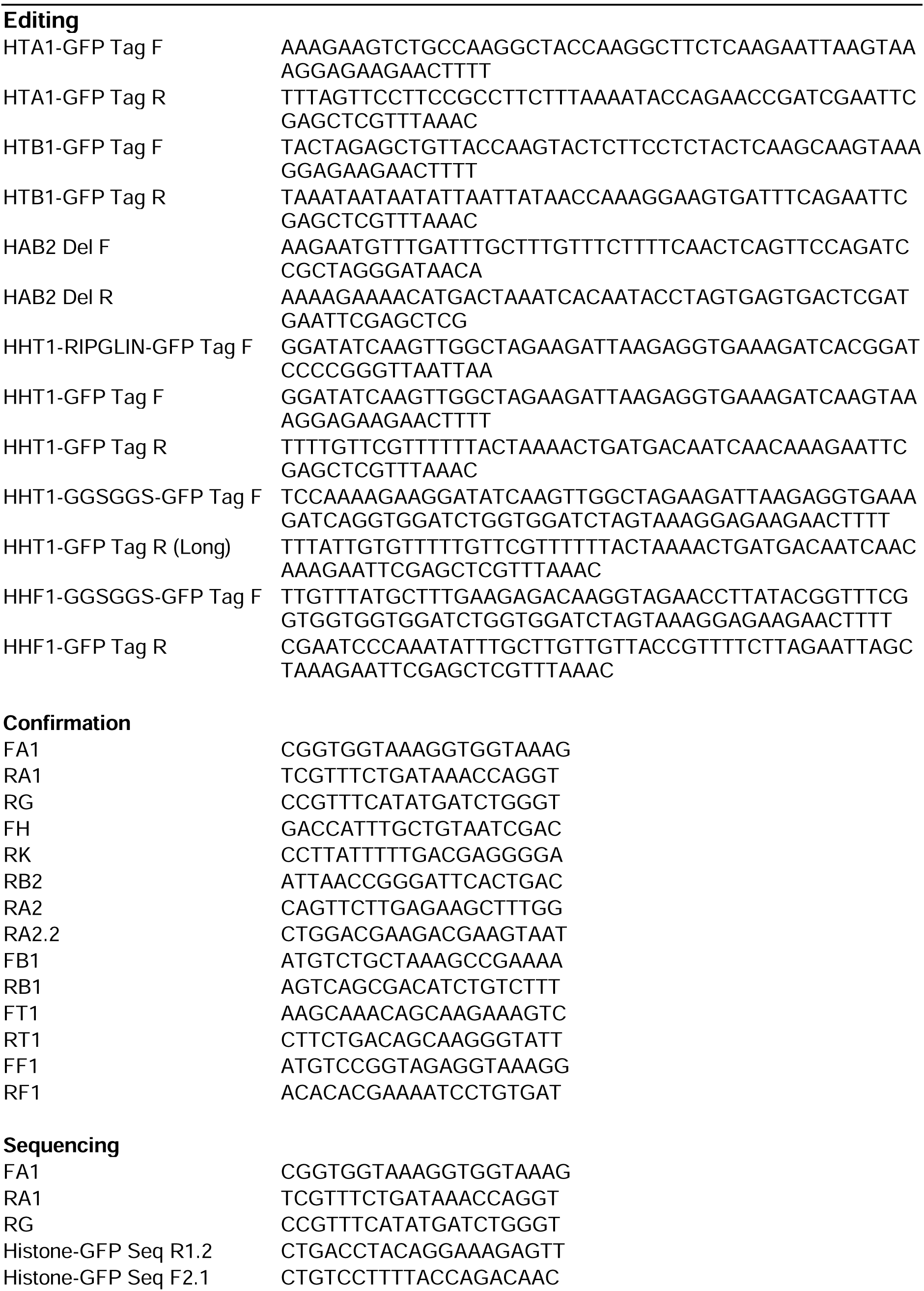
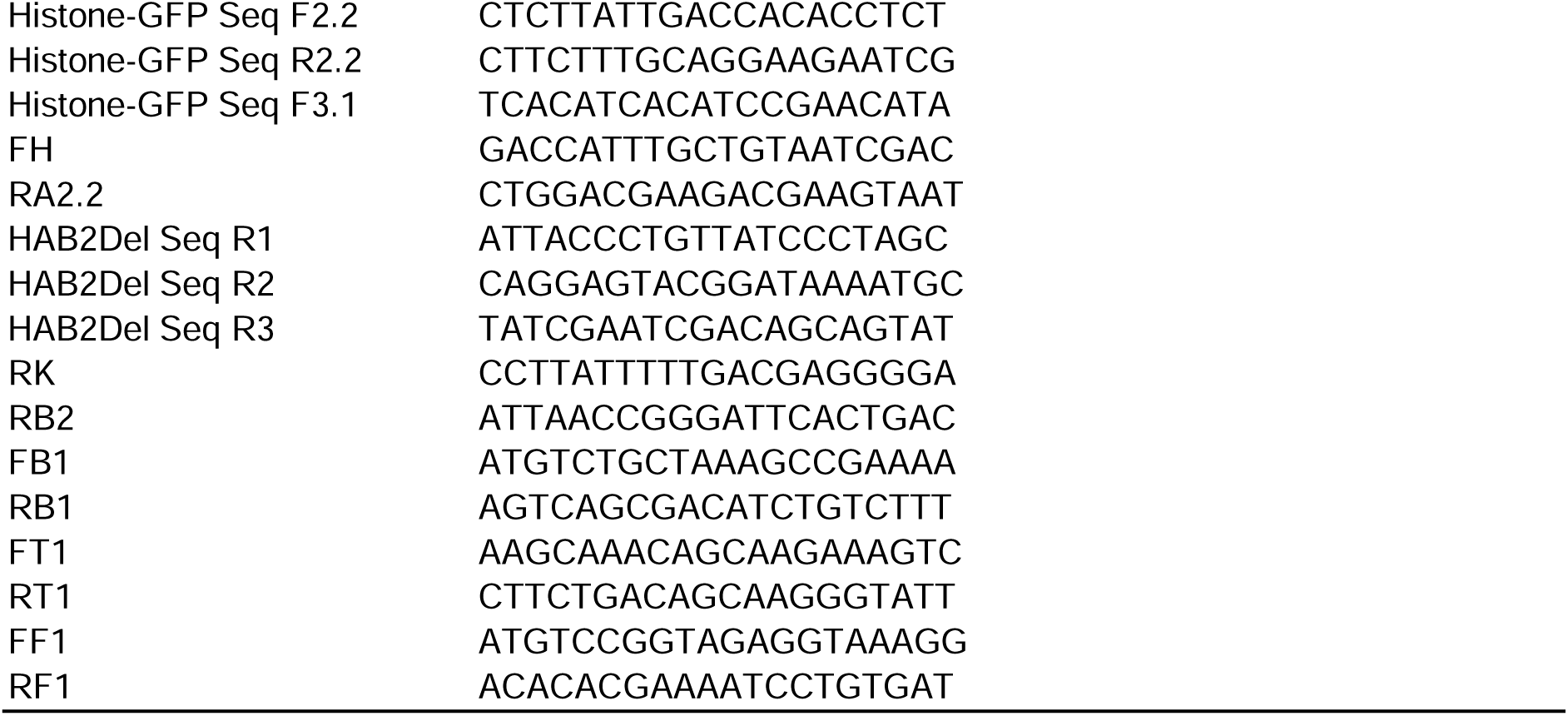
PCR primers, 5’ → 3’.

**Table S3.** Antibodies for immunoblots.

| Antigen | 1° antibody | 2° antibody | Dilution |  |
| --- | --- | --- | --- | --- |
|  |  |  | 1° | 2° |
| H2A | Active Motif 39235 | CST 7074S | 1:1000 | 1:5000 |
| H2B | Abcam ab1790 | CST 7074S | 1:1000 | 1:5000 |
| H3 | Abcam ab1791 | CST 7074S | 1:1000 | 1:5000 |
| H4 | Abcam ab10158 | CST 7074S | 1:1000 | 1:5000 |
| GFP | Santa Cruz sc9996 | CST 7076S | 1:1000 | 1:5000 |

CST = Cell Signaling Technology

**Table S4.**
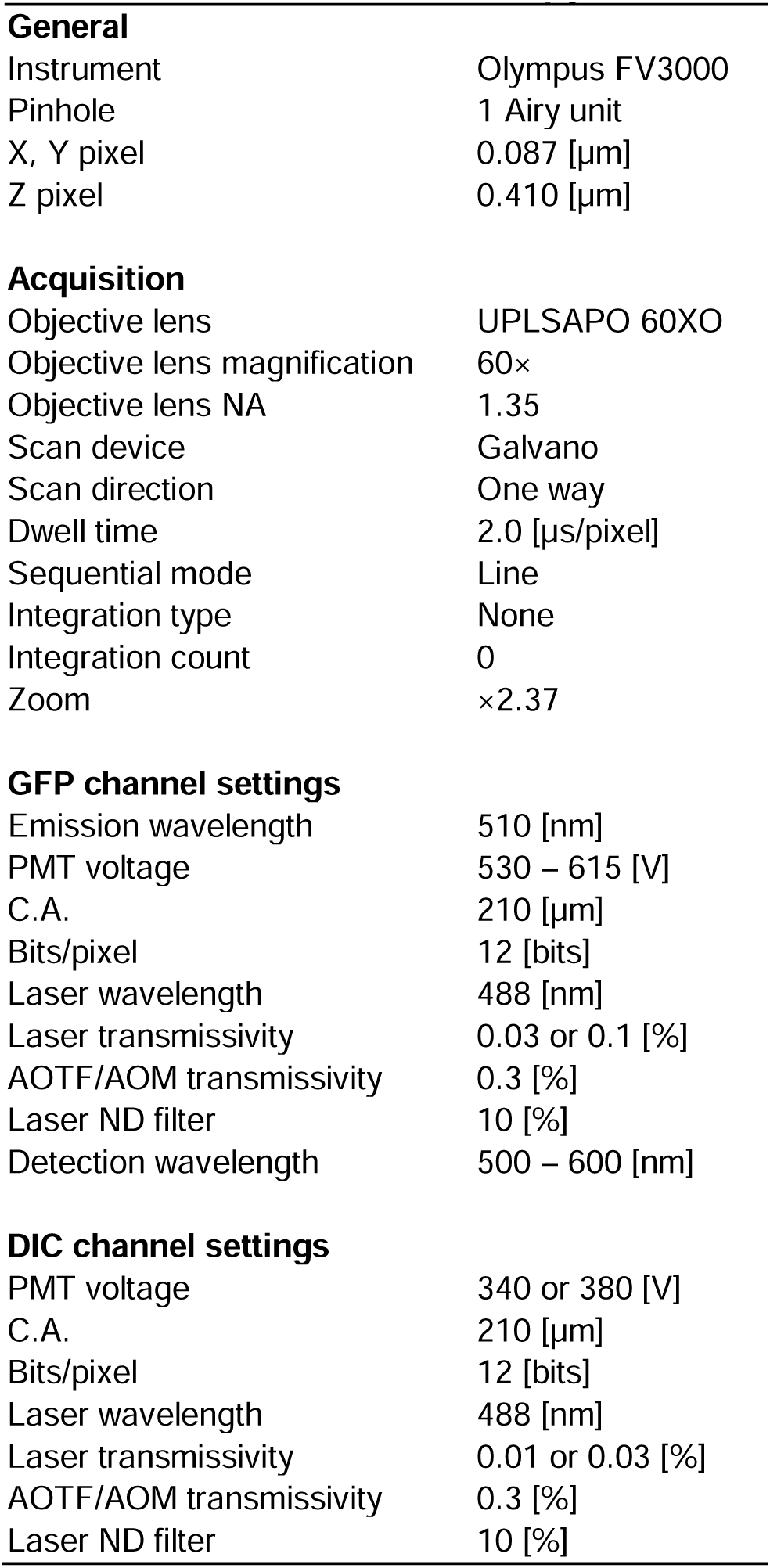
Confocal microscopy details.

**Table S5.**
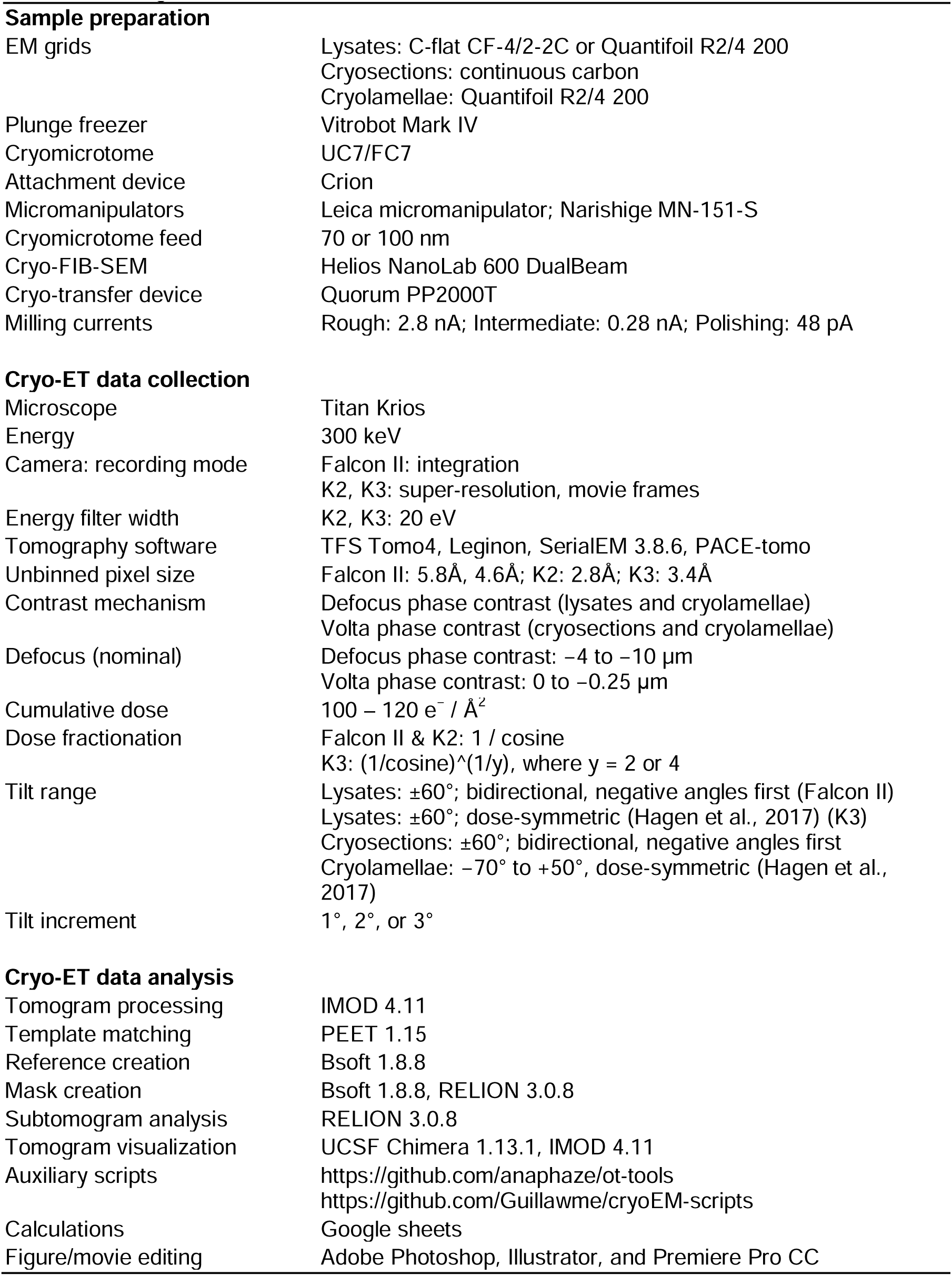
**Cryo-ET details**.

**Table S6.**
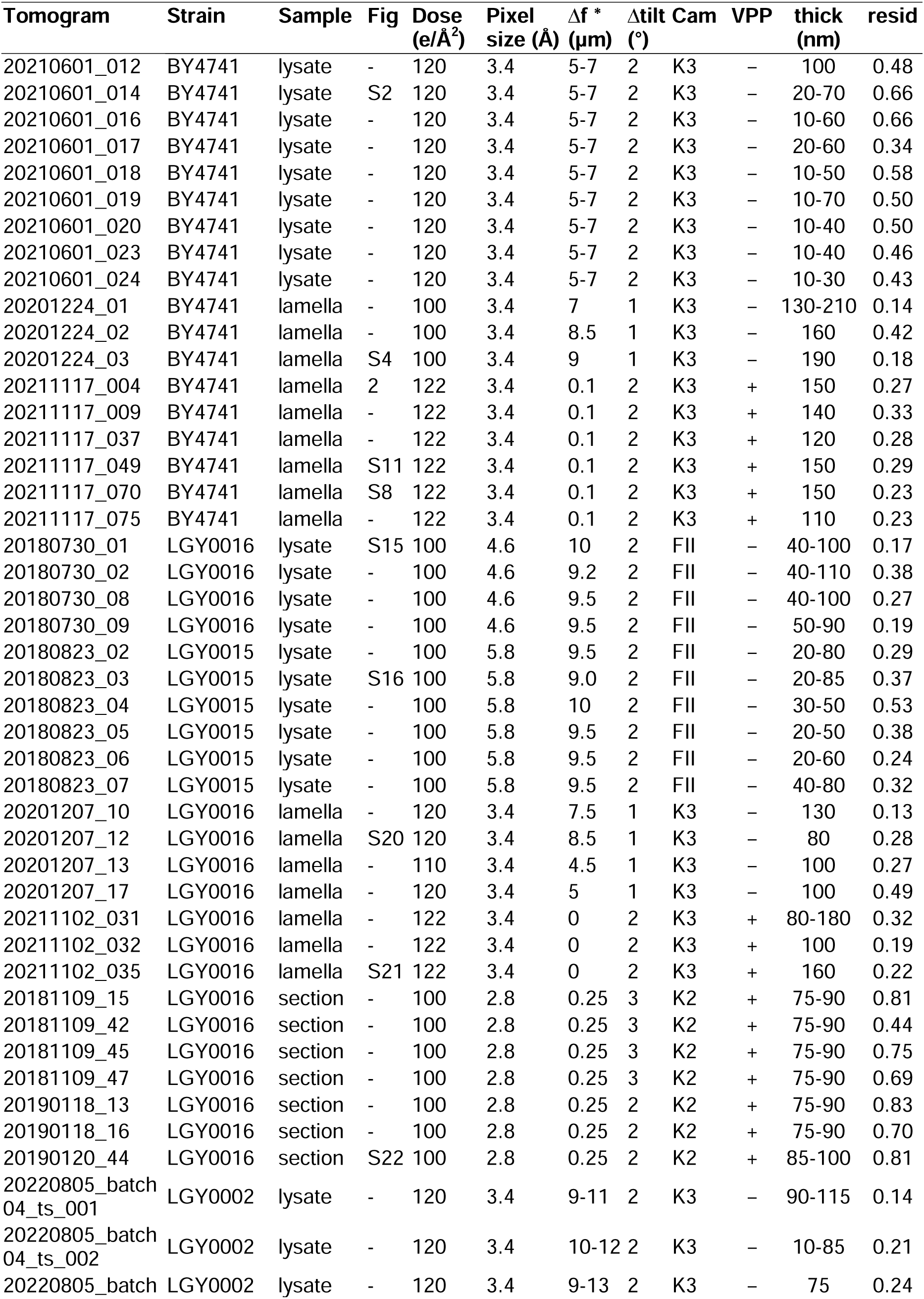
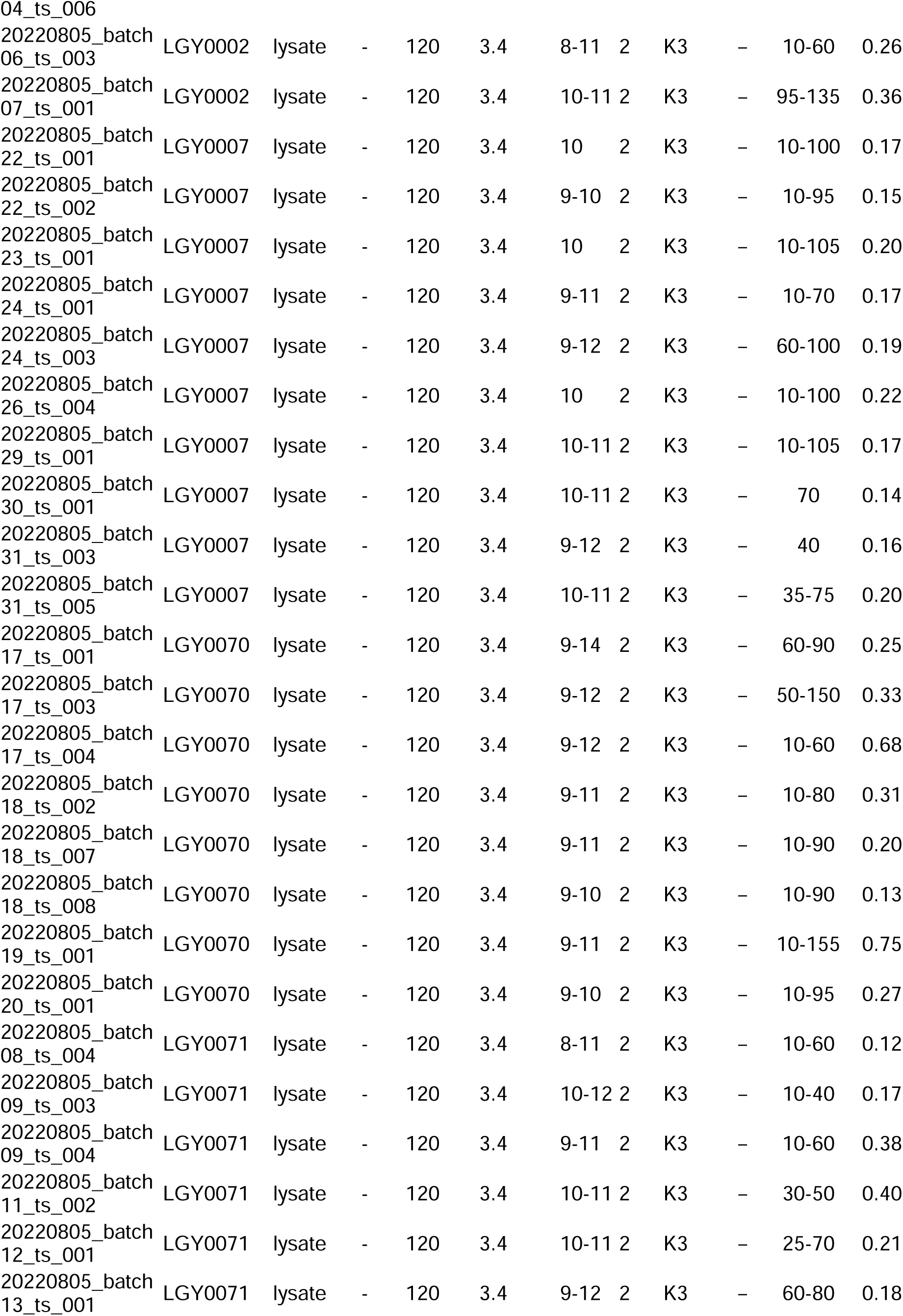
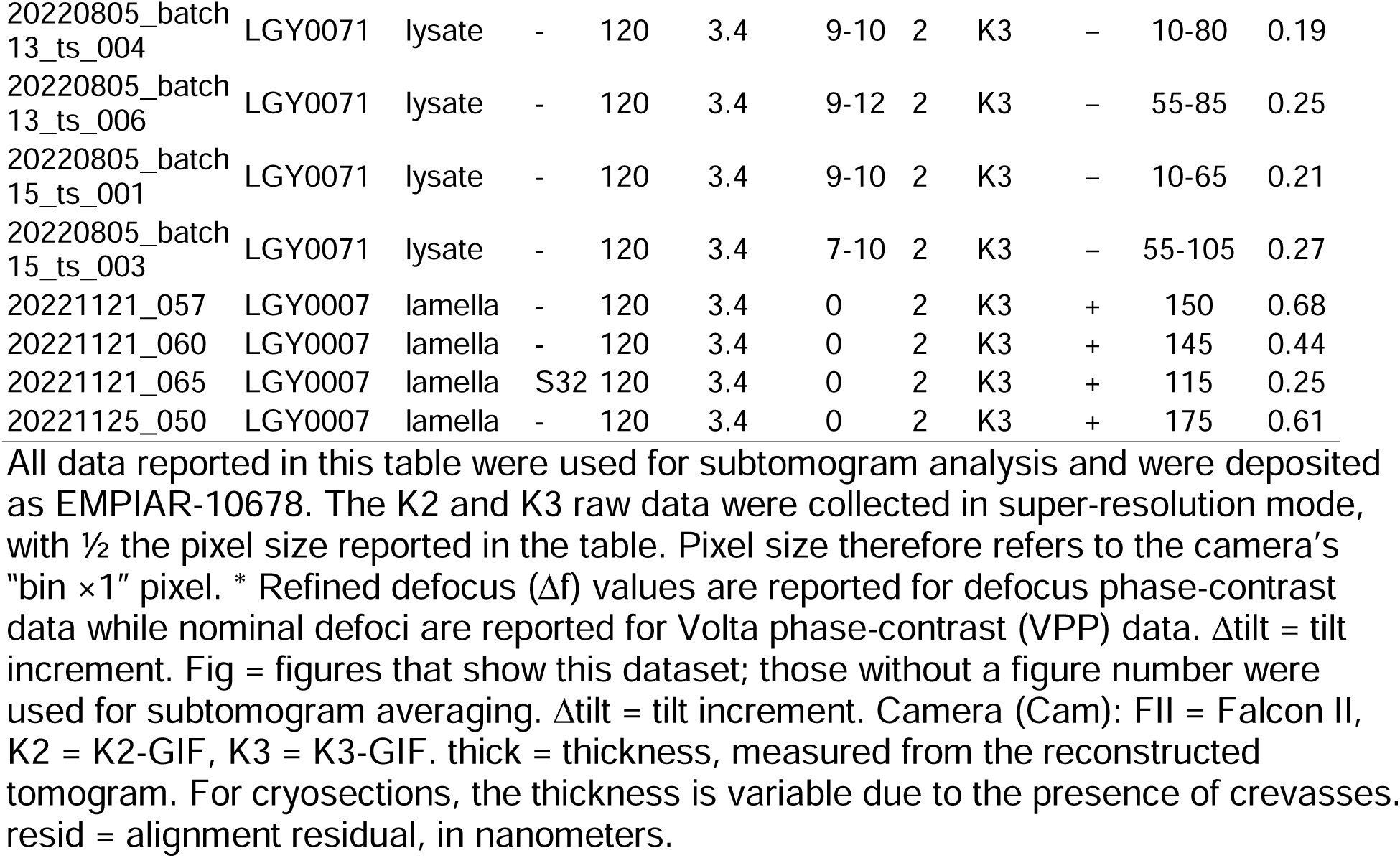
Cryotomogram details.

| Tomogram | Strain | Sample | Fig | Dose<br>(e/Å<sup>2</sup>) | Pixel<br>size (Å) | Δf *<br>(μm) | Δtilt<br>(°) | Cam | VPP | thick<br>(nm) | resid |
| --- | --- | --- | --- | --- | --- | --- | --- | --- | --- | --- | --- |
| 20210601_012 | BY4741 | lysate | - | 120 | 3.4 | 5-7 | 2 | K3 | - | 100 | 0.48 |
| 20210601_014 | BY4741 | lysate | S2 | 120 | 3.4 | 5-7 | 2 | K3 | - | 20-70 | 0.66 |
| 20210601_016 | BY4741 | lysate | - | 120 | 3.4 | 5-7 | 2 | K3 | - | 10-60 | 0.66 |
| 20210601_017 | BY4741 | lysate | - | 120 | 3.4 | 5-7 | 2 | K3 | - | 20-60 | 0.34 |
| 20210601_018 | BY4741 | lysate | - | 120 | 3.4 | 5-7 | 2 | K3 | - | 10-50 | 0.58 |
| 20210601_019 | BY4741 | lysate | - | 120 | 3.4 | 5-7 | 2 | K3 | - | 10-70 | 0.50 |
| 20210601_020 | BY4741 | lysate | - | 120 | 3.4 | 5-7 | 2 | K3 | - | 10-40 | 0.50 |
| 20210601_023 | BY4741 | lysate | - | 120 | 3.4 | 5-7 | 2 | K3 | - | 10-40 | 0.46 |
| 20210601_024 | BY4741 | lysate | - | 120 | 3.4 | 5-7 | 2 | K3 | - | 10-30 | 0.43 |
| 20201224_01 | BY4741 | lamella | - | 100 | 3.4 | 7 | 1 | K3 | - | 130-210 | 0.14 |
| 20201224_02 | BY4741 | lamella | - | 100 | 3.4 | 8.5 | 1 | K3 | - | 160 | 0.42 |
| 20201224_03 | BY4741 | lamella | S4 | 100 | 3.4 | 9 | 1 | K3 | - | 190 | 0.18 |
| 20211117_004 | BY4741 | lamella | 2 | 122 | 3.4 | 0.1 | 2 | K3 | + | 150 | 0.27 |
| 20211117_009 | BY4741 | lamella | - | 122 | 3.4 | 0.1 | 2 | K3 | + | 140 | 0.33 |
| 20211117_037 | BY4741 | lamella | - | 122 | 3.4 | 0.1 | 2 | K3 | + | 120 | 0.28 |
| 20211117_049 | BY4741 | lamella | S11 | 122 | 3.4 | 0.1 | 2 | K3 | + | 150 | 0.29 |
| 20211117_070 | BY4741 | lamella | S8 | 122 | 3.4 | 0.1 | 2 | K3 | + | 150 | 0.23 |
| 20211117_075 | BY4741 | lamella | - | 122 | 3.4 | 0.1 | 2 | K3 | + | 110 | 0.23 |
| 20180730_01 | LGY0016 | lysate | S15 | 100 | 4.6 | 10 | 2 | FII | - | 40-100 | 0.17 |
| 20180730_02 | LGY0016 | lysate | - | 100 | 4.6 | 9.2 | 2 | FII | - | 40-110 | 0.38 |
| 20180730_08 | LGY0016 | lysate | - | 100 | 4.6 | 9.5 | 2 | FII | - | 40-100 | 0.27 |
| 20180730_09 | LGY0016 | lysate | - | 100 | 4.6 | 9.5 | 2 | FII | - | 50-90 | 0.19 |
| 20180823_02 | LGY0015 | lysate | - | 100 | 5.8 | 9.5 | 2 | FII | - | 20-80 | 0.29 |
| 20180823_03 | LGY0015 | lysate | S16 | 100 | 5.8 | 9.0 | 2 | FII | - | 20-85 | 0.37 |
| 20180823_04 | LGY0015 | lysate | - | 100 | 5.8 | 10 | 2 | FII | - | 30-50 | 0.53 |
| 20180823_05 | LGY0015 | lysate | - | 100 | 5.8 | 9.5 | 2 | FII | - | 20-50 | 0.38 |
| 20180823_06 | LGY0015 | lysate | - | 100 | 5.8 | 9.5 | 2 | FII | - | 20-60 | 0.24 |
| 20180823_07 | LGY0015 | lysate | - | 100 | 5.8 | 9.5 | 2 | FII | - | 40-80 | 0.32 |
| 20201207_10 | LGY0016 | lamella | - | 120 | 3.4 | 7.5 | 1 | K3 | - | 130 | 0.13 |
| 20201207_12 | LGY0016 | lamella | S20 | 120 | 3.4 | 8.5 | 1 | K3 | - | 80 | 0.28 |
| 20201207_13 | LGY0016 | lamella | - | 110 | 3.4 | 4.5 | 1 | K3 | - | 100 | 0.27 |
| 20201207_17 | LGY0016 | lamella | - | 120 | 3.4 | 5 | 1 | K3 | - | 100 | 0.49 |
| 20211102_031 | LGY0016 | lamella | - | 122 | 3.4 | 0 | 2 | K3 | + | 80-180 | 0.32 |
| 20211102_032 | LGY0016 | lamella | - | 122 | 3.4 | 0 | 2 | K3 | + | 100 | 0.19 |
| 20211102_035 | LGY0016 | lamella | S21 | 122 | 3.4 | 0 | 2 | K3 | + | 160 | 0.22 |
| 20181109_15 | LGY0016 | section | - | 100 | 2.8 | 0.25 | 3 | K2 | + | 75-90 | 0.81 |
| 20181109_42 | LGY0016 | section | - | 100 | 2.8 | 0.25 | 3 | K2 | + | 75-90 | 0.44 |
| 20181109_45 | LGY0016 | section | - | 100 | 2.8 | 0.25 | 3 | K2 | + | 75-90 | 0.75 |
| 20181109_47 | LGY0016 | section | - | 100 | 2.8 | 0.25 | 3 | K2 | + | 75-90 | 0.69 |
| 20190118_13 | LGY0016 | section | - | 100 | 2.8 | 0.25 | 2 | K2 | + | 75-90 | 0.83 |
| 20190118_16 | LGY0016 | section | - | 100 | 2.8 | 0.25 | 2 | K2 | + | 75-90 | 0.70 |
| 20190120_44 | LGY0016 | section | S22 | 100 | 2.8 | 0.25 | 2 | K2 | + | 85-100 | 0.81 |
| 20220805_batch<br>04_ts_001 | LGY0002 | lysate | - | 120 | 3.4 | 9-11 | 2 | K3 | - | 90-115 | 0.14 |
| 20220805_batch<br>04_ts_002 | LGY0002 | lysate | - | 120 | 3.4 | 10-12 | 2 | K3 | - | 10-85 | 0.21 |
| 20220805_batch | LGY0002 | lysate | - | 120 | 3.4 | 9-13 | 2 | K3 | - | 75 | 0.24 |
| 04_ts_006 |  |  |  |  |  |  |  |  |  |  |  |
| 20220805_batch<br>06_ts_003 | LGY0002 | lysate | - | 120 | 3.4 | 8-11 | 2 | K3 | - | 10-60 | 0.26 |
| 20220805_batch<br>07_ts_001 | LGY0002 | lysate | - | 120 | 3.4 | 10-11 | 2 | K3 | - | 95-135 | 0.36 |
| 20220805_batch<br>22_ts_001 | LGY0007 | lysate | - | 120 | 3.4 | 10 | 2 | K3 | - | 10-100 | 0.17 |
| 20220805_batch<br>22_ts_002 | LGY0007 | lysate | - | 120 | 3.4 | 9-10 | 2 | K3 | - | 10-95 | 0.15 |
| 20220805_batch<br>23_ts_001 | LGY0007 | lysate | - | 120 | 3.4 | 10 | 2 | K3 | - | 10-105 | 0.20 |
| 20220805_batch<br>24_ts_001 | LGY0007 | lysate | - | 120 | 3.4 | 9-11 | 2 | K3 | - | 10-70 | 0.17 |
| 20220805_batch<br>24_ts_003 | LGY0007 | lysate | - | 120 | 3.4 | 9-12 | 2 | K3 | - | 60-100 | 0.19 |
| 20220805_batch<br>26_ts_004 | LGY0007 | lysate | - | 120 | 3.4 | 10 | 2 | K3 | - | 10-100 | 0.22 |
| 20220805_batch<br>29_ts_001 | LGY0007 | lysate | - | 120 | 3.4 | 10-11 | 2 | K3 | - | 10-105 | 0.17 |
| 20220805_batch<br>30_ts_001 | LGY0007 | lysate | - | 120 | 3.4 | 10-11 | 2 | K3 | - | 70 | 0.14 |
| 20220805_batch<br>31_ts_003 | LGY0007 | lysate | - | 120 | 3.4 | 9-12 | 2 | K3 | - | 40 | 0.16 |
| 20220805_batch<br>31_ts_005 | LGY0007 | lysate | - | 120 | 3.4 | 10-11 | 2 | K3 | - | 35-75 | 0.20 |
| 20220805_batch<br>17_ts_001 | LGY0070 | lysate | - | 120 | 3.4 | 9-14 | 2 | K3 | - | 60-90 | 0.25 |
| 20220805_batch<br>17_ts_003 | LGY0070 | lysate | - | 120 | 3.4 | 9-12 | 2 | K3 | - | 50-150 | 0.33 |
| 20220805_batch<br>17_ts_004 | LGY0070 | lysate | - | 120 | 3.4 | 9-12 | 2 | K3 | - | 10-60 | 0.68 |
| 20220805_batch<br>18_ts_002 | LGY0070 | lysate | - | 120 | 3.4 | 9-11 | 2 | K3 | - | 10-80 | 0.31 |
| 20220805_batch<br>18_ts_007 | LGY0070 | lysate | - | 120 | 3.4 | 9-11 | 2 | K3 | - | 10-90 | 0.20 |
| 20220805_batch<br>18_ts_008 | LGY0070 | lysate | - | 120 | 3.4 | 9-10 | 2 | K3 | - | 10-90 | 0.13 |
| 20220805_batch<br>19_ts_001 | LGY0070 | lysate | - | 120 | 3.4 | 9-11 | 2 | K3 | - | 10-155 | 0.75 |
| 20220805_batch<br>20_ts_001 | LGY0070 | lysate | - | 120 | 3.4 | 9-10 | 2 | K3 | - | 10-95 | 0.27 |
| 20220805_batch<br>08_ts_004 | LGY0071 | lysate | - | 120 | 3.4 | 8-11 | 2 | K3 | - | 10-60 | 0.12 |
| 20220805_batch<br>09_ts_003 | LGY0071 | lysate | - | 120 | 3.4 | 10-12 | 2 | K3 | - | 10-40 | 0.17 |
| 20220805_batch<br>09_ts_004 | LGY0071 | lysate | - | 120 | 3.4 | 9-11 | 2 | K3 | - | 10-60 | 0.38 |
| 20220805_batch<br>11_ts_002 | LGY0071 | lysate | - | 120 | 3.4 | 10-11 | 2 | K3 | - | 30-50 | 0.40 |
| 20220805_batch<br>12_ts_001 | LGY0071 | lysate | - | 120 | 3.4 | 10-11 | 2 | K3 | - | 25-70 | 0.21 |
| 20220805_batch<br>13_ts_001 | LGY0071 | lysate | - | 120 | 3.4 | 9-12 | 2 | K3 | - | 60-80 | 0.18 |
| 20220805_batch<br>13_ts_004 | LGY0071 | lysate | - | 120 | 3.4 | 9-10 | 2 | K3 | - | 10-80 | 0.19 |
| 20220805_batch<br>13_ts_006 | LGY0071 | lysate | - | 120 | 3.4 | 9-12 | 2 | K3 | - | 55-85 | 0.25 |
| 20220805_batch<br>15_ts_001 | LGY0071 | lysate | - | 120 | 3.4 | 9-10 | 2 | K3 | - | 10-65 | 0.21 |
| 20220805_batch<br>15_ts_003 | LGY0071 | lysate | - | 120 | 3.4 | 7-10 | 2 | K3 | - | 55-105 | 0.27 |
| 20221121_057 | LGY0007 | lamella | - | 120 | 3.4 | 0 | 2 | K3 | + | 150 | 0.68 |
| 20221121_060 | LGY0007 | lamella | - | 120 | 3.4 | 0 | 2 | K3 | + | 145 | 0.44 |
| 20221121_065 | LGY0007 | lamella | S32 | 120 | 3.4 | 0 | 2 | K3 | + | 115 | 0.25 |
| 20221125_050 | LGY0007 | lamella | - | 120 | 3.4 | 0 | 2 | K3 | + | 175 | 0.61 |

All data reported in this table were used for subtomogram analysis and were deposited as EMPIAR-10678. The K2 and K3 raw data were collected in super-resolution mode, with ½ the pixel size reported in the table. Pixel size therefore refers to the camera's "bin x1" pixel. \* Refined defocus ( $\Delta f$ ) values are reported for defocus phase-contrast data while nominal defoci are reported for Volta phase-contrast (VPP) data. $\Delta$ tilt = tilt increment. Fig = figures that show this dataset; those without a figure number were used for subtomogram averaging. $\Delta$ tilt = tilt increment. Camera (Cam): FII = Falcon II, K2 = K2-GIF, K3 = K3-GIF. thick = thickness, measured from the reconstructed tomogram. For cryosections, the thickness is variable due to the presence of crevasses. resid = alignment residual, in nanometers.

**Table S7.**
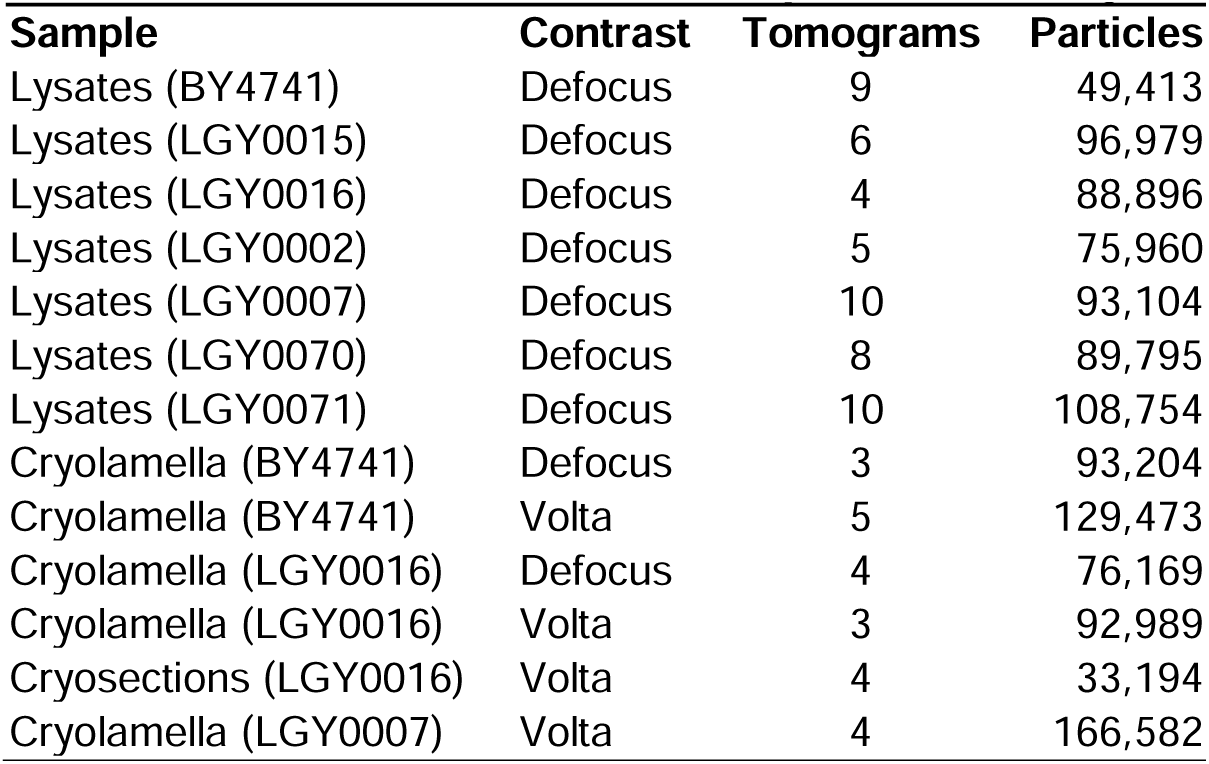
Total nucleosome-like particles analyzed.

| Sample | Contrast | Tomograms | Particles |
| --- | --- | --- | --- |
| Lysates (BY4741) | Defocus | 9 | 49,413 |
| Lysates (LGY0015) | Defocus | 6 | 96,979 |
| Lysates (LGY0016) | Defocus | 4 | 88,896 |
| Lysates (LGY0002) | Defocus | 5 | 75,960 |
| Lysates (LGY0007) | Defocus | 10 | 93,104 |
| Lysates (LGY0070) | Defocus | 8 | 89,795 |
| Lysates (LGY0071) | Defocus | 10 | 108,754 |
| Cryolamella (BY4741) | Defocus | 3 | 93,204 |
| Cryolamella (BY4741) | Volta | 5 | 129,473 |
| Cryolamella (LGY0016) | Defocus | 4 | 76,169 |
| Cryolamella (LGY0016) | Volta | 3 | 92,989 |
| Cryosections (LGY0016) | Volta | 4 | 33,194 |
| Cryolamella (LGY0007) | Volta | 4 | 166,582 |

**Table S8.** Nucleus volume sampled for subtomogram analysis.

| Tomogram | Volume (nm<sup>3</sup>) |
| --- | --- |
| 20211117_004 | 203,763,976 |
| 20211117_009 | 126,593,404 |
| 20211117_037 | 37,130,529 |
| 20211117_070 | 193,990,991 |
| 20211117_075 | 106,392,372 |
| Total | 667,871,272 |

